# Disentangling the Crabtree and Warburg effects across syngeneic non-tumor and tumor cells

**DOI:** 10.64898/2026.10.06.756891

**Authors:** Tomás Duraj, Eric Y. Miller, Linh Ta, Chloe Lee, Derek C. Lee, María Badiola Ramos, Juan J. Aristizabal-Henao, Michael Kiebish, Purna Mukherjee, Mar Boleda Pellicer, Caroline Kornreich, Oscar Shaver, Thomas N. Seyfried

## Abstract

Oxidative phosphorylation (OxPhos) and substrate-level phosphorylsation (SLP) are the primary biochemical mechanisms for ATP generation in mammalian cells. Increased dependency on SLP can arise from either transient (e.g., Crabtree effect) or chronic (e.g., Warburg effect) suppression of OxPhos. We evaluated the steady-state bioenergetics of quiescent and proliferating mouse cells matched in genetic background and cell biology, including primary thioglycollate-elicited peritoneal macrophages, malignant VM-M3 glioma with macrophage characteristics, and malignant CT-2A astrocytoma. Data are presented showing: 1) OxPhos insufficiency was the primary bioenergetic hallmark of proliferating malignant cells. 2) Dysregulated proliferation was associated with lower metabolic flexibility under nutrient stress than was growth arrest or regulated proliferation. 3) The Crabtree effect was present in macrophages and involved a glucose- and time-dependent suppression of OxPhos coupled to ATP production through SLP and resistance to mitochondrial inhibitors; lowering glucose concentration or substituting glucose with respiratory fuels increased OxPhos and restored sensitivity to mitochondrial inhibitors. 4) The Warburg effect in tumor cells involved a persistent SLP dependency and resistance to mitochondrial inhibitors under equivalent glucose-replete conditions. 5) OxPhos could sustain the regulated proliferation of non-tumor cells in the absence of a Warburg effect. Collectively, our findings emphasize that comparisons between normal and cancer cell metabolism require long-term experimental endpoints that ensure steady-state ATP sufficiency.

## Introduction

Otto Warburg proposed that “energy alone” was the central consideration for the origin and management of cancer [1]. The advent of molecular biology largely redirected research towards genetics and signal transduction, leaving many questions about what *energizes* cellular processes behind. Even as interest in cancer metabolism resurged, current models often operate under the assumption that “ATP availability is not limiting for cell growth” [2, 3], focusing instead on the anabolic demands of proliferation [3–5]. While it is conceivable that the maximal rate of cell division is limited by biosynthesis under nutrient-replete conditions, we postulate that “energy alone” is the fundamental determinant that separates life from death, driving thermodynamically unfavorable, endergonic processes and maintaining low-entropy states. In other words, while cell death can occur through a variety of mechanisms unrelated to energy [6], cells with sustained viability must continuously meet energy demands [7]. ATP sufficiency is the prerequisite that enables all essential cellular functions, including ion transport, macromolecular synthesis (e.g., RNA/DNA, protein and lipid synthesis), and ultimately cell division [8]. If the main goal of cancer therapy is to eradicate tumor cells, it becomes critical to understand not only the downstream processes enabled by energy (e.g., oncogenic signaling), but also the minimal requirements for ATP sufficiency. Differences in bioenergetic flexibility between malignant and normal cells may in turn reveal therapeutic vulnerabilities with direct application to the clinic [9, 10].

To define ATP sufficiency, it is important to differentiate between the metabolic substrate (e.g., glucose), the catabolic pathway (e.g., glycolysis), and the biophysical mechanisms that convert potential energy into usable energy (ATP synthesis). Substrate-level phosphorylation (SLP) and oxidative phosphorylation (OxPhos) are the primary mechanisms for ATP generation in eukaryotic mammalian cells. The Embden-Meyerhof-Parnas glycolytic pathway encompasses cytosolic SLP, producing ATP, NADH, and biosynthetic precursors. Beyond the cytosol, metabolites feeding into the tricarboxylic acid (TCA) cycle also contribute to mitochondrial SLP (mSLP), where succinyl-CoA ligase transfers inorganic phosphate to ADP (or GDP). Glutaminolysis in particular has been shown to drive high flux through mSLP in cancer cells [11, 12]. Conversely, Complex V produces ATP through the highly integrated, multi-step process of OxPhos, which requires generation of reducing equivalents via the TCA cycle or transfer from other cellular compartments (e.g., malate-aspartate or glycerol-3-phosphate shuttle), a functional electron transport chain (Complexes I through IV), structural integrity, and the maintenance of the proton motive force (ΔΨm). Because OxPhos is a complex multi-step process, defects in the structure, function, and number of the cellular chondriome may significantly reduce its efficiency and ultimately diminish ATP yield [13].

Total ATP content in living cells can be determined using ATP-linked reporters (e.g., firefly luciferin-luciferase), but the underlying ATP-generating mechanisms are largely estimated using surrogate markers such as extracellular lactate (cytosolic SLP via glycolysis, Crabtree/Warburg effect) and oxygen consumption rate (electron transport chain, OxPhos).

Analogous to lactate, extracellular succinate has been suggested as a marker of glutamine fermentation via mSLP (Q-effect) during acute (e.g., hypoxia) or chronic OxPhos insufficiency [12, 14–18]. All three proxy markers are used in this manuscript to estimate SLP and OxPhos.

While the entire intracellular ATP pool turns over in seconds [19], a transient drop in ATP is generally not accompanied by immediate cell death, but rather by a series of rapid and protracted adaptations aimed at increasing energy production and/or reducing demand [20–22]. By thermodynamic definition, only cells with sustained, long-term viability must have met ATP demands and thus achieved steady-state ATP sufficiency. It is estimated that a substantial fraction of baseline ATP hydrolysis (≈ 30 to 70%) is dedicated to fueling transmembrane ion pumps to preserve ion gradients [7, 23–25]. Proliferating cells must also meet the energy demand of biosynthesis [7, 8, 26]. Although the relative ATP cost of proliferation is debated, with some suggesting that the amount of ATP required for cell division may be less than that required for normal cell function (e.g., synthesis of proteins such as enzymes or hormones) [2, 27, 28], dysregulated growth must cover both baseline and biosynthetic ATP requirements [8, 26]. Thus, bioenergetic flexibility under nutrient stress involves dynamic adjustments to both ATP production and demand, seemingly at thermodynamic odds with the “uncontrolled proliferation” that defines cancer. However, extrapolations about steady-state ATP sufficiency based on short-term experimental endpoints, dying cells, or failure to account for the contribution of SLP/OxPhos are ubiquitous in the literature [29–33], which has generated significant confusion about the bioenergetic flexibility of cancer cells.

Evaluations of ATP sufficiency are further confounded by the Warburg/Crabtree effect. The Embden-Meyerhof-Parnas glycolytic pathway can provide ATP via cytosolic SLP coupled with complete substrate oxidation (OxPhos) or via cytosolic SLP with lactic acid fermentation (Warburg effect or Crabtree effect). The term “Warburg effect” was coined by Racker as “the presence of high aerobic glycolysis in tumors” [34] and is now recognized as a metabolic hallmark of cancer [35]. Importantly, the Warburg effect has been repeatedly demonstrated in tumors *in vivo* [36–40]. Several mechanistic explanations have been offered for this phenotype, including OxPhos insufficiency by Warburg himself [41, 42], faster ATP production by glycolysis [43, 44], generation of glycolytic intermediates [28], demand for oxidized NAD outpacing ATP turnover [45], uncoupling of biosynthesis from NADH generation [46], saturation of mitochondrial NADH shuttles [47], and other hypotheses such as mitigation of oxidative stress [48–51].

Contrary to Warburg’s energy-centric focus, the term “Warburg effect” is not defined by energy but by a juxtaposition of oxidation and fermentation, higher than expected glycolytic flux, and its end-product (lactate). Warburg frequently used “aerobic glycolysis” and “aerobic fermentation” interchangeably. The modifier “aerobic” establishes the physiological expectation that, in the presence of sufficient oxygen, OxPhos should be the predominant energy-generating mechanism in eukaryotic cells, thus yielding a state of “aerobic OxPhos”, wherein glycolysis proceeds without lactic acid production (a regulatory principle known as the Pasteur effect, the inhibition of fermentation by oxygen) [52]. In contrast, the Warburg effect is defined by persistent production of lactic acid despite oxygen tensions sufficient to support the full, terminal oxidation of glucose [53]. While energy is generated via cytosolic SLP in the glycolytic pathway regardless of oxygen, the defining step of “aerobic lactic acid fermentation” via lactate dehydrogenase yields no ATP, but rather cytosolic NAD⁺ required for continuous glycolytic flux [45]. Consequently, characterizing the Warburg effect requires accounting for both cytosolic ATP and redox balance.

Phenotypically equivalent, the Crabtree effect is often defined as reversible *glucose-induced* repression of respiration accompanied by lactic acid fermentation in the presence of oxygen [54]. The precise biochemical mechanisms driving the Crabtree effect remain a topic of debate. Historically, glucose-induced respiratory suppression was attributed to intracellular competition for free ADP and inorganic phosphate [55–58], a reduction in cytosolic pH [59], as well as glucose-induced Ca^2+^ uptake by the mitochondria, which inhibits ATP synthase [60]. Contemporary models also incorporate allosteric inhibition of respiration by glycolytic intermediates [61, 62]. In cell culture, the Crabtree effect is considered an artifact driven by adaptation to unphysiologically high glucose concentrations, although it may manifest under high glycolytic flux *in vivo* [62–65]. Technically, the Crabtree/Warburg effect can be observed with any glycolytic substrate capable of driving high glycolytic flux, such as fructose or uridine [66–70].

Experimentally, the Crabtree effect can be mitigated by reducing glucose concentrations or eliminating glucose from the culture medium altogether. However, given that total glucose deprivation is generally accompanied by cell death, most experimental approaches substitute glucose with galactose. Galactose is metabolized through the Leloir pathway and limits cytosolic ATP yield and glycolytic kinetic rate, diverting aldohexose carbons to the pentose phosphate pathway [71–73], thus forcing a higher reliance on OxPhos and mSLP [63]. Because the Crabtree and Warburg effects may coexist and elicit identical surrogate markers (high lactate, low oxygen consumption), we postulate that distinguishing them requires experimental approaches that uncouple oxygen consumption and redox balance from lactic acid fermentation over extended intervals.

Mitochondrial SLP refers to succinyl-CoA ligase catalyzed production of ATP (or GTP, rapidly interconverted to ATP via nucleoside-diphosphate kinases) in the mitochondrial matrix. In cancer cells, glutamine drives mSLP flux through conversion to glutamate, α-ketoglutarate, succinyl-CoA and ultimately succinate [12]. When OxPhos is impaired (e.g., hypoxia or electron transport blockade), or when flux through mSLP exceeds succinate oxidation, succinate may be exported outside the mitochondria as a “fermentation” byproduct (Q-effect), similar to lactate during aerobic fermentation [12, 13]. Continuous glutamine-driven mSLP flux requires intramitochondrial NAD^+^ regeneration via Complex I, or alternative processes to oxidize NADH, to support α-ketoglutarate dehydrogenase [11, 74–77]. While mSLP has been shown to provide sufficient matrix ATP to preserve mitochondrial membrane potential (ΔΨm) and prevent reverse operation of ATP synthase under OxPhos inhibition over short intervals and in isolated mitochondria [11, 14, 74, 78, 79], it is not known whether mSLP could sustain ATP sufficiency in the simultaneous absence of ATP from Complex V and cytosolic SLP.

Finally, OxPhos must be functionally differentiated between oxygen consumption for electron transport (“Ox”) and ATP output by Complex V (“Phos”), as respiration is not necessarily linked to ATP generation [80]. While it has been argued that some level of functionality of the “mitochondrial respiratory chain” is required for optimal proliferation, supporting redox balance and biosynthesis [81–85], this only refers to the “Ox” in “OxPhos”. In tumor cells, “Phos” may be dispensable as long as cytosolic and mitochondrial SLP are sufficient [86–90]. Furthermore, “Ox” is mechanistically entangled with mSLP through NAD^+^ regeneration. Therefore, while oxygen consumption is not a quantitative marker for the ATP yield of OxPhos, subject to many experimental caveats and assumptions [91], it can be directionally informative about the relative dependency on mitochondrial metabolism when accounting for other ATP sources.

To disentangle these phenomena, we evaluated the steady-state ATP sufficiency of biologically and genetically matched non-malignant and malignant cells. We compared primary non-tumor macrophages with malignant cells (VM-M3 and CT-2A), matched in aspects of cell biology, genetic background and species of origin. We hypothesized that dysregulated proliferation may be less bioenergetically flexible than regulated activity (including regulated proliferation), based on evolutionary principles that drive normal cells to respond effectively and efficiently to environmental cues such as nutrient stress. Applying long-term viability tracking, nutrient restriction, metabolomic profiling and metabolic assays, we show that non-tumor cells display the Crabtree effect during both quiescence and regulated proliferation. Conversely, malignant cells exhibit the Warburg effect during both dysregulated proliferation and forced growth arrest. By halting the cell cycle, baseline ATP sufficiency can be restored via glutaminolysis in the absence of lactic acid fermentation. These findings challenge the contemporary view that the Warburg effect is a universal feature of all proliferating cells. Instead, our findings suggest that the Warburg effect, and by extension chronic SLP dependency, may be specific to dysregulated proliferation. We propose that comparisons of bioenergetic flexibility between normal cells and cancer cells should be performed under conditions of steady-state ATP sufficiency, while controlling for confounding from the Crabtree/Warburg effect.

## Materials and Methods

### Mice

Mice (*Mus musculus*) of the C57BL/6J inbred strain were obtained from the Jackson Laboratory, Bar Harbor, Maine, USA. Mice from the VM/Dk inbred strain were originally obtained as gifts from G. Carlson (McLaughlin Research Institute, Great Falls, Montana, USA) and H. Fraser (University of Edinburgh, Scotland, UK). Additionally, C57BL/6J × VM/Dk F_1_ hybrid mice were bred in-house. All mice used in this study were housed and bred in the Boston College Animal Care Facility under IACUC-approved husbandry conditions as previously described [92]. Male and female mice between 8 and 10 weeks of age were used for all studies. C57BL/6J mice elicited a higher peritoneal macrophage yield upon thioglycollate stimulation than VM/Dk mice. Following the 3Rs ethical principles and after confirming equivalent metabolic profiles between strains (cf. Figure S1.1), C57BL/6J-derived macrophages were examined in most experiments, unless otherwise noted. Preliminary comparisons also revealed no differences in yield or viability between sexes (data not shown), as already reported in [93]; consequently, cells isolated from males and females were pooled. All animal procedures and protocols were in strict accordance with the NIH Guide for the Care and Use of Laboratory Animals and approved by the Institutional Animal Care Committee at Boston College under assurance number D16-00521.

### Thioglycollate-elicited peritoneal macrophages

The protocol for generating and collecting thioglycollate-elicited peritoneal macrophages was adapted from previously published methods [94, 95]. Briefly, 8-10-week-old mice were intraperitoneally injected with 1 mL of sterile 3% thioglycollate broth (Sigma-Aldrich, St. Louis, MO, USA). After 96 h, mice were euthanized by cervical dislocation in accordance with the approved IACUC protocol, and the peritoneal cavity was lavaged with 10 mL of glucose-, glutamine-, and serum-free Dulbecco’s Modified Eagle Medium (DMEM) (D5030; Sigma-Aldrich) supplemented with 3.7 g/L sodium bicarbonate (S5761, Sigma-Aldrich) and 1% penicillin-streptomycin (15140-122; Gibco, Waltham, MA, USA). The lavage fluid was centrifuged at 300 × g for 5 min, and the cell pellet was resuspended in seeding medium for 2 h in the same glucose-, glutamine-, and serum-free DMEM as before. After 2 h, non-adherent cells and red blood cells were removed by washing three times with Dulbecco’s Phosphate Buffered Saline (DPBS) with calcium and magnesium (SH30264.02, Cytiva, Marlborough, MA, USA). Cells were then maintained in experimental media for all subsequent analyses.

Thioglycollate-elicited peritoneal macrophages are mature, mitotically quiescent primary cells [96, 97]. We also note that thioglycollate-elicited macrophages are not fully committed to either M1 or M2 phenotypes [98]. We explicitly avoided LPS-mediated polarization, which increases glycolysis and intracellular itaconate production that sequesters Coenzyme A while inhibiting succinate oxidation, thus potentially impairing mitochondrial ATP generation [99–101]. For regulated proliferation assays, cell division in macrophages was elicited using recombinant mouse macrophage colony-stimulating factor (M-CSF) (AF-315-02, PeproTech, Rocky Hill, NJ, USA) [102]. Unless otherwise stated, M-CSF was used at a concentration of 100 ng/mL. Total protein was quantified using the Pierce BCA Protein Assay (23225, Thermo Scientific, Rockford, IL, USA) for normalization and to confirm proliferation. To generate functionally ρ^0^-like macrophages with suppressed respiratory capacity, thioglycollate-elicited peritoneal macrophages were exposed to low-dose ethidium bromide (50 ng/mL) for mtDNA depletion following previously described methods [103]. For this experiment, macrophages were maintained in a 96-well plate in complete DMEM-based medium (10-017-CV, Corning, NY, USA) with 10% fetal bovine serum (F-0500-DR, Atlas Biologicals, Fort Collins, CO, USA) and 1% penicillin/streptomycin (30-002-CI, Corning, NY, USA), supplemented with 100 µg/mL pyruvate (P5280, Sigma-Aldrich) and 50 µg/mL uridine (U3003, Sigma-Aldrich) for 8-10 weeks, with medium replacement once per week, and maintained alongside matched, untreated isogenic parallel controls (no ethidium bromide). After the indicated period, cells were washed three times with DPBS prior to exposure to experimental media.

### Cell lines

The VM-M3 cell line was established by our group from a spontaneously arising tumor in the cerebrum of an adult VM/Dk inbred mouse, as previously described [104, 105]. The highly invasive VM-M3 cells express multiple properties of macrophages/microglia including morphological appearance, surface adhesion, phagocytic behavior, total lipid composition (glycosphingolipids and phospholipids) and gene expression (positive for CD11b, Iba1, F4/80, CD68, CD45 and CXCR4; negative for nestin, GFAP, NF200 and CD19) [104, 105]. The CT-2A cell line was established by our group from a tumor induced via 20-methylcholanthrene in the cerebral cortex of a C57BL/6J mouse [106]. CT-2A is histologically classified as a poorly differentiated, highly malignant anaplastic astrocytoma (GFAP-positive), p53 wild-type, PTEN-deficient, recapitulating features of high-grade gliomas, including high mitotic index, nuclear polymorphism, hemorrhage, pseudopalisading necrosis, and microvascular proliferation when grown orthotopically *in vivo* [106–108]. CT-2A cells express stemness markers (Oct4 and Nestin, with CD133 positive populations) [109, 110]. Both models have been deposited in cell line collections (VM-M3: ABM T8016; CT-2A: Millipore SCC194). The v-raf/v-myc immortalized BV-2 mouse microglial cell line derived from C57BL/6 mice was obtained from Cytion (305156). All cell lines were transfected with a lentiviral vector containing the firefly luciferase gene under the control of the CMV promoter, resulting in the CT-2A/Fluc, VM-M3/Fluc and BV-2/Fluc cell lines [104, 111]. All cell lines were routinely tested and free of mycoplasma.

### Culture conditions and experimental media

CT-2A, VM-M3 and BV-2 cell lines were maintained in standard glucose- and glutamine-containing DMEM-based medium (10-017-CV, Corning) supplemented with 10% fetal bovine serum (F-0500-DR, Atlas Biologicals) and 1% penicillin/streptomycin (30-002-CI, Corning) in a humidified incubator at 37 °C and 5% CO_2_. Depending on the experimental design, cell lines were seeded in the medium described above at 4.0×10^4^ cells/well in 6-well plates, or 2.0×10^4^ cells/well in 24-well plates. After seeding, cells were allowed to adhere for 24 h. Cells were then washed twice with DPBS to remove any residual medium and incubated in the corresponding media for the duration specified in each protocol.

For metabolic analyses, experimental media were formulated from glucose- and glutamine-free basal DMEM (D5030; Sigma-Aldrich) or DPBS with calcium and magnesium (SH30264.02, Cytiva), both supplemented with 1% penicillin-streptomycin (30-002-CI, Corning). When indicated, experimental media were also supplemented with 10% dialyzed fetal bovine serum (SH3007903, Cytiva). To ensure trace glucose was not a confounder, glucose concentrations in 100% dialyzed serum were independently verified to be ≤ 5 mg/dL (as reported by the manufacturer) using a glucose oxidase assay (1070-125, Stanbio Laboratory, Boerne, TX, USA).

To evaluate the role of the cell cycle, DPBS was utilized to induce cell cycle arrest, or conversely, DPBS was reconstituted with the formulation of DMEM by addition of 8 vitamins and 14 amino acids. In experiments where DPBS was reconstituted to recapitulate the formulation of DMEM, the following amino acids were supplemented at the indicated final concentrations: Glycine (0.4 mM, 410225, Sigma-Aldrich), L-Arginine (0.4 mM, BP2505100, Fisher BioReagents), L-Cystine (0.2 mM, J63745.14, Thermo Scientific), L-Histidine (0.2 mM, H8125, Sigma-Aldrich), L-Isoleucine (0.8 mM, I2752, Sigma-Aldrich), L-Leucine (0.80 mM, J62824, Thermo Scientific), L-Lysine (0.8 mM, A16249, Thermo Scientific), L-Methionine (0.2 mM, M9625, Sigma-Aldrich), L-Phenylalanine (0.4 mM, A13238, Thermo Scientific), L-Serine (0.4 mM, J62187, Thermo Scientific), L-Threonine (0.8 mM, AC138930050, Thermo Scientific), L-Tryptophan (0.08 mM, J62508, Thermo Scientific), L-Tyrosine (0.4 mM, J63511, Thermo Scientific), and L-Valine (0.8 mM, J62943, Thermo Scientific). Vitamins were supplemented using the MEM Vitamin Solution 100X (11120-052, Gibco) at a 4X working concentration to achieve the following final concentrations (in mM): i-Inositol (0.04440), Nicotinamide (0.03275), Choline chloride (0.02865), Pyridoxal hydrochloride (0.01964), Thiamine hydrochloride (0.01186), Folic acid (0.00907), D-Calcium pantothenate (0.00839), and Riboflavin (0.00106).

Metabolites were supplemented at the specified concentrations: D-Glucose (12.0 mM or 5.0 mM, G7021, Sigma-Aldrich), D-Galactose (12.0 mM, G5388, Sigma-Aldrich), L-Glutamine (2.0 mM, 68540, Sigma-Aldrich), L-Lactate (12.0 mM, 71718, Sigma-Aldrich), non-racemic (R)-β-hydroxybutyrate (5.0 mM, HY-W015851, MedChemExpress), acetoacetate (5.0 mM, A8509, Sigma-Aldrich). Where indicated, media were supplemented with 500 μM exogenous oxidized NAD (NAD^+^, 481911, Millipore Sigma) or reduced NAD (NADH, J61638, Thermo Scientific). To evaluate the temporal stability of the Crabtree and Warburg effects, cells were cultured in standard “high glucose” (12.0 mM) and physiological “low glucose” (5.0 mM) environments, and their respirometry divergence was tracked continuously over 72 h. Pharmacological interventions included the following ETS inhibitors: rotenone (R8875, Sigma-Aldrich), sodium azide (S227I, Fisher Scientific), and oligomycin (495455, Sigma-Aldrich); the CPT1 inhibitor etomoxir (11969, Cayman Chemical); and the CDK4/6 inhibitor palbociclib (HY-50767A, MedChemExpress). Palbociclib pre-treatment was performed by exposing cells to 5 μM palbociclib in glucose- and glutamine-containing DMEM with 10% dialyzed serum for 120 h, washing cells three times with DPBS, and switching to the indicated media conditions.

### Luciferin-luciferase bioluminescence assay

Following the designated experimental time, luciferin (10 mg/mL, Syd Labs, Hopkinton, MA, USA) was added to each well containing luciferase-tagged cells using a repeater pipette and incubated at room temperature for 1 min, as previously described [104]. Bioluminescence was quantified using the Ami HT imaging device (Spectral Instruments, Tucson, AZ, USA) with a 300-second exposure. Photon values were obtained using Aura software (version 2.3.1) via ROI selection. In viable luciferase-tagged cells, luciferase catalyzes the reaction between luciferin and ATP, producing oxyluciferin and light, with luminescence intensity proportional to the ATP content in the sample; non-viable cells fail to maintain metabolic activity and ATP levels, and thus exhibit no bioluminescence [112]. When bioluminescence values are tracked over time, total ATP content can be used to estimate cell proliferation [14, 112].

### Calcein-AM and Ethidium Homodimer III viability assay

Ethidium homodimer III (EthD-III) and calcein-AM were used to analyze cell viability [113, 114]. EthD-III is a membrane-impermeant dye that emits red fluorescence when bound to nucleic acids. EthD-III only enters cells when the plasma membrane is compromised, thus marking dead or late apoptotic cells. Calcein-AM is a membrane-permeable dye that is hydrolyzed by intracellular esterases to produce green fluorescence. After the designated incubation period, a freshly prepared 50 µL DPBS solution containing 4 mM calcein-AM and 2 mM EthD-III (30002, Biotium, Fremont, CA, USA) was added to each well. Plates were incubated for 60 min in a cell culture incubator at 37 °C and 5% CO_2_. Fluorescence images were acquired using an EVOS Imaging System (Life Technologies, Carlsbad, CA, USA) with a 10× objective and Texas Red or GFP filter cubes. CellProfiler software (version 4.2.6, Broad Institute, Cambridge, MA, USA) was used to algorithmically quantify the live/dead ratio, as previously described [93, 115, 116].

### Cell cycle analysis via flow cytometry

Flow cytometry was utilized to determine the percentage of cells in G1, S, and G2/M phases. Cells cultured in DMEM, DPBS, or experimental media with palbociclib were harvested after the indicated incubation time, fixed with ice-cold ethanol and stained with propidium iodide to quantify total DNA content, as previously described [117]. Propidium iodide (P4170, Sigma-Aldrich) was used at a final concentration of 40 µg/mL and RNase A (BP2539, Fisher BioReagents) at 200 µg/mL. Data were acquired on a FACSAria IIIu Cell Sorter (BD Biosciences) and the cell cycle was analyzed using FlowJo 10.7.1 software (BD Life Sciences) by applying the Watson Pragmatic algorithm [118].

### Oxygen Consumption Rate (OCR) measurement

Oxygen consumption rates were quantified utilizing the Resipher system (Lucid Scientific, Atlanta, GA, USA), adapting established protocols for longitudinal *in vitro* respirometry [119]. Briefly, cells were seeded in a 96-well plate at a density of 2.0×10^4^ cells/well and switched to experimental media after overnight attachment (cell lines) or directly after the isolation procedure (macrophages). Empty wells were filled with sterile water to prevent evaporation artifacts. For calibration, the sensing lid of the Resipher was placed on top of the plate for 10 min prior to starting OCR measurements. The cells and sensing lid were maintained at 37 °C in a humidified incubator at 5% CO_2_ and 90% relative humidity. Data were processed and analyzed on the Lucid Lab web application interface.

### Western Blot protein expression analysis

Western blots were performed to determine the presence of the mitochondrial ketolysis enzymes 3-hydroxybutyrate dehydrogenase (BDH1) and 3-oxoacid CoA-transferase 1 (OXCT1), as previously described [92]. Briefly, thioglycollate-elicited peritoneal macrophages, VM-M3 and CT-2A cells were homogenized in ice-cold lysis buffer (9803; Cell Signaling Technology, Danvers, MA, USA). Lysates were subjected to one freeze-thaw cycle at -80°C, transferred to 1.5 mL Eppendorf tubes, mixed on a rocker for 30 min at 4 °C and then centrifuged at 16,000 × g at 4°C for 20 minutes. Supernatants were collected and protein concentrations estimated using the Pierce BCA Protein Assay. A total of 30 μg of protein from each lysate was denatured with NuPAGE LDS Sample Buffer (NP0007, Invitrogen, Waltham, MA, USA) and Sample Reducing Agent (NP0009, Invitrogen). Samples were resolved by SDS-PAGE on 4-12% Bis-Tris gels (NP0321BOX, Invitrogen). Proteins were transferred to a PVDF membrane (1704156, Bio-Rad Laboratories, Hercules, CA, USA) and blocked in 5% nonfat powdered milk in TBS with Tween 20 (pH 7.6) for 1 hour at room temperature. Membranes were probed with BDH1 (15417-1-AP, Proteintech, Rosemont, IL, USA) or OXCT1 (67836-1-Ig, Proteintech) primary antibodies overnight at 4 °C with gentle shaking. Blots were then incubated with the corresponding secondary antibody (7074 and 7076, Cell Signaling) for 1 hour at room temperature. β-Actin (MABT825, Sigma) was used as a loading control. Bands were visualized with enhanced chemiluminescence (32106, Thermo Scientific).

### Extracellular lactate and succinate assays

Extracellular lactate (L-Lactate Assay Kit I, Eton Bioscience, San Diego, CA, USA) and extracellular succinate (EnzyChrom Succinate Assay Kit, BioAssay Systems, Hayward, CA, USA) were measured using colorimetric assays following the manufacturers’ instructions. Aliquots of media were collected at the indicated time points and absorbance was read at 490 nm (lactate) and 570 nm (succinate) using a SpectraMax M5 microplate reader (Molecular Devices, San Jose, CA, USA) and compared against standard curves. Data were normalized to protein concentration and cell number.

### NAD^+^/NADH Quantification

Intracellular NAD, NADH, and their ratios were quantified using the Amplite Colorimetric NAD/NADH Ratio Assay Kit (AAT Bioquest, Pleasanton, CA, USA; Cat 15273). CT-2A cells were seeded at 5.0×10^5^ cells/well in 6-well plates and allowed to adhere for 24 h. Cells were then washed twice with DPBS and the medium was changed to the indicated conditions. After 6 h, cells were washed three times with DPBS to remove any residual medium, lysed by incubation with Lysis Buffer (Component G) at 37 °C for 15 minutes, and the resulting supernatant was collected for quantification following the manufacturer’s instructions. Absorbance was monitored at 460 nm using the SpectraMax M5 microplate reader. Absorbance values were quantified against a parallel NADH standard curve and normalized to total protein content.

### Semiquantitative Nontargeted Metabolomics by Ultra-High Performance Liquid Chromatography-Tandem Mass Spectrometry (UHPLC-MS/MS)

Cells were seeded at 1.0×10⁶ cells in 100 mm culture-treated dishes (130182, Thermo Scientific) using standard complete medium (10-017-CV, Corning) supplemented with 10% fetal bovine serum and 1% penicillin/streptomycin. After overnight attachment, the plate was washed three times with DPBS and the medium was replaced with the indicated conditions supplemented with 10% dialyzed fetal bovine serum. Cells and corresponding protein were collected after 48 h from their respective plates using ice-cold methanol (047192K2, Thermo Scientific) and cell lysis buffer (9803; Cell Signaling Technology). Metabolites were isolated using a solid-phase extraction (SPE) 96-well plate (Isolute PLD+, Biotage, Salem, NH, USA) and 700 µL of 8:1:1 acetonitrile:methanol:acetone +0.1% formic acid at -20 °C in the presence of known amounts of ^13^C_6_ phenylalanine (Cayman Chemical, Ann Arbor, MI, USA). After elution with a 96-well plate positive pressure manifold, samples were fully dried under nitrogen gas at 25 °C, followed by reconstitution in 100 µL of 1:1 methanol:water and storage at 4 °C until analysis.

UHPLC-MS/MS analyses were performed on an Agilent 1290 Infinity HPLC coupled to a Thermo Q-Exactive Plus mass spectrometer. A 33-minute binary multi-step gradient was used with A: 100% water and B: 100% acetonitrile, both with 0.1% formic acid. The column was a Waters Acquity UPLC HSS-T3, 150 mm × 2.1 mm × 1.8 µm. The flow rate was 250 µL/min, and the column compartment temperature was 35 °C. The gradient elution program was as follows: starting at 0% B from 0 – 5 min, linear increase to 20% B from 5 – 15 min, followed by linear increase to 100% from 15 – 20 min, hold at 100% B from 20 – 25 min, drop to 0% B at 25.1 min and equilibration until the 33 min mark. Samples were analyzed using a polarity-switching full-scan mass spectrometry method at 17,500 resolution with spray voltages +3.5kV and -2.5kV in the positive and negative ion modes, respectively, with scan range *m/z* 60 -900. Centroided spectra were automatically lock-mass corrected during acquisition (leucine enkephalin infusion post-column using an isocratic pump). Representative pooled samples were used as targets in pseudo top-40 data-dependent tandem mass spectrometry experiments with iterative exclusion lists using IE-Omics [120]. Metabolites were putatively identified and peak areas were integrated using CompoundDiscoverer software (v3.3, Thermo Fisher Scientific, Waltham, MA, USA) as described earlier [121]. Raw peak areas for all analytes were normalized using the internal standard and total protein content of each sample as determined using the Pierce BCA Protein Assay. Data were analyzed using the MetaboAnalyst platform [122]. For multivariate analysis, data were normalized using median normalization, log10-transformed for variance stabilization, and scaled via Pareto scaling prior to principal component analysis (PCA). For heatmap visualization, data were preprocessed using median normalization, log10-transformed, and scaled via Auto scaling. Differential features were evaluated using linear models adjusted for covariates (cell media) with FDR-adjusted p < 0.05 [123], and the adjusted data matrix was used to construct the hierarchical clustering heatmap. The complete metabolomics dataset is available in Supplementary Table S1.

### Live-cell mitochondrial morphological profiling

To evaluate the basal and adaptive changes in the mitochondrial network, cells were cultured under viable experimental conditions for 120 h. Following incubation, cells were stained with MitoView™ 650 (70075, Biotium, Fremont, CA, USA), a mitochondrial stain for live/intact cells. Fluorescence images were acquired on a Leica Stellaris 5 microscope and 3D volumetric renderings were subsequently used to algorithmically quantify the total mitochondrial footprint (3D volume) and mitochondrial branch length using the Mitochondrial Network Analysis (MiNA) toolset [124].

### Statistics

Statistical analyses and data plotting were performed using GraphPad Prism 10.1.0 (GraphPad, La Jolla, CA, USA). Each experiment was performed with a minimum of 3 independent biological replicates. Individual biological replicates are plotted as distinct data points on the graphs, with sample sizes ranging from n = 3 to n = 14, as indicated. Data were analyzed by one-way or two-way ANOVA with Tukey, Šidák, and Dunnett multiple comparison post hoc analyses, or unpaired Student’s t-tests, as appropriate. A p-value ≤ 0.05 was considered statistically significant. Asterisks indicate the following: *p ≤ 0.05; **p ≤ 0.01; ***p ≤ 0.001; and ****p ≤ 0.0001.

## Results

### Long-term experimental endpoints are required to define steady-state ATP sufficiency in matched non-tumor and tumor cells

To elucidate the metabolic requirements of steady-state ATP sufficiency under matching aspects of cell biology and genetic background, we compared primary, quiescent, non-tumor thioglycollate-elicited peritoneal macrophages (myeloid origin) [125, 126] with the malignant VM-M3 cell line, a highly invasive glioma with macrophage characteristics (myeloid origin) [104, 105], as well as the CT-2A cell line, a high-grade astrocytoma with stem cell markers (neuroectodermal origin), differing in cell biology but sharing the C57BL/6J genetic background [106, 108, 110]. Importantly, comparisons were restricted to mouse host species to prevent inter-species discrepancies in basal metabolic rates [127].

To avoid capturing transient adaptations in cells undergoing or destined for cell death, we defined “ATP sufficiency” strictly through the lens of sustained, long-term cellular viability (Figure 1A). While cell death can occur through multiple mechanisms despite sufficient ATP, a cell that maintains long-term viability must, by thermodynamic definition, continuously meet its ATP demands [19].

**Figure 1.**
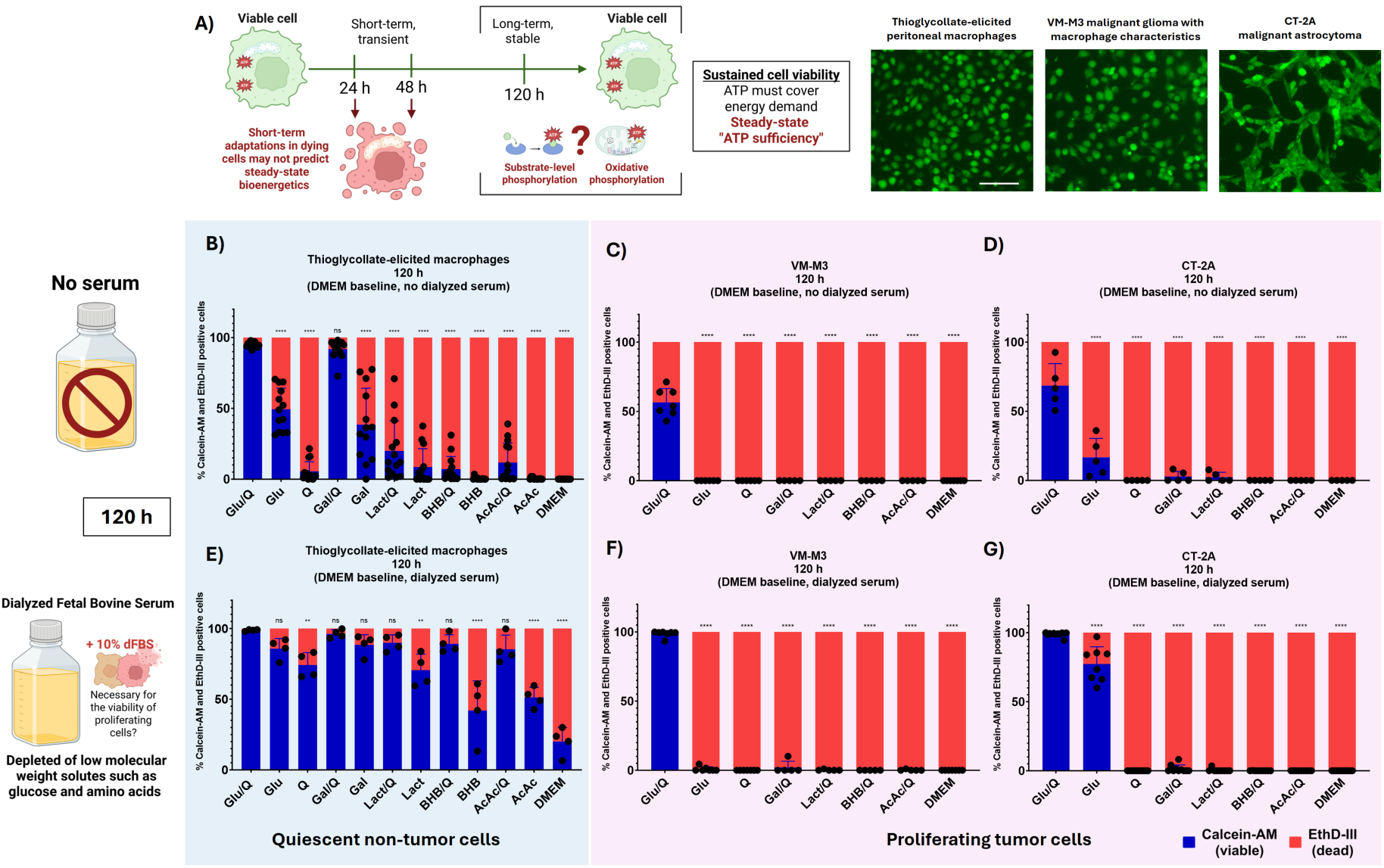
Long-term viability endpoints are required to define steady-state ATP sufficiency in non-tumor macrophages and malignant glioma cells. **(A)** Schematic of the experimental design to determine steady-state ATP sufficiency in primary macrophages compared with the VM-M3 glioma with macrophage characteristics, matched in cell biology (myeloid origin), and the CT-2A astrocytoma, differing in cell biology but matching the C57BL/6J genetic background. Scale bar: 100 μm. **(B–D)** Cell viability was assessed using Calcein-AM and EthD-III staining after 120 h (5 days) under various nutrient conditions in the absence of serum. **(B)** Primary non-proliferating macrophages exhibited high viability in glutamine combined with glucose or galactose, whereas **(C)** VM-M3 and **(D)** CT-2A cells survived only in the presence of glucose and glutamine. Single metabolites are omitted due to cell death at earlier time points (cf. Figure S1.1). **(E–G)** Cell viability in media supplemented with 10% dialyzed serum. **(E)** Macrophages sustained viability across multiple conditions, particularly those containing glutamine. Conversely, **(F)** VM-M3 and **(G**) CT-2A cells only maintained complete viability in the combination of glucose and glutamine. Data represent mean ± SD of at least 3 independent biological replicates. Statistical significance was determined using a one-way ANOVA followed by Dunnett’s multiple comparisons test (* p ≤ 0.05, ** p ≤ 0.01, *** p ≤ 0.001, **** p ≤ 0.0001). Abbreviations: Glu (Glucose), Gal (Galactose), Q (Glutamine), Lact (Lactate), BHB (β-hydroxybutyrate), AcAc (Acetoacetate), DMEM (non-supplemented, basal Dulbecco’s Modified Eagle Medium).

Consequently, we defined steady-state ATP sufficiency by focusing our primary analysis on long-term experimental endpoints of 120 h (5 days). Under serum deprivation, quiescent thioglycollate-elicited C57BL/6J macrophages required glutamine paired with metabolites capable of supporting glycolytic flux (glucose or galactose) to preserve unimpaired viability (Figure 1B). Although other metabolites conferred higher metabolic flexibility in macrophages compared with tumor cells, such combinations were unsuitable for assessing steady-state ATP sufficiency, as defined above, due to progressively declining viability. To confirm that this metabolic profile was not unique to the C57BL/6J genetic background and to validate our matching with the VM-M3 cell line (derived from VM/Dk mice), we additionally evaluated macrophages from VM/Dk and C57BL/6J × VM/Dk F_1_ hybrids (Figure S1.1, A-B). Macrophages from these genetic backgrounds exhibited comparable 120-h viability profiles, suggesting shared metabolic plasticity across the tested strains.

In contrast, VM-M3 (Figure 1C) and CT-2A (Figure 1D) cancer cells exhibited complete loss of viability at 120 h in all conditions under serum deprivation except the combination of glucose and glutamine. We hypothesized that this could be due to a requirement for serum components in proliferating cells [128, 129]. When supplemented with dialyzed fetal bovine serum (depleted of glucose and amino acids), quiescent macrophages demonstrated broad, sustained viability, particularly in metabolite combinations containing glutamine (Figure 1E). Yet, even when supplemented with dialyzed serum, VM-M3 (Figure 1F) and CT-2A cells (Figure 1G) remained viable only in media with glucose and glutamine. To ensure that trace glucose from dialyzed serum did not confound targeted nutrient deprivation, we confirmed glucose depletion via a glucose oxidase assay (Figure S1.2).

The need for protracted experimental endpoints is apparent when comparing 120-h results to short-term cultures (24 h and 48 h). We evaluated early viability under the same nutrient conditions, both in the absence (Figure S1.3, A-F) and presence (Figure S1.3, G-J) of dialyzed serum. In serum-free conditions, macrophages retained high viability across a broad spectrum of metabolite combinations (Figure S1.3, A, D), highlighting increased metabolic flexibility compared with VM-M3 and CT-2A tumor cells, which were highly sensitive to nutrient stress and serum deprivation (Figure S1.3, B-F). The addition of dialyzed serum partially improved the short-term viability of VM-M3 and CT-2A cells at 24 h (Figure S1.3, G, H). At 48 h, even in the presence of dialyzed serum, a gradual decline in viability was noted across most nutrient conditions, particularly single metabolites (Figure S1.3, I, J). However, despite a downward trajectory, residual viability was observed in combinations of glutamine with other oxidative metabolites (galactose, lactate, β-hydroxybutyrate and acetoacetate). Such transient adaptations could be confused with steady-state ATP sufficiency, as cells may temporarily survive on incomplete metabolic adaptations or internal stores without meeting long-term demands.

Taken together, our data indicate that while differences in metabolic flexibility can be observed early on, sustained viability and thus steady-state ATP sufficiency are only discernible at extended, cell-specific time points (≥ 120 h in the tested mouse cells). Thus, long-term experimental endpoints are necessary to circumvent confounding from transient but ultimately unsustainable metabolic adaptations.

### Restricting the proliferative capacity of tumor cells restores viability under galactose and glutamine culturing

There are three experimental approaches to circumvent the Crabtree/Warburg effect: replacing glucose with galactose, total glucose deprivation, and relative glucose deprivation. Galactose culturing reduces the cytosolic SLP rate while preserving glycolytic intermediates and the pentose phosphate pathway, thus promoting oxidative metabolism and glutaminolysis [63, 71, 130, 131]. We noted that non-proliferating macrophages maintained long-term viability in galactose and glutamine, while actively proliferating VM-M3 and CT-2A tumor cells did not. To determine whether cell death under galactose and glutamine culturing was due to an inability to meet the metabolic demands of proliferation, we evaluated the long-term viability of tumor cells under forced growth arrest.

Dulbecco’s Modified Eagle Medium (DMEM) is a standard culturing formulation enriched with amino acids and vitamins required for cell cycle progression and proliferation. Conversely, Dulbecco’s Phosphate Buffered Saline (DPBS), a balanced salt solution with a similar inorganic composition but lacking all amino acids and vitamins (see Methods), is typically used for short-term washing instead of long-term maintenance (Figure 2A). As expected, culturing VM-M3 and CT-2A cancer cells in DMEM supplemented with dialyzed serum drives robust proliferation over 120 h. In equivalent concentrations of glucose, glutamine, and dialyzed serum, DPBS-based medium preserved cell viability but completely abrogated proliferative capacity (Figure S2.1, A-B). Flow cytometry analysis of CT-2A cells revealed that baseline glucose and glutamine DMEM with dialyzed serum promoted high S-phase populations indicative of active proliferation. In contrast, DPBS culturing for 120 h caused a significant accumulation in the G1 phase with a reciprocal decrease in the S phase, even in the presence of glucose, glutamine and dialyzed serum (Figure 2B).

**Figure 2.**
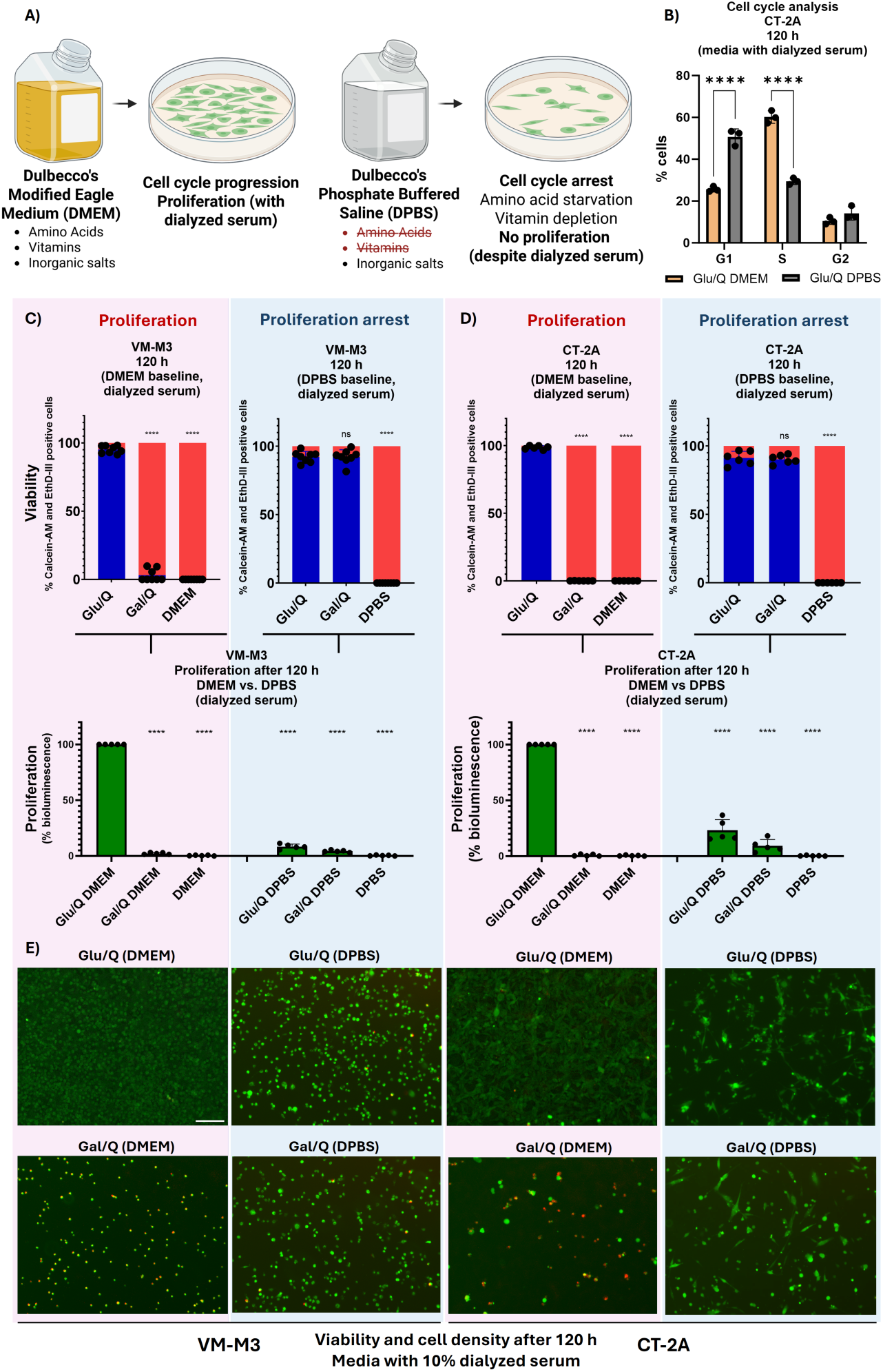
Proliferation arrest restores the long-term viability of VM-M3 and CT-2A tumor cells under galactose and glutamine culturing. **(A)** Experimental design. Dulbecco’s Modified Eagle Medium (DMEM) is enriched with amino acids and vitamins required for proliferation. Dulbecco’s Phosphate Buffered Saline (DPBS) contains only inorganic salts for osmotic balance. **(B)** Cell cycle analysis of CT-2A cells. DNA content was analyzed using propidium iodide (PI) flow cytometry after 120 h. Baseline DMEM exhibits high S-phase populations (DNA synthesis), whereas DPBS culturing shows a significant accumulation in the G1 phase with a reciprocal decrease in S phase. **(C)** Viability and proliferative capacity of VM-M3 cells cultured for 120 h in DMEM versus DPBS. Long-term DPBS culturing is typically avoided due to lack of amino acids and vitamins. Paradoxically, long-term viability was fully restored in galactose and glutamine when formulated in DPBS. **(D)** Viability and proliferative capacity of CT-2A cells cultured for 120 h in DMEM versus DPBS. **(E)** Representative microscopy images illustrating viability and cell density after 120 h in the indicated media conditions. Calcein-AM (green) and EthD-III (red). Non-viable cells are double-positive. Scale bar: 150 μm. Data represent mean ± SD of at least 3 independent biological replicates. Statistical significance was determined using a one-way ANOVA followed by Dunnett’s multiple comparisons test (* p ≤ 0.05, ** p ≤ 0.01, *** p ≤ 0.001, **** p ≤ 0.0001). Abbreviations: Glu (Glucose), Gal (Galactose), Q (Glutamine), DMEM (Dulbecco’s Modified Eagle Medium), DPBS (Dulbecco’s Phosphate Buffered Saline).

Paradoxically, formulating galactose and glutamine in DPBS rather than DMEM fully restored the 120-h viability of both VM-M3 and CT-2A cells (Figure 2, C, D). Proliferation (measured via total bioluminescence) and cell density on fluorescence microscopy (Figure 2E) were significantly reduced in DPBS compared with the growth-enabling DMEM. While dialyzed serum improved the overall robustness of VM-M3 and CT-2A cells, it was not strictly required for the rescue of long-term viability under galactose and glutamine culturing in DPBS (Figure S2.2, A, B).

To more precisely attribute the bioenergetic failure in galactose culturing to the pro-proliferative components of DMEM, we reconstituted the exact formulation of DMEM by supplementing DPBS with the corresponding concentrations of 8 vitamins and/or 14 amino acids (Figure S2.3A). In glucose and glutamine, the addition of amino acids to DPBS significantly improved proliferation, while the combined addition of vitamins and amino acids was necessary to fully mimic the growth rate of DMEM (Figure S2.3, B, C). Strikingly, instead of promoting viability by replenishing essential nutrients and cofactors, restoring proliferative capacity by replenishing vitamins and amino acids caused cell death in both cell lines when cultured in galactose and glutamine (Figure S2.3, B-D).

To further disentangle the mechanical demands of proliferation from broad nutrient deprivation signaling, we treated CT-2A cells with palbociclib, a clinical-stage CDK4/6 inhibitor, forcing cell cycle arrest despite pro-proliferative DMEM. A 120-h dose-response titration identified 5 μM palbociclib as an optimal concentration to halt proliferation without compromising cell viability, whereas 10 μM induced significant toxicity (Figure S2.4, A, B). At 5 μM, palbociclib successfully induced cell cycle arrest, characterized by a reduction in the S phase and an increase in the G2/M phase (Figure S2.3, E, F). Consistent with aberrant cell cycle dynamics, palbociclib-treated cells exhibited a markedly enlarged morphology alongside accumulation in the G2 phase. Crucially, pretreatment and continued exposure to palbociclib improved the viability and total bioluminescence of CT-2A cells cultured in galactose and glutamine formulated in DMEM to levels approaching DPBS (Figure S2.3, G).

Finally, to support the correlation with proliferation, we also tested the nutrient response of the v-raf/v-myc-immortalized but non-tumorigenic BV-2 microglia, originally isolated from C57BL/6 mice, matching macrophages and VM-M3 in cell biology (myeloid origin). Contrary to malignant VM-M3 and CT-2A cells, non-tumorigenic BV-2 cells sustained viability under galactose and glutamine culturing by undergoing spontaneous growth arrest despite pro-proliferative DMEM with dialyzed serum (Figure S2.5, A-C).

Taken together, our data suggest that uncoupling viability from the bioenergetic and biosynthetic demands of cell proliferation renders galactose and glutamine metabolism sufficient for the sustained survival of “non-proliferating tumor cells”.

### Non-proliferating normal cells sustain steady-state ATP sufficiency via cytosolic SLP (Crabtree effect) or OxPhos, whereas OxPhos fails to drive dysregulated tumor cell proliferation (Warburg effect)

Thus far, we have only examined the input metabolites enabling long-term viability, not the mechanism of ATP generation itself (SLP vs. OxPhos). To define the biochemical mechanisms enabling steady-state ATP sufficiency, we first analyzed the oxygen consumption rate (OCR) and extracellular lactate in thioglycollate-elicited macrophages. At both 24 h and 120 h, macrophages cultured in glucose and glutamine exhibited a significantly lower OCR compared with those cultured in galactose and glutamine (Figure 3A). This glucose-induced suppression of oxygen consumption (Crabtree effect) was accompanied by substantial extracellular lactate accumulation over 24 h, whereas macrophages cultured in galactose and glutamine produced minimal lactate (Figure 3B). To determine whether this high glycolytic flux was sufficient to meet basal ATP demands without OxPhos, we treated macrophages with electron transport system (ETS) inhibitors targeting Complex I (rotenone), Complex IV (azide), and Complex V (oligomycin). Macrophages were resistant to mitochondrial inhibitors when cultured in glucose, confirming that cytosolic SLP could preserve steady-state ATP sufficiency under the Crabtree effect. Conversely, macrophages cultured in galactose experienced complete loss of viability upon ETS inhibition (Figure 3C).

**Figure 3.**
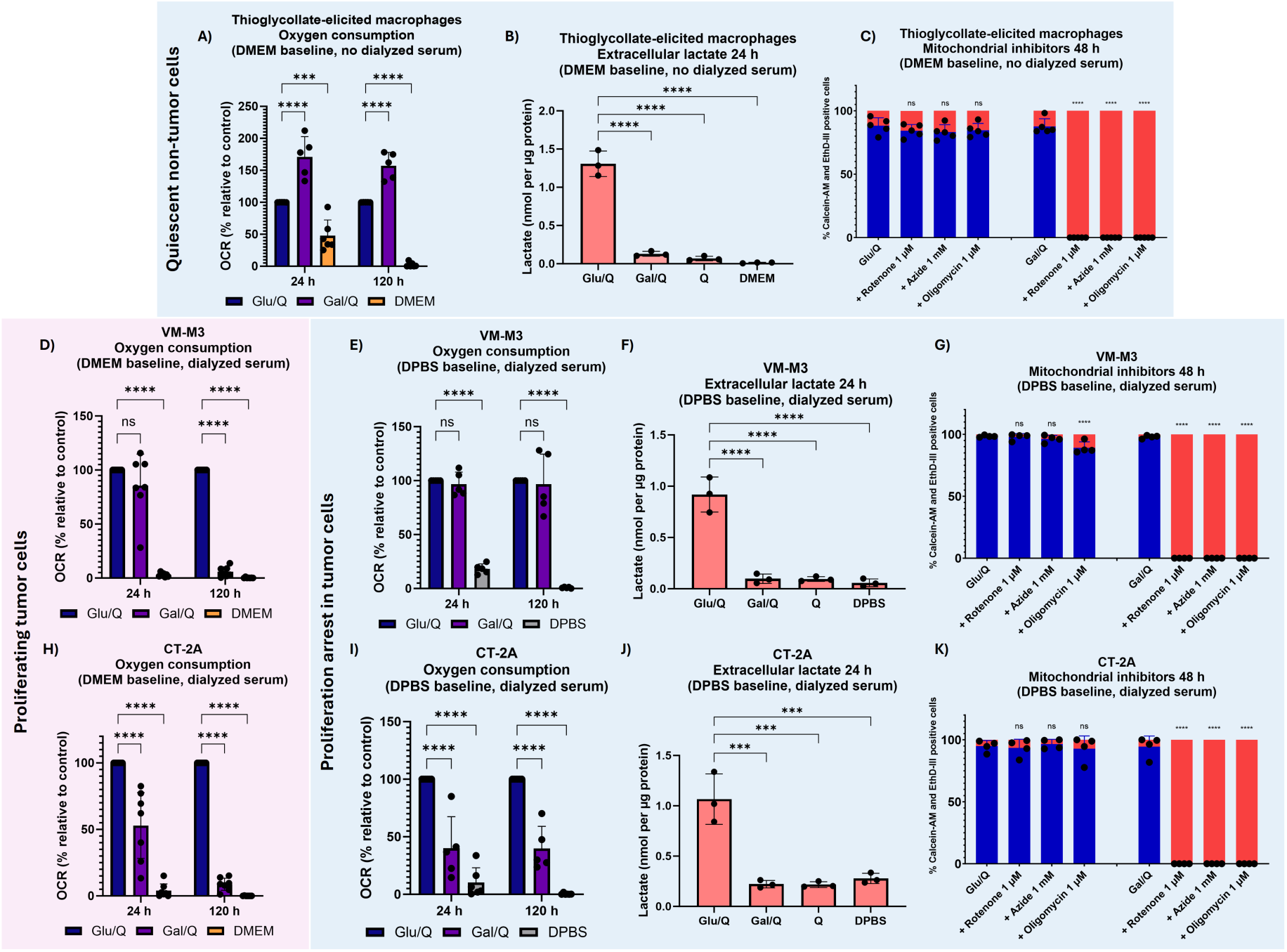
Non-proliferating macrophages sustain steady-state ATP sufficiency by cytosolic SLP with lactic acid fermentation when cultured in glucose (Crabtree effect) or OxPhos without lactic acid fermentation when cultured in galactose; conversely, malignant cells cannot sustain dysregulated proliferation using OxPhos alone. (A–C) Metabolic assessment of thioglycollate-elicited macrophages. **(A)** Oxygen consumption rate (OCR) at 24 h and 120 h. Glucose and glutamine culturing exhibited lower OCR than galactose and glutamine (glucose-induced suppression of OCR, i.e., Crabtree effect). **(B)** Extracellular accumulation of lactate over 24 h. Galactose and glutamine exhibited reduced lactate production. **(C)** Cell viability following treatment with the mitochondrial inhibitors rotenone (Complex I), azide (Complex IV), and oligomycin (Complex V). Macrophages were resistant to mitochondrial inhibitors in glucose, suggesting SLP was sufficient to cover ATP demands. In galactose, mitochondrial inhibitors induced complete loss of viability across all ETS complexes. **(D–G)** Metabolic assessment of VM-M3 cells. **(D)** OCR of proliferating VM-M3 cells at 24 h and 120 h (DMEM). Proliferating VM-M3 cells in galactose exhibited residual OCR at 24 h, but minimal OCR at 120 h. **(E)** OCR of non-proliferating VM-M3 cells at 24 h and 120 h (DPBS). In non-proliferating VM-M3 cells, OCR was maintained in galactose and glutamine at 120 h. **(F)** Extracellular accumulation of lactate over 24 h. In growth-arrested cells, galactose and glutamine preserved ATP sufficiency in the absence of lactic acid fermentation (Warburg effect). **(G)** Cell viability under DPBS culturing following treatment with mitochondrial inhibitors. Growth-arrested VM-M3 cells demonstrated high sensitivity to mitochondrial inhibitors under galactose culturing. **(H–K)** Metabolic assessment of CT-2A cells. **(H)** OCR of proliferating CT-2A cells (DMEM), **(I)** OCR of non-proliferating CT-2A cells (DPBS), **(J)** extracellular accumulation of lactate and **(K)** cell viability in DPBS following treatment with mitochondrial inhibitors. Data represent mean ± SD of at least 3 independent biological replicates. Statistical significance was determined using a two-way ANOVA with Šidák’s correction (A, D, H) and one-way ANOVA with Dunnett’s correction (B, C, E, F, G, I, J, K) (* p ≤ 0.05, ** p ≤ 0.01, *** p ≤ 0.001, **** p ≤ 0.0001). Abbreviations: Glu (Glucose), Gal (Galactose), Q (Glutamine), DMEM (Dulbecco’s Modified Eagle Medium), DPBS (Dulbecco’s Phosphate Buffered Saline).

To further confirm that this reliance on OxPhos was not a transient artifact of cell culture, we continuously maintained macrophages *in vitro* for 8-10 weeks (Figure S3.1A). Macrophages were left undisturbed or treated with low-dose ethidium bromide to induce mitochondrial DNA depletion. Ethidium bromide treatment significantly reduced basal OCR without affecting viability in glucose and glutamine (Figure S3.1B), while eliciting complete cell death in all oxidative nutrient conditions (galactose and glutamine) (Figure S3.1C). Isogenic control macrophages maintained high viability and transitioned to OxPhos, indicating that primary macrophages retain robust OxPhos capacity after extended cell culture maintenance.

Next, we examined whether the malignant VM-M3 and CT-2A cells mirrored the same metabolic profile. In short-term experimental endpoints, proliferating tumor cells cultured in DMEM were seemingly reliant on OxPhos when glucose was substituted with galactose, as evidenced by 24-h OCR profiles (Figure 3, D, H) and sensitivity to rotenone, azide, and oligomycin (Figure S3.2, A, B). However, despite transient adaptation, extended 120-h OCR tracking revealed a progressive decline in OCR. Both tumor cell lines produced robust amounts of lactate under glucose culturing over 24 h, regardless of dialyzed serum, indicative of prototypical Warburg phenotypes (Figure S3.3, A-D). This suggests that while the ETS may remain active in these cancer cells, OxPhos may be insufficient to support the bioenergetic, biosynthetic or redox demands of proliferation.

Conversely, growth-arrested VM-M3 and CT-2A cells maintained intermediate levels of OCR after 120 h under DPBS galactose culturing (Figure 3, E, I). VM-M3 and CT-2A cells cultured in glucose produced lactate, but those restricted to galactose and glutamine produced minimal lactate (Figure 3, F, J). Under proliferation arrest, both cell lines exhibited high sensitivity to mitochondrial inhibitors when cultured in galactose (Figure 3, G, K). Consequently, the metabolic profile of non-proliferating tumor cells more closely resembled that of quiescent primary cells, upholding steady-state ATP sufficiency without lactic acid fermentation. However, it is important to note that both galactose and glutamine were required for survival. Galactose may provide residual cytosolic ATP despite slower glycolytic flux, while glutaminolysis provides mitochondrial ATP via both OxPhos and mSLP.

### Supplementation with oxidized NAD circumvents the Crabtree/Warburg effect, restoring ATP sufficiency via glutaminolysis

Total glucose deprivation was accompanied by progressive cell death, and consequently not suitable for evaluation of steady-state ATP sufficiency. Beyond providing ATP, a non-bioenergetic consequence of lactic acid fermentation (Crabtree/Warburg effect) is the regeneration of cytosolic oxidized nicotinamide adenine dinucleotide (NAD^+^) via lactate dehydrogenase, which is required for upstream glycolytic flux. To determine whether glutaminolysis alone would be sufficient to cover ATP demands when uncoupled from cytosolic redox balance, we bypassed glycolysis entirely by providing oxidized NAD in the complete absence of glucose or galactose (Figure 4A). Culturing in glutamine as the sole carbon source ultimately results in complete loss of viability. Strikingly, direct medium supplementation with exogenous oxidized NAD fully restored the 120-h viability of macrophages cultured in glutamine alone (Figure 4B). This rescue was uniquely dependent on glutamine, as NAD^+^ supplementation failed to improve the viability of macrophages cultured in other single metabolites (glucose, galactose, lactate, β-hydroxybutyrate, or acetoacetate) (Figure S4.1). While not required, the addition of dialyzed fetal bovine serum further improved the long-term viability of macrophages in glutamine with NAD^+^ (Figure S4.2A).

**Figure 4.**
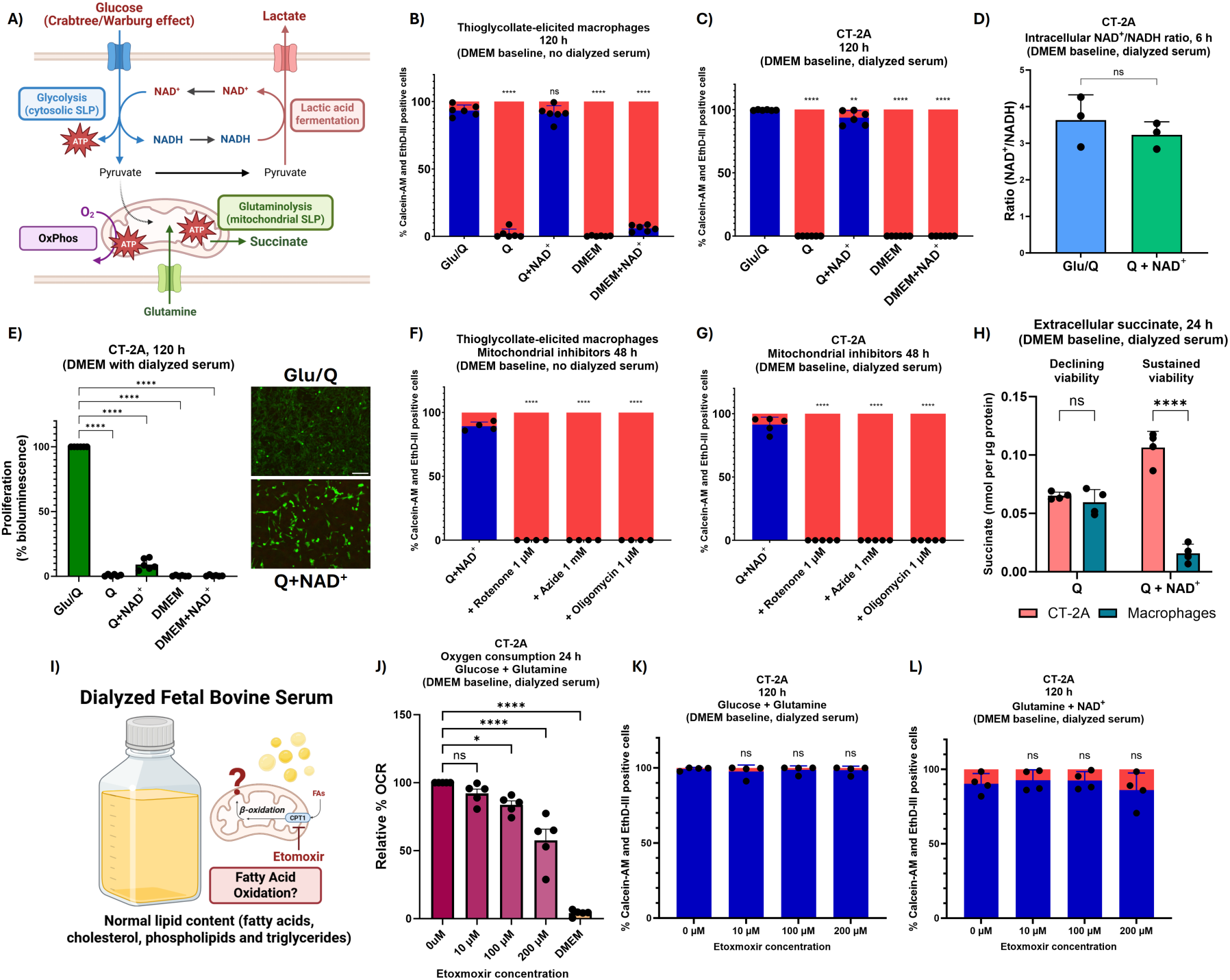
Supplementation of oxidized NAD (NAD^+^) rescues glutaminolysis-driven ATP sufficiency in macrophages and CT-2A tumor cells, circumventing the Crabtree/Warburg effect. **(A)** Illustration of NAD^+^/NADH balance. NAD**^+^** is consumed in the glycolytic Embden-Meyerhof-Parnas pathway and can be regenerated via lactic acid fermentation. To determine whether glutaminolysis could cover bioenergetic demands, ATP synthesis must be decoupled from cytosolic redox balance. **(B)** 120-h viability of macrophages and **(C)** CT-2A cells supplemented with NAD**^+^** (500 μM). NAD**^+^** restored long-term viability in glutamine alone. **(D)** NAD**^+^**/NADH ratio from CT-2A whole-cell extracts. **(E)** Proliferation rate of CT-2A cells in glutamine with NAD**^+^**. CT-2A cells did not proliferate in the absence of glucose, despite DMEM and dialyzed serum. Representative microscopy images with Calcein-AM and EthD-III staining. Scale bar: 150 µm. **(F)** Macrophage and **(G)** CT-2A viability under mitochondrial inhibition. Both cell types were sensitive to mitochondrial inhibitors when cultured in glutamine with NAD^+^. **(H)** Extracellular succinate after 24 h. CT-2A cells exported more succinate than macrophages when cultured in glutamine and NAD^+^. **(I)** Dialyzed serum is depleted of glucose and amino acids but retains normal lipid content. To determine whether fatty acid oxidation (FAO) was contributing to CT-2A viability, cells were treated with the CPT1 inhibitor etomoxir. **(J)** High-dose etomoxir (200 μM) significantly reduced OCR in CT-2A cells cultured in glucose and glutamine with dialyzed serum. **(K)** 120-h etomoxir treatment had no effect on the viability of CT-2A cells cultured in glucose and glutamine. **(L)** Similarly, etomoxir did not compromise the viability of CT-2A in glutamine with NAD**^+^** even at concentrations that reduced OCR, suggesting that FAO was dispensable under these conditions. Data represent mean ± SD of at least 3 independent biological replicates. One-way ANOVA with Dunnett’s correction (B-C, E-G, J-L), two-tailed paired Student’s t-test (D) and two-way ANOVA with Šidák’s correction (H) (* p ≤ 0.05, ** p ≤ 0.01, *** p ≤ 0.001, **** p ≤ 0.0001). Abbreviations: Glu (Glucose), Q (Glutamine), DMEM (Dulbecco’s Modified Eagle Medium).

This metabolic rescue was also observed in CT-2A cells. In the complete absence of glycolytic precursors, supplementation with NAD^+^ successfully restored steady-state ATP sufficiency in CT-2A cells cultured in glutamine alone (Figure 4C). Supplementation with exogenous NAD^+^ stabilized the NAD^+^/NADH ratio (Figure 4D), preventing the rapid depletion of the NAD^+^/NADH pool observed in glutamine alone (Figure S4.2B). CT-2A cells maintained high viability but ceased proliferation despite amino acids and vitamins from DMEM as well as growth factors and lipids from dialyzed serum (Figure 4E). Importantly, this rescue was specific to the oxidized state of NAD: whereas exogenous NAD^+^ restored viability in both DMEM and DPBS, reduced NAD (NADH) failed to do so, and even exacerbated cell death under glucose and glutamine culturing in DPBS, suggesting a narrow redox buffering capacity in CT-2A cells (Figure S4.3, A, B). Conversely, macrophages could utilize both oxidized and reduced NAD to support their viability in glutamine alone (Figure S4.3, C). Lastly, the VM-M3 cell line could not be rescued by NAD^+^ in glutamine alone, regardless of serum supplementation (Figure S4.2, C, D).

While exogenous NAD^+^ may alleviate cytosolic redox imbalance, glutamine utilization for both OxPhos and mSLP requires intramitochondrial NAD^+^ regeneration, which relies on forward-operating ETS to provide NAD^+^ for α-ketoglutarate dehydrogenase [74, 75]. To confirm whether glutaminolysis could be uncoupled from the ETS, we treated both cell types with mitochondrial inhibitors under glutamine and NAD^+^. Both macrophages and CT-2A cells (Figure 4, F, G) experienced complete cell death upon Complex I, IV or V inhibition, suggesting that viability under these conditions was also dependent on electron transport. This response was conserved in CT-2A cells cultured in DPBS medium (Figure S4.2E).

Succinate has diverse intracellular fates, but inefficient OxPhos (e.g., hypoxia or mitochondrial dysfunction) and increased mSLP can be accompanied by extracellular succinate accumulation (Q-effect), similar to extracellular lactate via the Warburg effect [14, 17, 132]. Given that the ETS and mSLP are mechanistically entangled, we explored the efficiency of succinate oxidation in both cell types by quantifying extracellular succinate accumulation. Under steady-state ATP sufficiency in glutamine with NAD^+^, CT-2A cells exported significant quantities of succinate, whereas macrophages exported minimal extracellular succinate (Figure 4H). This disparity suggests that primary non-tumor cells may oxidize glutamine more efficiently through canonical OxPhos, while CT-2A tumor cells export succinate as a byproduct of glutamine fermentation.

Unlike macrophages, we noted that the NAD^+^-mediated rescue of CT-2A cells required supplementation with dialyzed serum (Figure S4.2F). Because dialyzed serum contains normal lipid levels, we questioned whether fatty acid oxidation (FAO) was contributing to ATP sufficiency under these conditions. To test this, we treated cells with etomoxir, a well-characterized inhibitor of carnitine palmitoyltransferase 1 (CPT1), the rate-limiting enzyme of FAO (Figure 4I). High-dose etomoxir (200 μM) significantly reduced the OCR of CT-2A cells cultured in glucose and glutamine supplemented with dialyzed serum (Figure 4J). High doses of etomoxir also reduced OCR in the absence of serum (Figure S4.4A), recapitulating previously described off-target effects on Complex I [133, 134]. Crucially, continuous 120-h treatment with up to 200 μM etomoxir had no significant effect on the viability of CT-2A cells cultured in glucose and glutamine (Figure 4K) or glutamine with NAD^+^ (Figure 4L). ATP-linked bioluminescence confirmed that while proliferation was halted in glutamine with NAD^+^, viability was not compromised under high-dose etomoxir (Figure S4.4B). Similar results were observed in macrophages, which did not require serum supplementation for NAD^+^ rescue, but were similarly unaffected by etomoxir when dialyzed serum was provided (Figure S4.4, C, D). Together, these data suggest that glutaminolysis, rather than serum-derived lipid β-oxidation, was the primary bioenergetic pathway sustaining ATP sufficiency in the complete absence of glycolytic sources.

### Steady-state ATP sufficiency without confounding from the Crabtree/Warburg effect enables the quantification of metabolite-specific oxygen consumption

Accurate description of steady-state bioenergetics requires long-term experimental endpoints, but surrogate markers such as oxygen consumption are influenced by confounding from the Crabtree/Warburg effect, proliferation, and cell death under conditions of nutrient stress (Figure 5A). For example, over a 120-h tracking period, quiescent macrophages cultured in glucose and glutamine exhibit lower baseline OCR than those cultured in galactose and glutamine, illustrating glucose-induced suppression of respiration (Crabtree effect) (Figure 5B). OCR is commonly recorded in short-term experimental windows, but respirometry in conditions that ultimately fail to support long-term viability precludes conclusions about ATP sufficiency. Additionally, the presence of glucose in the culture medium can drive proliferation, the Crabtree effect and the Warburg effect, making it difficult to estimate whether baseline OCR is linked to ATP production (Figure 5C). Complex V inhibitors such as oligomycin, traditionally used to estimate ATP-linked OCR, are confounded by the Crabtree/Warburg effect under glucose, or lethal under oxidative conditions such as galactose.

**Figure 5.**
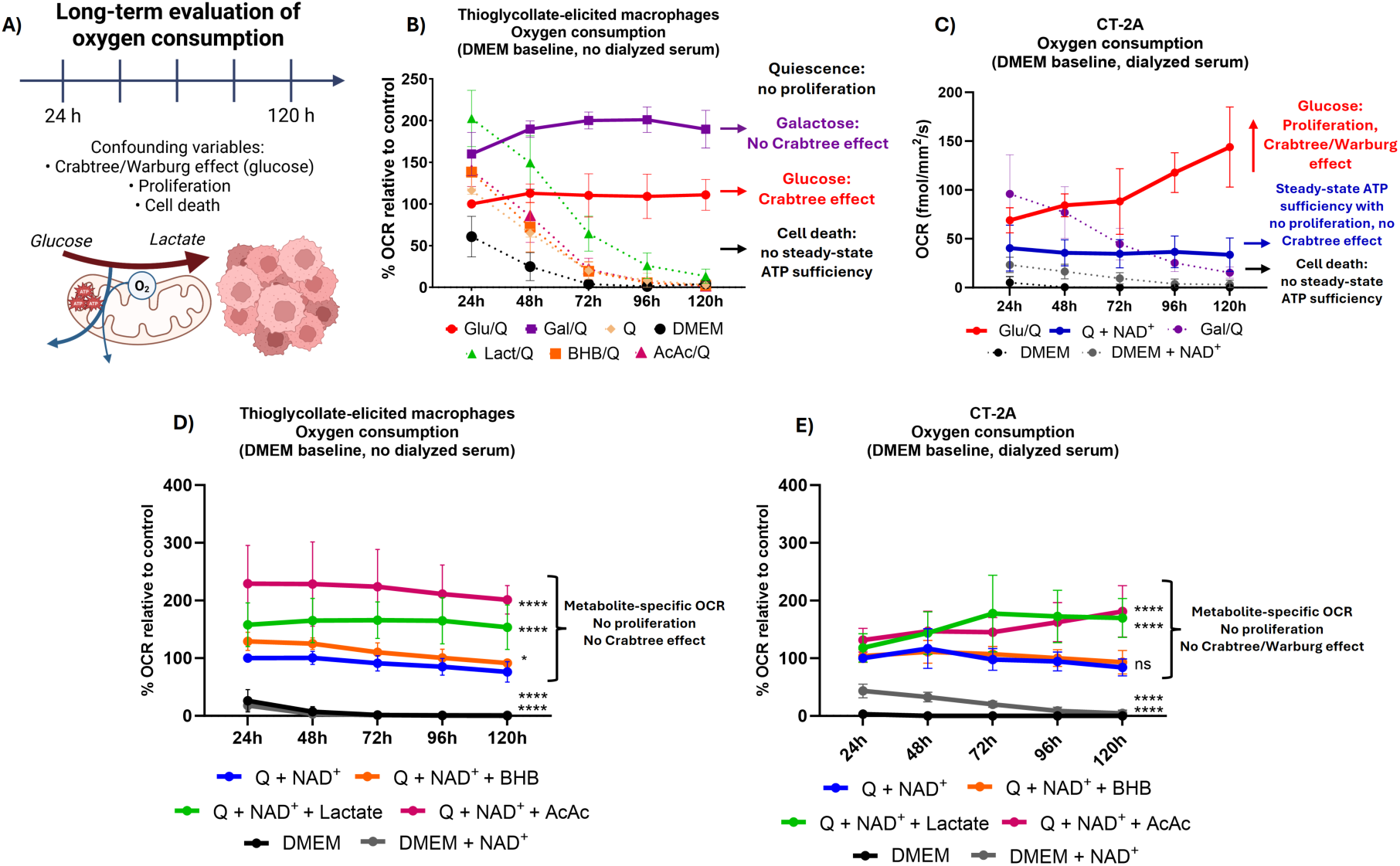
Steady-state ATP sufficiency without confounding from the Crabtree/Warburg effect, proliferation or cell death enables the quantification of metabolite-specific oxygen consumption. **(A)** Experimental design. Evaluating ATP sufficiency requires long-term experimental endpoints, but oxygen consumption rate (OCR) can be influenced by confounding variables such as the Crabtree/Warburg effect, proliferation and cell death. (**B)** OCR of macrophages over 120 h. Glucose and glutamine culturing exhibits lower OCR than galactose and glutamine (Crabtree effect, i.e., glucose-induced suppression of oxygen consumption). OCR can be measured in short-term endpoints (e.g., 24 h), but only metabolite combinations that support long-term viability can evaluate steady-state bioenergetics. **(C)** OCR of CT-2A tumor cells over 120 h. Proliferation and the Crabtree/Warburg effect are potential confounders in the presence of glucose. Culturing cells in glutamine with NAD^+^ enables stable OCR without proliferation or the Crabtree/Warburg effect. **(D-E)** Metabolite-specific OCR under steady-state ATP sufficiency without the Crabtree/Warburg effect. OCR of **(D)** macrophages and **(E)** CT-2A cells cultured in glutamine with NAD^+^ for up to 120 h, with the addition of lactate, acetoacetate, or β-hydroxybutyrate. OCR increased with lactate and acetoacetate, but not β-hydroxybutyrate. Metabolite-specific OCR was not predictive of overall metabolic flexibility during active proliferation in cancer cells (cf. Figure 1). Data represent mean ± SD of at least 3 independent biological replicates. Two-way ANOVA with Dunnett’s correction (D-E) (* p ≤ 0.05, ** p ≤ 0.01, *** p ≤ 0.001, **** p ≤ 0.0001).). Abbreviations: Glu (Glucose), Gal (Galactose), Q (Glutamine), Lact (Lactate), BHB (β-hydroxybutyrate), AcAc (Acetoacetate), DMEM (Dulbecco’s Modified Eagle Medium).

To circumvent these limitations, we applied our experimental design of glutamine with NAD^+^ culturing, halting proliferation and eliminating the Crabtree/Warburg effect (absence of glycolytic sources), effectively “locking” cells into a fixed state of ATP sufficiency. Under this baseline, we introduced individual oxidative metabolites and measured the metabolite-specific OCR in real time for 120 h in macrophages (Figure 5D) and CT-2A cells (Figure 5E). The addition of L-lactate and acetoacetate significantly increased the steady-state OCR in both cell types, whereas the addition of β-hydroxybutyrate did not alter baseline respiration. Consistent with this metabolite-specific OCR, we confirmed that 3-hydroxybutyrate dehydrogenase (BDH1), the mitochondrial enzyme that converts β-hydroxybutyrate to acetoacetate, had low protein expression in macrophages, VM-M3 and CT-2A cells, while 3-oxoacid CoA transferase 1 (OXCT1), the mitochondrial enzyme converting acetoacetate to acetoacetyl-CoA, was expressed in all three cell types (Figure S5.1). Surprisingly, this increase in oxygen consumption was not predictive of overall response to nutrient stress. For example, while CT-2A cells exhibited increased OCR from lactate, they were highly sensitive to glucose deprivation and experienced rapid cell death when cultured in lactate alone (cf. Figure 1). Thus, while both normal and tumor cells present variable rates of metabolite-specific OCR, we propose that bioenergetic flexibility should be determined empirically by comparative analysis of differential responses to metabolic stress.

### Untargeted metabolomics reveals distinct intracellular metabolic rewiring and pathway-specific steady states during nutrient stress

To characterize the metabolic adaptations underpinning steady-state ATP sufficiency across cell lines and nutrient environments, we performed untargeted metabolomic profiling of intracellular metabolites in media conditions that supported long-term viability. Principal component analysis (PCA) demonstrated clear separation and distinct clustering of intracellular metabolite profiles driven by cell identity (macrophages, VM-M3, and CT-2A), proliferation status (DMEM vs. DPBS), and nutrient availability (glucose, galactose and glutamine alone with NAD^+^) (Figure 6A). Hierarchical clustering of the top 25 differentially abundant metabolites further confirmed that environmental nutrient composition was the primary driver of metabolite variance, with minor clustering by cell type (Figure 6B). As expected, DPBS produced distinct clustering marked by depletion of intracellular amino acid pools, despite the presence of dialyzed serum. The complete metabolite dataset is included in Supplementary Table S1.

**Figure 6.**
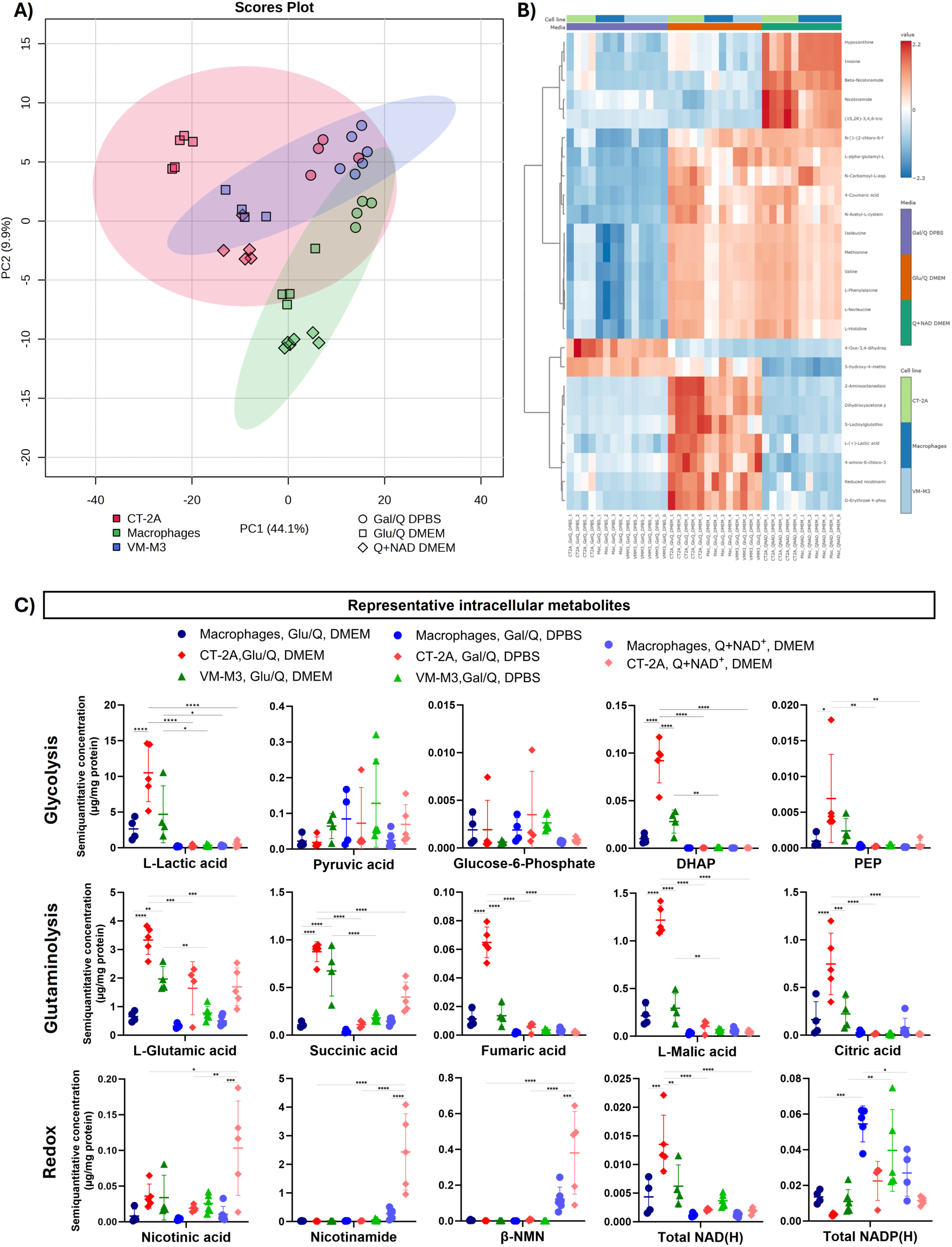
Untargeted intracellular metabolomic profiling across cell types and nutrient environments. **(A)** Principal component analysis (PCA) score plot with distinct clustering of intracellular metabolites from CT-2A, VM-M3, and thioglycollate-elicited macrophages cultured for 48 h in glucose + glutamine (DMEM), galactose + glutamine (DPBS), or glutamine alone with NAD+ (DMEM), supplemented with 10% dialyzed serum. **(B)** Hierarchical clustering heatmap showing the top 25 differentially abundant intracellular metabolites ranked by statistical significance based on linear modeling, grouped by media and cell type. **(C)** Relative abundances of representative intracellular metabolites spanning glycolysis, glutaminolysis, and NAD metabolism after 48 h. Glucose deprivation lowers the lactate/pyruvate ratio. Glutamine catabolism in CT-2A leads to succinic acid accumulation (mSLP) with limited flux into fumaric acid, while exogenous supplementation with NAD^+^ elevates NAD precursors to maintain the NAD(H) pool. Data represent mean ± SD, *n* = 4-6 biological replicates per group. One-way ANOVA with Tukey’s post hoc test, with key biologically relevant comparisons displayed and the complete dataset provided in Supplementary Table S1 (* p ≤ 0.05, ** p ≤ 0.01, *** p ≤ 0.001, **** p ≤ 0.0001). Abbreviations: Glu (Glucose), Gal (Galactose), Q (Glutamine), DMEM (Dulbecco’s Modified Eagle Medium), DPBS (Dulbecco’s Phosphate Buffered Saline).

A detailed evaluation of representative metabolites across key bioenergetic pathways provides mechanistic insight into how cancer and normal cells adapt to nutrient stress (Figure 6C). Glucose deprivation (galactose/glutamine or glutamine with NAD^+^) elicited marked depletion of downstream glycolytic intermediates, including dihydroxyacetone phosphate (DHAP) and phosphoenolpyruvate (PEP), suppressing lactic acid production and reducing the lactate/pyruvate ratio across all cell types, thereby reinforcing the extracellular lactate concentrations measured via enzymatic assay. Unlike macrophages, however, CT-2A cells cultured in glutamine with NAD^+^ exhibited marked upregulation of glutaminolysis (mSLP), characterized by accumulation of succinic acid with limited downstream flux into fumaric acid, malic acid, or reductive carboxylation to citric acid, likewise supporting our previous extracellular succinate quantification. Exogenous supplementation with NAD^+^ elevated intracellular precursors (nicotinic acid, nicotinamide, and β-nicotinamide mononucleotide), thereby supporting the total NAD(H) pool, albeit at lower levels than glycolysis. Conversely, total NADP(H) pools were upregulated under galactose culturing. Together, these data indicate that glutamine acts as a major source of energy and redox homeostasis for proliferating cancer cells (glucose) and non-proliferating cancer cells (glucose deprivation).

### Basal and adaptive changes in the mitochondrial network do not predict OxPhos sufficiency

To determine whether the physical mass of the chondriome dictates bioenergetic flexibility, we quantified changes in the basal and adaptive mitochondrial footprint (total 3D volume) of live cells using MitoView 650 mitochondrial staining. Despite demonstrating a higher degree of metabolic flexibility in previous assays, macrophages exhibited a similarly sized mitochondrial footprint relative to VM-M3 cells, and a significantly smaller mitochondrial footprint compared with CT-2A cells (Figure 7A).

**Figure 7.**
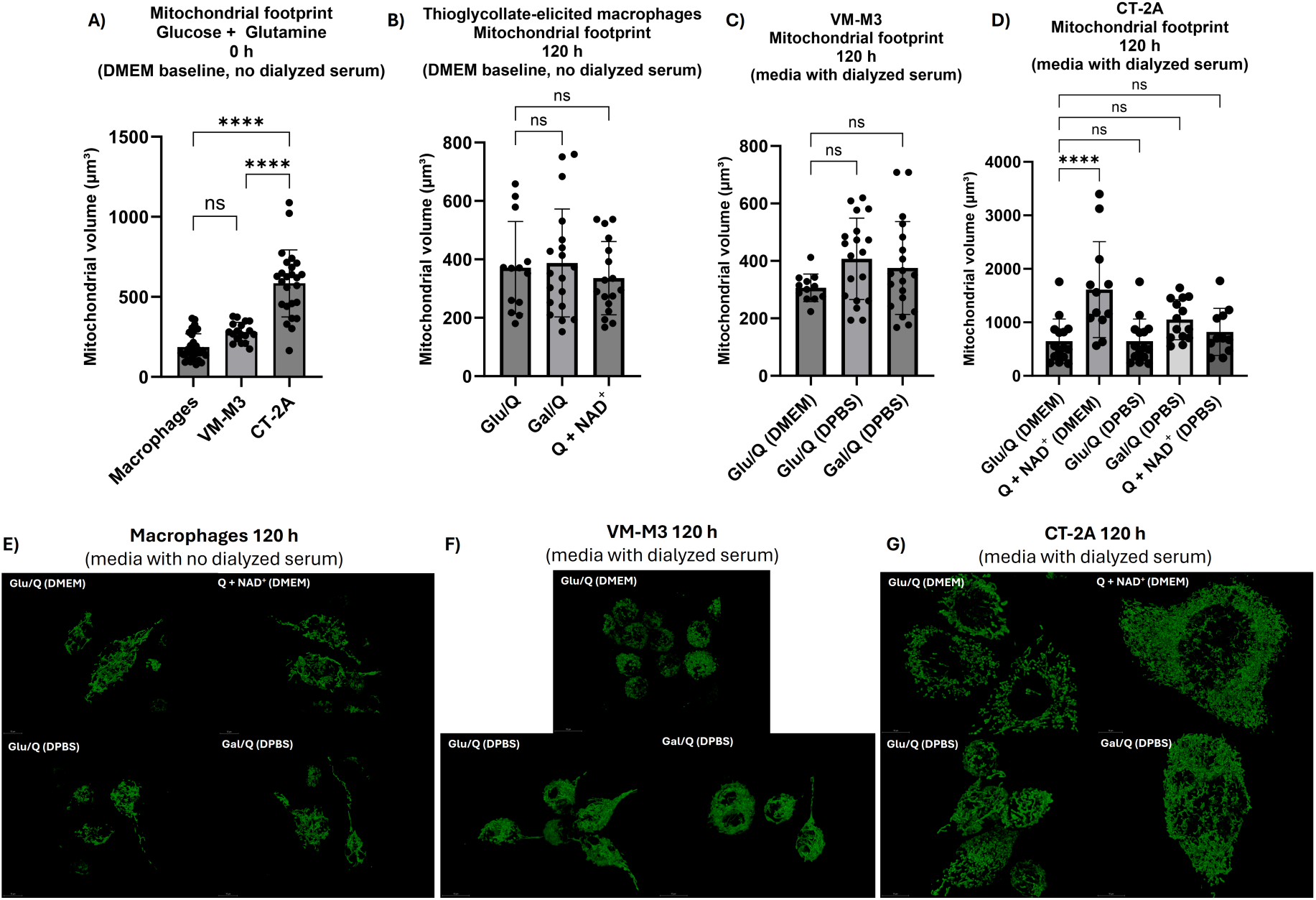
Basal and adaptive changes in the mitochondrial network do not predict OxPhos sufficiency. (A-D) The total mitochondrial footprint (3D volume) was quantified in live cells using mitochondrial staining. **(A)** Total mitochondrial footprint in basal conditions. Macrophages exhibited a comparable mitochondrial footprint to VM-M3 cells, and a significantly smaller mitochondrial footprint than CT-2A cells. **(B)** The mitochondrial footprint of macrophages did not change after 120 h under different nutrient conditions. (**C)** The mitochondrial footprint of VM-M3 cells did not change after 120 h under glucose or galactose DPBS culturing; note that staining could not be performed in other conditions due to cell death. **(D)** In CT-2A, glutamine with NAD^+^ increased the average mitochondrial footprint after 120 h. **(E-G)** Representative 3D visualization of **(E)** macrophages, **(F)** VM-M3 and **(G)** CT-2A cells after 120 h in the indicated conditions. Scale bar: 10 μm. Data represent mean ± SD of individual cells from 3 independent biological replicates. One-way ANOVA with Dunnett’s correction (* p ≤ 0.05, ** p ≤ 0.01, *** p ≤ 0.001, **** p ≤ 0.0001).

To evaluate adaptive morphological responses, we measured the mitochondrial footprint after 120 h in metabolite conditions that supported long-term viability (ATP sufficiency). In primary macrophages, the mitochondrial footprint remained stable after culturing in glucose and glutamine, galactose and glutamine, or glutamine with NAD^+^ (Figure 7B). This lack of adaptive morphological changes was also observed for the mitochondrial branch length (Figure S7.1B). Furthermore, extended profiling confirmed limited changes in the total 3D volume or branch length across combinations of basal media (DMEM versus DPBS) and dialyzed serum (Figure S7.2, A, B). Representative 3D renderings of mitochondria illustrate this morphological stability (Figure S7.3).

Analogously, VM-M3 cells demonstrated a stable mitochondrial footprint in response to glucose (DMEM and DPBS) and galactose (DPBS) (Figure 7C). Mitochondrial branch length remained unchanged in VM-M3 across viable conditions (Figure S7.1C). On the other hand, CT-2A cells cultured in glutamine with NAD^+^ experienced a significant increase in total mitochondrial footprint, suggesting a structural adaptation to glutaminolysis as the primary bioenergetic pathway (Figure 7D). While CT-2A cells cultured in DPBS (galactose and glutamine or glutamine with NAD^+^) did not show a clearly enlarged mitochondrial footprint, they demonstrated a significant increase in mitochondrial branch length compared with glucose and glutamine controls (Figure S7.1D). Representative 3D images illustrate the differences in mitochondrial networks across macrophages, VM-M3, and CT-2A (Figure 7, E-G).

### Non-proliferating macrophages exhibit a transient Crabtree effect, whereas proliferating VM-M3 and CT-2A tumor cells display a persistent Warburg effect

To distinguish between physiological, reversible suppression of respiration (Crabtree effect), and the corresponding metabolic phenotype in cancer (Warburg effect), we analyzed oxygen consumption dynamics under relative glucose deprivation (Figure 8A). Both phenomena are characterized by high glycolytic flux with enhanced cytosolic SLP and lactic acid fermentation. Glucose concentrations as low as 5 mM can elicit the Crabtree effect [63, 135], but the Crabtree and Warburg effects are rarely disentangled experimentally.

**Figure 8.**
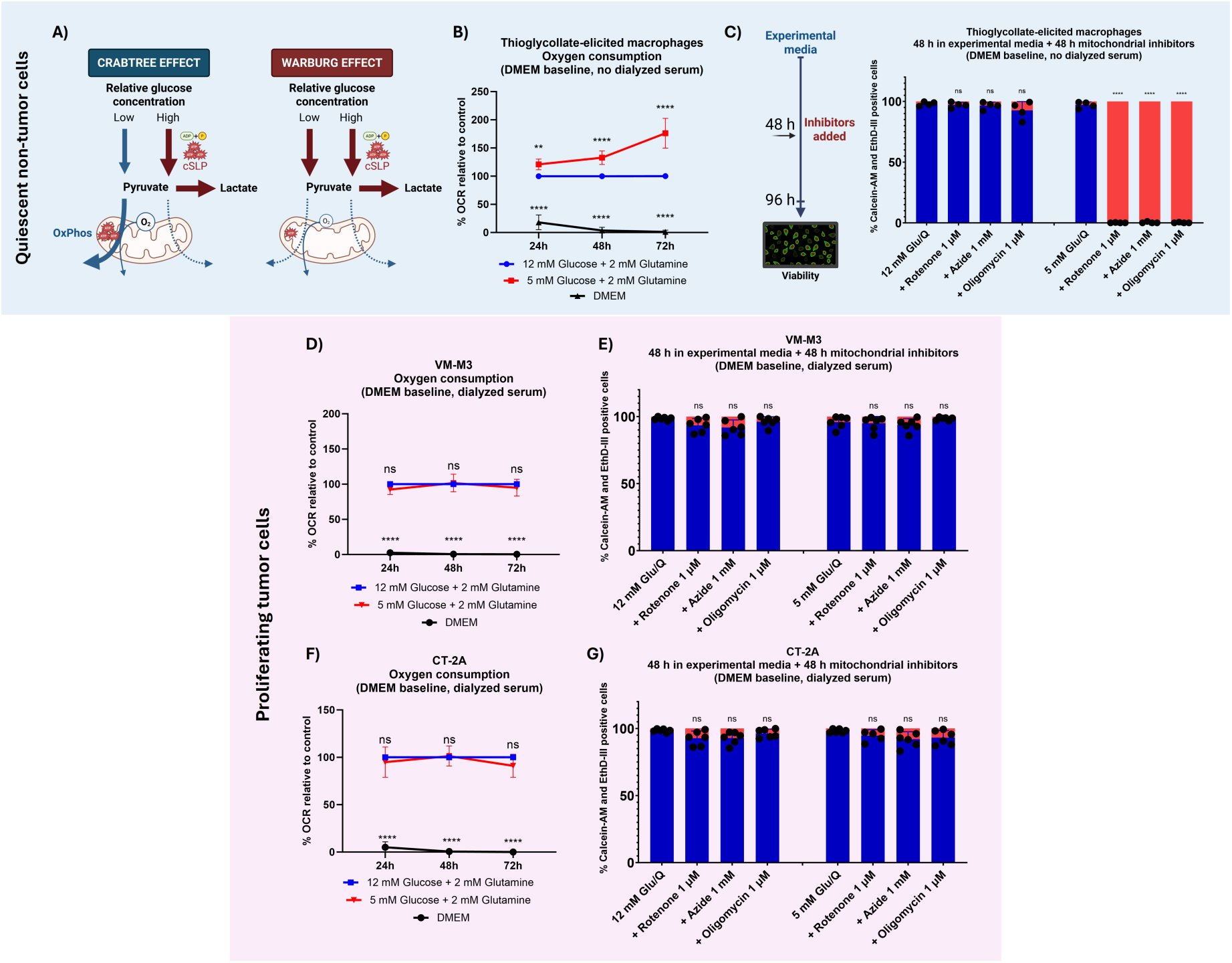
Non-proliferating macrophages exhibit a glucose concentration- and time-dependent Crabtree effect, whereas proliferating VM-M3 and CT-2A tumor cells display a persistent Warburg effect. **(A)** Conceptual schematic contrasting the Crabtree and Warburg effects. The Crabtree effect is defined as the reversible, glucose-induced suppression of respiration accompanied by lactic acid fermentation. The Warburg effect is also characterized by lactic acid fermentation and dependency on cytosolic SLP, but its regulation by glucose, time, and proliferation remains to be defined experimentally. **(B–C)** Metabolic phenotype of macrophages. **(B)** Oxygen consumption rate (OCR) of macrophages cultured in 12 mM or 5 mM glucose over 72 h. Freshly isolated macrophages in glucose-containing medium display the Crabtree effect regardless of glucose concentration, but OCR gradually increases when glucose is below 5 mM. **(C)** Macrophages are resistant to mitochondrial inhibitors when cultured for 48 h in 12 mM glucose but become sensitive when cultured in 5 mM glucose. **(D–E)** Metabolic phenotype of VM-M3 cells. **(D)** OCR of proliferating VM-M3 cells cultured in 12 mM or 5 mM glucose over 72 h, showing comparable patterns regardless of glucose concentration. **(E)** Viability of VM-M3 cells cultured in 12 mM and 5 mM glucose for 48 h and treated with mitochondrial inhibitors for 48 h in the same conditions. VM-M3 cells were resistant to ETS inhibition. **(F-G)** Metabolic phenotype of CT-2A cells. A comparable phenotype was observed in proliferating CT-2A cells for **(F)** OCR and **(G)** resistance to mitochondrial inhibitors. Data represent mean ± SD of at least 3 independent biological replicates. Two-way ANOVA with Dunnett’s correction (B, D, F) and one-way ANOVA with Dunnett’s correction (C, E, G) (* p ≤ 0.05, ** p ≤ 0.01, *** p ≤ 0.001, **** p ≤ 0.0001). Abbreviations: Glu (Glucose), Q (Glutamine), DMEM (Dulbecco’s Modified Eagle Medium).

Thus, we cultured cells in standard “high-glucose” (12 mM) and physiological “low-glucose” (5 mM) environments. In quiescent macrophages, the introduction of glucose-containing medium rapidly elicited a robust Crabtree effect. At short-term endpoints (24 h), freshly isolated macrophages maintained suppressed OCR regardless of glucose concentration (Figure 8B). Consequently, macrophages cultured for 24 h in 5 mM glucose remained resistant to mitochondrial inhibitors, relying on cytosolic SLP to meet ATP demands regardless of dialyzed serum (Figure S8.1, A, B). However, continuous longitudinal tracking over 72 h revealed a distinct, glucose concentration- and time-dependent OCR divergence. Macrophages cultured in 5 mM glucose progressively bypassed the Crabtree effect, steadily increasing oxygen consumption over time. By 48 h, macrophages cultured in 5 mM glucose transitioned to dependency on OxPhos and became sensitive to mitochondrial inhibitors, whereas those cultured in 12 mM glucose remained fully resistant (Figure 8C). This dynamic metabolic shift and time-dependent separation in oxygen consumption were conserved when the medium was supplemented with dialyzed serum (Figure S8.2, A, B).

In contrast, proliferating VM-M3 and CT-2A tumor cells displayed a persistent Warburg phenotype. Neither VM-M3 (Figure 8D) nor CT-2A cells (Figure 8F) exhibited separation or compensatory increase in OCR over a 72-h period when cultured in 5 mM versus 12 mM glucose. Despite cell proliferation driving glucose consumption, tumor cells failed to upregulate respiratory flux. As a result, after 48 h of culturing, both VM-M3 (Figure 8E) and CT-2A (Figure 8G) remained resistant to ETS inhibition in both high and low glucose conditions. The absence of dialyzed serum did not alter this reliance on cytosolic SLP, as both malignant cell lines maintained non-diverging OCR profiles and resistance to mitochondrial inhibitors (Figure S8.2, C-F).

### Dysregulated proliferation, but not regulated proliferation, is associated with the Warburg effect

To determine whether the Warburg effect may be a common feature of all proliferating cells [4] or a pathological hallmark specific to cancer, as originally proposed by Warburg [1], we evaluated the bioenergetics of primary macrophages stimulated to undergo regulated proliferation (Figure 9A). Upon stimulation with the mitogen macrophage colony-stimulating factor (M-CSF), macrophages exhibited a gradual increase in cell number over 10 days in glutamine paired with either glucose or galactose (Figure 9B). Analogous to tumor cells, supplementation with dialyzed serum was required for sustained proliferation. The expansion in cell density following M-CSF stimulation is illustrated via fluorescence microscopy (Figure S9.1). Proliferative capacity was confirmed through protein quantification and optimized via M-CSF dose-response titrations (Figure S9.2, A, B). Interestingly, M-CSF-stimulated macrophages displayed a relative decrease in normalized extracellular lactate accumulation in glucose, while producing little lactate when proliferating in galactose (Figure 9C, Figure S9.2C). When stimulated with M-CSF for 48 h in 5 mM glucose and subsequently treated with mitochondrial inhibitors for 48 h, proliferating macrophages still bypassed the Crabtree/Warburg effect and experienced complete loss of viability (Figure 9D). Prolonged M-CSF exposure for 120 h prior to mitochondrial inhibition yielded the same results (Figure S9.3A). This suggests that regulated proliferation in macrophages may rely on OxPhos for steady-state ATP sufficiency and is not intrinsically associated with the Warburg effect.

**Figure 9.**
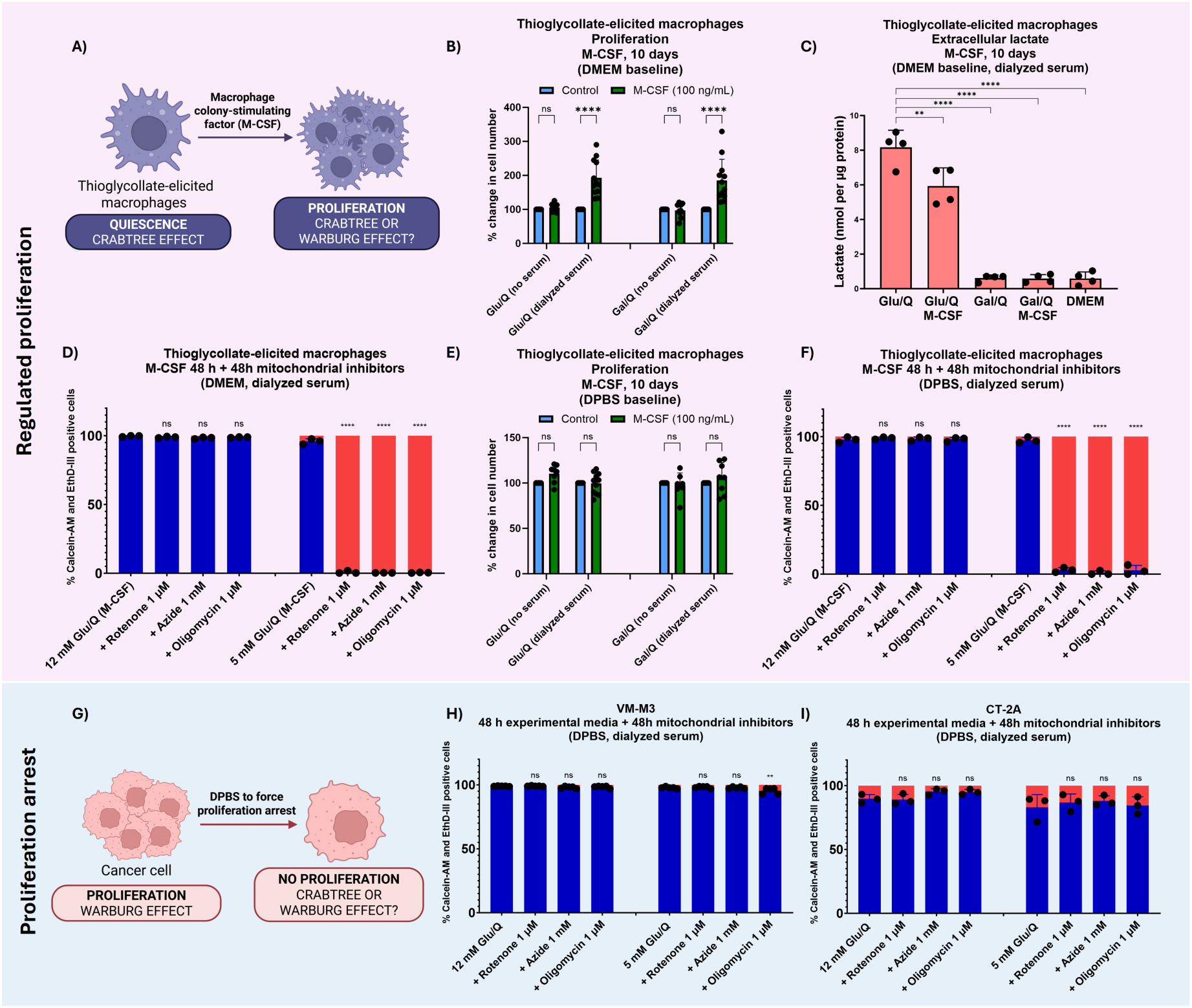
Proliferating macrophages exhibit a reversible Crabtree effect and regulated growth via OxPhos, whereas growth-arrested VM-M3 and CT-2A tumor cells display a persistent Warburg effect. **(A)** Quiescent macrophages exhibit the Crabtree effect, but this phenotype could be influenced by proliferation. **(B)** Regulated proliferation of macrophages stimulated with macrophage colony-stimulating factor (M-CSF) over 10 days. Cell number increased in glucose and glutamine as well as galactose and glutamine in DMEM with dialyzed serum. **(C)** Extracellular lactate over 10 days. M-CSF-stimulated macrophages show a relative decrease in normalized lactate production when cultured in glucose and minimal lactate in galactose. **(D)** Viability of M-CSF-stimulated macrophages exposed to mitochondrial inhibitors. Complete loss of viability was observed in macrophages cultured for 48 h in 5 mM glucose, regardless of M-CSF stimulation. **(E)** DPBS induced growth arrest despite M-CSF and dialyzed serum. **(F)** M-CSF-stimulated macrophages cultured in DPBS remained sensitive to mitochondrial inhibitors in 5 mM glucose. **(G–I)** Growth-arrested tumor cells. **(G)** Proliferating VM-M3 and CT-2A cells exhibited a persistent Warburg effect, but this phenotype could be influenced by proliferation arrest. Viability of **(H)** VM-M3 and **(I)** CT-2A cells cultured for 48 h in DPBS and treated with mitochondrial inhibitors for 48 h. Growth-arrested cancer cells remained resistant to mitochondrial inhibitors even when cultured in 5 mM glucose. Data represent mean ± SD of at least 3 independent biological replicates. Two-way ANOVA with Šidák’s correction (B, E) and one-way ANOVA with Dunnett’s correction (C, D, F, H, I) (* p ≤ 0.05, ** p ≤ 0.01, *** p ≤ 0.001, **** p ≤ 0.0001). Abbreviations: Glu (Glucose), Q (Glutamine), DMEM (Dulbecco’s Modified Eagle Medium), DPBS (Dulbecco’s Phosphate Buffered Saline)

To isolate the bioenergetic compartment from the demands of cell division, we forced M- CSF-stimulated macrophages into growth arrest via DPBS culturing (Figure 9E). Growth-arrested mitogen-stimulated macrophages maintained the respiratory bioenergetic shift, with high sensitivity to mitochondrial inhibitors after 48 h of culturing in 5 mM glucose (Figure 9F). We then induced growth arrest in VM-M3 and CT-2A tumor cells to determine whether their metabolic profile would resemble quiescence (Figure 9G). Cultured for 48 h in DPBS medium with dialyzed serum and subsequently challenged with mitochondrial inhibitors, VM-M3 (Figure 9H) and CT-2A (Figure 9I) remained resistant to ETS blockade in both 12 mM and 5 mM glucose. These results persisted in the absence of dialyzed serum (Figure S9.3, B-D). Thus, unlike growth-regulated cells that efficiently adapt to nutrient cues and metabolic demands, the examined cancer cells demonstrated a more rigid dependency on cytosolic SLP regardless of glucose concentration or active proliferation.

## Discussion

The goal of cancer treatment is to normalize or eliminate cancer cells, but the goal of cancer research is also to study their behavior, which is considerably more interesting when they are alive. It should therefore come as no surprise that standard culturing techniques provide cancer cells with abundant nutrients, optimizing for proliferation and robustness. Classic media formulations such as DMEM were developed to maximize the “growth and multiplication of mammalian cells” [136], not to define the minimum constraints on ATP sufficiency [137, 138]. This could be viewed as mimicking the nutrient-rich extracellular environments in multicellular organisms, which have evolved complex structures and systems (e.g., gluconeogenesis) to maintain metabolic homeostasis [139, 140]. From an evolutionary perspective, unlike unicellular organisms, individual cells residing in metazoans are less vulnerable to acute fluctuations in environmental resources [141]. Even so, multiple overlapping pathways have evolved across all strata of life to buffer transient disruption of ATP, as immediate cell death upon minor energy stress would hinder biological inheritance [142]. The question therefore arises whether cells with dysregulated proliferation exhibit differences in bioenergetic flexibility compared to regulated cells, and whether the genetic and mitochondrial abnormalities that accumulate in cancer cells confer a survival advantage or limit adaptive versatility under nutrient stress.

Our experimental methodology placed special emphasis on steady-state ATP sufficiency. Sustained viability is generally achieved by culturing cells in glucose and glutamine, thus enabling the study of other metabolites or drug interventions, but is inherently confounded by the Crabtree/Warburg and Q-effect. The Crabtree effect can be circumvented by substituting glucose with galactose, which limits the rate of cytosolic SLP and is accompanied by a rapid increase in oxygen consumption [143]. While this spike in oxygen consumption is often interpreted as a successful transition to OxPhos, most cancer cells eventually die or fail to proliferate in galactose medium [69, 71, 130, 144–147]. In some cases, the use of glucose-containing, non-dialyzed serum with frequent medium replacement may have confounded results [71, 148–152], as elegantly demonstrated by Skolik et al. [153]. Interestingly, non-tumorigenic HepG2 hepatoma cells proliferate slowly over 20-30 weeks in medium supplemented with galactose, glutamine, pyruvate and dialyzed serum, but continue producing a baseline level of lactate [153]. Conversely, primary cells such as lymphocytes, fibroblasts and muscle cells have been described to sustain long-term viability in galactose-based medium [131, 154–158]. Short-term respirometry and isotope labeling (without accounting for baseline cytosolic and mitochondrial SLP dependency) are frequently referenced to suggest that lactate, fatty acids, ketones and other oxidative metabolites are major fuels for cancer cells [29–31, 33]. We show that despite not fully replacing glycolytic substrates, as previously reported [159, 160], lactate and ketone bodies contributed to long-term viability in macrophages when combined with glutamine and dialyzed serum. On the other hand, proliferating tumor cells exhibited rapid loss of viability under identical conditions. Our results highlight that short-term experimental endpoints, relative changes in oxygen consumption, as well as mitochondrial content are not predictive of bioenergetic flexibility under partial or complete SLP inhibition, which should be empirically determined by exposing tumor cells and normal counterparts to nutrient stress.

It has been proposed that biosynthesis is rate-limiting for proliferation under nutrient-rich conditions, where dividing cells balance the kinetic rate of ATP synthesis via cytosolic SLP with the preservation of carbon backbones for macromolecular synthesis, eliciting the Warburg effect despite functional OxPhos [4, 161]. By uncoupling these variables, our results challenge the assumption that “aerobic glycolysis” is a universal feature of all proliferating cells [4, 162]. We demonstrate that primary non-tumor macrophages display a transient, glucose-dependent Crabtree effect that resolves over time, enabling reliance on mitochondrial ATP generation during regulated proliferation. While mitogen-stimulated primary cells have been shown to upregulate glycolytic flux to support biosynthetic demands [163, 164], our data suggest that this may not necessarily involve a Warburg phenotype. The biosynthetic argument for the Warburg effect relies on *in vitro* evidence that normal proliferating cells, including enterocytes, stem cells, endothelial cells, and stimulated lymphocytes and thymocytes, can exhibit upregulated aerobic glycolysis and suppressed oxygen consumption [41, 160, 165–175]. However, such extrapolations were derived from *in vitro* conditions and thus inadvertently confounded by the Crabtree effect.

In mammalian cells, the Crabtree effect is elicited in seconds to minutes upon addition of glucose, which is consistent with flux mechanisms rather than changes in protein expression [62, 153, 176]. Consistent with their specialized physiological roles, macrophages shift their bioenergetic phenotype towards cytosolic SLP (glycolysis) upon encountering acute hypoxia [177]. However, despite collection from a hypoxic niche in the peritoneal cavity [178–180], thioglycollate-elicited peritoneal macrophages effectively upregulated OxPhos upon exposure to standard *in vitro* oxygen tensions. While freshly isolated macrophages were used for most studies, we also verified that the cell culture environment, including penicillin/streptomycin exposure [181], did not impair OxPhos after 8-10 weeks of culture, suggesting that the phenotypic differences were not related to cell culture maintenance. Other non-tumor primary cells, such as astrocytes and fibroblasts, have been described as producing lactate via the Crabtree effect but subsequently transition to OxPhos under glucose depletion [182, 183].

Our demonstration that primary macrophages have the capacity to sustain proliferation using OxPhos without the Crabtree effect (galactose) or with a reversible Crabtree effect (low glucose) aligns with recent evidence that normal proliferating cells actively upregulate OxPhos to meet bioenergetic demands [184]. However, it is possible that the maximal proliferation rate also influences the Warburg phenotype, as mitogen-stimulated macrophages display exponential growth but retain contact inhibition [96, 102, 185]. Additionally, normal cells preserve homeostatic control to balance environmental and nutrient cues with cell cycle progression, while tumor cells, by definition, do not [186–188]. Future studies could apply a similar methodology to define the Crabtree effect in other primary cell types with rapid turnover, such as intestinal crypts or other epithelial cells, which can exceed the proliferation rate of tumor cells under physiological conditions [189–191].

In contrast, the evaluated malignant cells exhibited a persistent Warburg phenotype that was conserved across both active proliferation and forced growth arrest. It is possible, however, that exposure to lower glucose concentrations over time would increase the sensitivity to mitochondrial inhibitors, as demonstrated by continuous-flow culture techniques that maintain constant glucose levels in the medium [192, 193]. Interestingly, cancer cells under such conditions still demonstrated a Warburg effect (extracellular lactate) and absolute dependency on cytosolic and mitochondrial SLP for proliferation, even as glucose levels were kept very low (e.g., 0.75 mM). Lactate production was also demonstrated at the single-cell level using genetically encoded lactate sensors, with higher accumulation in glioma cells than in normal astrocytes, irrespective of proliferation [194]. Hexose transporters and glycolytic enzymes such as hexokinase II and lactate dehydrogenase may account for baseline differences in fermentative flux across cancer cells [195–197], but do not explain why these traits appear unresponsive to nutrient stress, which could be considered an evolutionary maladaptation and therapeutic vulnerability across fluctuating environmental contexts.

The ATP requirements of proliferation compared with cellular maintenance continue to be debated [2, 8, 26–28, 198]. By forcing cell-cycle arrest in cancer cells via amino acid and vitamin deprivation [199–201], we reveal that active dysregulated proliferation was associated with lower metabolic flexibility. From a thermodynamic perspective, halting cell division may lower global ATP demand, allowing VM-M3 and CT-2A “non-proliferating” tumor cells to fuel steady-state ATP sufficiency using galactose and glutamine with minimal lactic acid fermentation. We further show that cell cycle arrest via the targeted CDK4/6 inhibitor palbociclib paradoxically improved viability under galactose culturing. Ferguson et al. reported that while palbociclib re-engaged differentiation in neuroblastoma cells, complete cell-cycle arrest was not achieved [202]. Furthermore, palbociclib can induce arrest at different phases of the cell cycle [203]. Thus, the metabolic rescue achieved via CDK4/6 inhibition may not perfectly phenocopy DPBS due to the inherent heterogeneity of cyclin-dependent kinases across tumor cells [204]. Overall, our results support the recent work by Kukurugya et al. showing that inhibition of translation (protein synthesis) via cycloheximide reduced total ATP demand, implying that a significant portion of ATP consumption at high growth rates may be associated with new biomass synthesis [26]. By examining the transplantability of ascites after short-term nutrient deprivation, Warburg himself speculated that “cancer cells require much less energy to keep them alive than they do for growth” [1].

In our models, reintroducing pro-proliferative components restored proliferation but triggered cell death under oxidative culturing. While non-glutamine amino acids significantly contribute to carbon mass in proliferating cells *in vitro* [205], our findings identify a clear bioenergetic and biosynthetic hierarchy: non-glutamine amino acids fail to support viability in the absence of glucose and glutamine, whereas glucose and glutamine are necessary and sufficient for ATP maintenance, but do not enable sustained proliferation in the absence of amino acids, vitamins and serum components. It is well established that cancer cells require serum or specific growth factors (e.g., insulin, transferrin and selenium) for continued propagation in serum-free media [128, 129]. We note that while dialyzed serum was required for active proliferation in both tumor and non-tumor cells, it was not required for baseline ATP sufficiency. Given that serum is inherently present *in vivo*, restricting individual serum elements may be impractical. We highlight that the absolute dependency of cancer cells on glycolysis and glutaminolysis for both viability and proliferation, regardless of intracellular fate or mechanism, should be prioritized over other secondary metabolic pathways [9, 10].

It is important to note, however, that while ATP sufficiency can be claimed under sustained viability, cell death may be unrelated to bioenergetics [6, 21], as demonstrated by the rescue of cytosolic redox balance with NAD^+^ under total glucose deprivation. Our results indicate that the Warburg effect can be bypassed in some cell types by directly supplying the redox end-product of lactic acid fermentation (NAD^+^) into the extracellular space, revealing that glutaminolysis can drive steady-state ATP sufficiency in macrophages and CT-2A cells. The complete absence of glycolytic sources (glucose or galactose) also arrests proliferation, although a limited number of cell divisions may be completed by scavenging intracellular or exogenous precursor pools [206, 207]. Extracellular NAD^+^ at supraphysiological concentrations (500 μM) has been shown to successfully enter mammalian cells [208]. Extracellular NAD^+^ can enter cells via connexin hemichannels [208, 209] or ectoenzyme-mediated degradation (CD38, CD73) into oxidized precursors, which are subsequently resynthesized into NAD^+^ inside the cell [210]. Both mechanisms preserve the total NAD(H) pool, as observed in our metabolomic profiling. Conversely, extracellular NADH degrades into reduced nicotinamide riboside (NRH), elevating the reduced cytosolic pool [211] and potentially exacerbating cell death under conditions of limited redox buffering capacity.

Furthermore, the potential contribution of fatty acid oxidation (FAO) from dialyzed serum was excluded using etomoxir, with both CT-2A and macrophages presenting unimpaired viability despite high-dose (200 μM) etomoxir treatment under glutamine and NAD^+^; of note, macrophages did not require serum for viability but were similarly unaffected when it was provided. Previous pharmacological profiling established that low concentrations of etomoxir (10 μM) are sufficient to inhibit > 90% CPT1 activity and halt FAO [134]. Recent evidence also suggests that high-dose etomoxir can act as a promiscuous fatty acid mimetic that elicits off-target effects on respiratory Complex I [133, 134, 212, 213]. The preservation of viability under complete FAO blockade and partial Complex I inhibition suggests that glutaminolysis, rather than lipid β-oxidation, was the primary ATP source under these conditions. This reinforces the notion that the lipid droplets frequently observed in cancer cells are not necessarily linked to energy metabolism [214], since the contribution of FAO to ATP sufficiency was seemingly dispensable.

Oxygen consumption is not a precise quantitative marker of ATP synthesis through Complex V due to confounding from cell division, the Crabtree/Warburg effect (cytosolic SLP), uncoupling, non-mitochondrial OCR, variable metabolic demand (e.g., growth signals), redox activity, oxygen diffusion, and multiple mathematical assumptions required to convert OCR into tentative energy [32, 82, 91, 215, 216]. Nevertheless, in extended experimental setups where substrates are controlled, it is reasonable to assume that a portion of basal OCR is correlated with ATP production. We report that macrophages and CT-2A cells cultured in glutamine with NAD^+^, without confounding from cell death, proliferation or the Crabtree/Warburg effect, exhibited a stable level of OCR that was responsive to substrate addition. Metabolite-specific OCR was consistent with ketolysis enzyme expression, as well as LDHB expression, which has been previously confirmed in thioglycollate-elicited peritoneal macrophages [217] and CT-2A cells [218]. This provides a more comprehensive quantification of metabolite-specific OCR but does not indicate whether any given OCR is sufficient or insufficient to meet the bioenergetic, redox or biosynthetic demands of proliferation.

Principal component analysis of intracellular metabolites indicated congruent clustering based on culture media and cell identity, with glycolysis and NAD^+^ responses aligning with enzymatic measurements to glucose deprivation and exogenous NAD^+^. Galactose induced the depletion of downstream glycolytic intermediates and enhanced reliance on glutamine catabolism, consistent with previous reports [73, 153]. However, glutaminolysis-derived metabolites also suggest a role for mSLP by driving succinate accumulation in CT-2A cells without concurrent accumulation of fumarate or malate, showcasing differential mitochondrial rewiring. Similarly, we noted that the total mitochondrial mass was not predictive of bioenergetic flexibility. Macrophages and VM-M3 cells contain a similarly sized mitochondrial footprint, while CT-2A cells display a larger network, but all three differ in overall responses to metabolic stress. The accumulation of mitochondrial mass without commensurate functional capacity may indicate impaired efficiency or mitophagy [151, 219–222]. We also observed that aglycemic galactose culturing did not significantly expand the mitochondrial footprint, in accordance with Schmidt et al. [150]. Nevertheless, we detected an increase in the mitochondrial footprint in CT-2A cells relying predominantly on glutaminolysis to meet their bioenergetic demands.

In line with this, we show that CT-2A cells produced significant quantities of extracellular succinate under glutamine and NAD^+^, whereas macrophages did not. Importantly, steady-state ATP sufficiency under these conditions excluded confounding from cell death. Succinate export may reflect reduced OxPhos capacity and/or increased mSLP [17, 75], a truncated form of the TCA cycle where glutamine is fermented to succinate, yielding ATP via succinyl-CoA ligase independent of the proton motive force [12, 223, 224]. However, continuous mSLP flux requires Complex I-driven intramitochondrial NAD^+^ regeneration via forward electron transport to preserve α-ketoglutarate dehydrogenase activity [75, 225], as evidenced by complete loss of viability upon ETS inhibition. While mitochondrial malate dehydrogenase, mitochondrial diaphorases and other matrix oxidoreductases have been described to transiently bypass ETS blockade to regenerate matrix NAD^+^ [226], chronic inhibition depletes finite quinone pools, precipitating mSLP collapse [74, 78]. Intact NAD^+^ could also be imported into mitochondria from the cytosol, although the efficiency of this transport is debated [227–229]. Reductive carboxylation via IDH1/2 contributes to macromolecular synthesis under ETS impairment [227, 230], but bypasses succinyl-CoA ligase and thus yields no ATP via mSLP. Consequently, although multiple mechanisms may support intramitochondrial redox cycling during acute stress such as transient hypoxia, such adaptations were insufficient for sustained viability under simultaneous cytosolic SLP and ETS inhibition. Future work could examine whether uncoupling intramitochondrial NAD^+^ regeneration from electron transport using genetic tools such as mitoLbNOX [231] or ideally more efficient NAD^+^ import [228] would allow glutamine-driven mSLP to meet total cellular ATP demands, mirroring cytosolic SLP.

While we provide a comprehensive exploration of the Warburg/Crabtree effect across matched non-tumor and tumor cells, our approach has several limitations that could be addressed in future studies. First, although macrophages were paired with VM-M3 cell biology (myeloid origin), we did not examine normal astrocytes for juxtaposition with CT-2A astrocytoma. We did explore an intermediate state of transformation with myc/raf-immortalized BV-2 microglia, which shows a high proliferation rate *in vitro* but is non-tumorigenic *in vivo* [232]. Previous studies reported enhanced glycolysis with LPS stimulation [233] and short-term survival in galactose [234]. In our hands, BV-2 cells under galactose culturing spontaneously halted proliferation and preserved ATP sufficiency despite pro-growth factors. Thus, mitochondrial ATP generation in BV-2 was sufficient for baseline viability when cells appropriately engaged energy-sensing checkpoints to arrest the cell cycle, which may be one of the differentiating features between immortalization and malignant transformation [235, 236]. Persistent and unresponsive cell cycle progression (malignancy) due to mitochondrial abnormalities and oncogenic signaling may account for the inability to downregulate proliferation when pro-growth requirements have been met (e.g., amino acids from DMEM, growth factors from serum) [237, 238]. Additionally, we focused exclusively on the species *Mus musculus* to ensure a direct comparison between primary and malignant cells. While ATP-generating mechanisms are evolutionarily conserved across mammalian cells [13, 239], mouse cells have faster metabolic rates than human cells [127, 240]. Based on preliminary findings with U-87MG astrocytoma (data not shown), we note that even longer experimental endpoints may be required in human cells to ensure bioenergetics are not confounded by transient adaptations. Lastly, our goal was to define conditions of steady-state ATP sufficiency in viable cells, not to investigate the cell death mechanisms of dying cells. While causal inference regarding ATP depletion is plausible when cells are deprived of all nutrients, cell death under specific nutrient deprivation (such as glucose) may occur for reasons beyond ATP (e.g., redox imbalance). Thus, in our view, true steady-state bioenergetics should be examined in cells with sustained, unimpaired viability.

Several explanations have been put forward for the origin of the Warburg effect since it was first described in 1923 [241]. Biochemical and molecular descriptions (e.g., upregulated protein expression, such as lactate dehydrogenase) can retroactively justify phenomenological observations (e.g., high lactate production), but do not necessarily explain their primary cause (“why is lactate dehydrogenase elevated?”), which can be interpreted from either the somatic mutation theory or the mitochondrial metabolic theory of cancer [242–245]. Our results are consistent with reports suggesting that one of the roles of aerobic glycolysis may be to satisfy energy demand in the context of limited OxPhos capacity [46, 51]. Our findings can also be related to the ultrastructural (cristolysis) and lipidomic (cardiolipin) alterations described in the VM-M3 and CT-2A tumors as well as human gliomas more broadly [246–251], which could serve as the molecular basis for OxPhos insufficiency [13].

Warburg originally proposed that respiratory insufficiency was the common metabolic hallmark of cancer, leading not only to compensatory fermentation, but also to the origin of cancer itself through the “dedifferentiation of life” from eukaryotic multicellular respiration to “unicellular fermentation energy” [41, 245, 252, 253]. Indeed, evidence that restoring OxPhos via chemogenetic tools or mitochondrial transplantation inhibits tumorigenicity supports the notion that OxPhos sufficiency, leading to a reduction in SLP, may act as a tumor suppressor [254–260]. Subsequent biochemical evaluations by Weinhouse argued that surrogate markers of OxPhos, such as oxygen consumption, electron transport or enzymatic activity, were seemingly intact in some tumors, with both increased glycolysis and respiration necessary for the energy economy of the cancer cell [261, 262]. Although Warburg initially used the term “irreversible respiratory damage”, sparking lasting controversy, during his later work he reclassified the respiration of cancer as “insufficient” [263]. This nomenclature may be preferable because “sufficiency” is a functional definition that is agnostic to mechanistic minutiae and encompasses the entire chondriome, energy production versus demand, as well as the vast array of mitochondrial functions (e.g., biosynthesis, redox homeostasis, signaling). In reconciliation of both perspectives, while our data support the role of the respiratory chain (“Ox”) to drive mSLP and redox balance via glutaminolysis, we highlight SLP dependency as the primary bioenergetic phenotype of dysregulated proliferation. This is consistent with reports of continued proliferation upon limited ATP yield from Complex V (“Phos”), as long as cytosolic and mitochondrial SLP are operational [87–90].

In summary, the Crabtree effect could be defined as a glucose-dependent increase in glycolytic flux (cause) leading to a transient suppression of OxPhos (consequence). On the other hand, the Warburg effect may represent a chronic loss of OxPhos sufficiency (cause) leading to a compensatory increase in SLP flux (consequence). With this in mind, we return to Warburg’s central idea that the endeavor of cancer research should be “to discover the differences between cancer cells and normal growing cells” [41]. From a therapeutic perspective, if ATP sufficiency is the primordial prerequisite for life, it would be of interest to systematically categorize the metabolic adaptability of the least flexible normal cell compared with the most flexible cancer cell. We may learn that the “proliferation advantage” of cancer cells is no advantage after all.

## Abbreviations

AcAc: Acetoacetate
BDH1: 3-hydroxybutyrate dehydrogenase
BHB: *β*-hydroxybutyrate
CPT1: Carnitine palmitoyltransferase 1
DHAP: Dihydroxyacetone phosphate
DMEM: Dulbecco’s Modified Eagle Medium
DPBS: Dulbecco’s Phosphate Buffered Saline
EthD-III: Ethidium homodimer III
ETS: Electron transport system
FAO: Fatty acid oxidation
M-CSF: Macrophage colony-stimulating factor
MiNA: Mitochondrial Network Analysis
mSLP: Mitochondrial substrate-level phosphorylation
NAD+: Oxidized nicotinamide adenine dinucleotide
NADH: Reduced nicotinamide adenine dinucleotide
NRH: Reduced nicotinamide riboside
OCR: Oxygen consumption rate
OXCT1: 3-oxoacid CoA-transferase 1
OxPhos: Oxidative phosphorylation
PCA: Principal component analysis
PEP: Phosphoenolpyruvate
PI: Propidium iodide
SLP: Substrate-level phosphorylation
SPE: Solid-phase extraction
TCA: Tricarboxylic acid
UHPLC-MS/MS: Ultra-High Performance Liquid Chromatography-Tandem Mass Spectrometry

## Declarations

### Author contributions

T.D. conceived, designed, and performed the experiments; analyzed the data; and wrote the manuscript. E.Y.M., L.T., C.L., M.B.R., M.B.P., C.K., and O.S. assisted with data collection.

J.J.A.H. and M.K. performed the nontargeted metabolomics. D.C.L. and P.M. assisted with reviewing the manuscript and figures. T.N.S. reviewed and edited the manuscript, acquired funding, and supervised the project. All authors of this manuscript have provided consent for publication.

## Acknowledgments

We thank the Foundation for Metabolic Cancer Therapies, Dr. Edward Miller, Dr. Joseph Maroon, Robert Corkin Charitable Foundation, Elizabeth Ann Weathers Breast Cancer Research Fund, Beth El Pancreatic Cancer Research Fund, Campbell Family Foundation, Children with Cancer UK, Kenneth Rainin Foundation, Delaware County Special Deputies Benevolent Fund, Broken Science Initiative, and the Boston College Research Expense Fund for their support.

We thank Bret Judson and the Boston College Imaging Core for their assistance with microscopy. We thank Dr. Patrick Autissier and the Flow Cytometry Core Facility for their assistance with flow cytometry. Graphical representations were created with BioRender.com.

## Availability of data and material

All data supporting the findings of this study are available within the manuscript and its supplementary material.

