## Supplementary Figures for "Disentangling the Crabtree and Warburg effects across syngeneic non-tumor and tumor cells"

### Slide 1
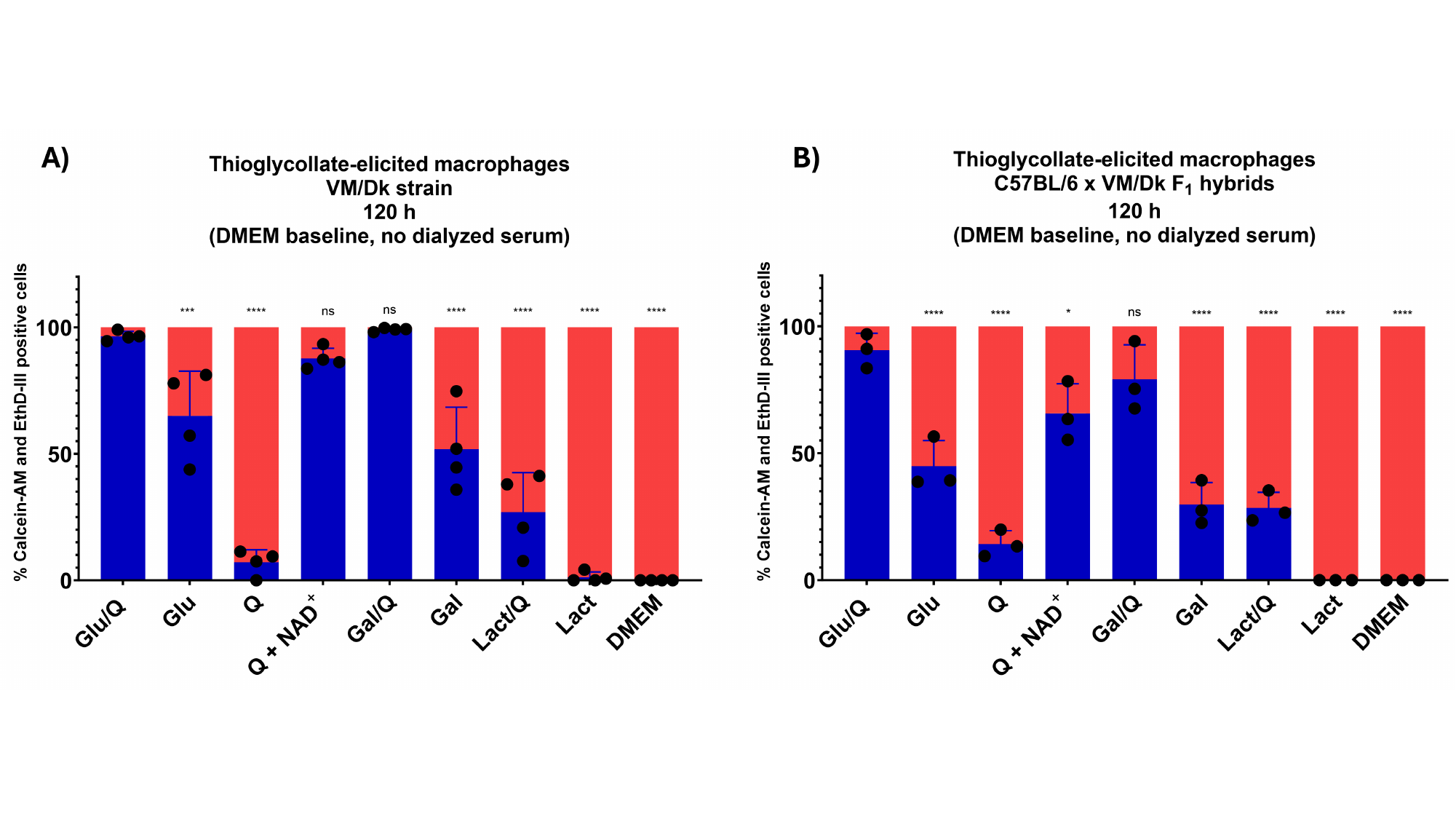

### Slide 2
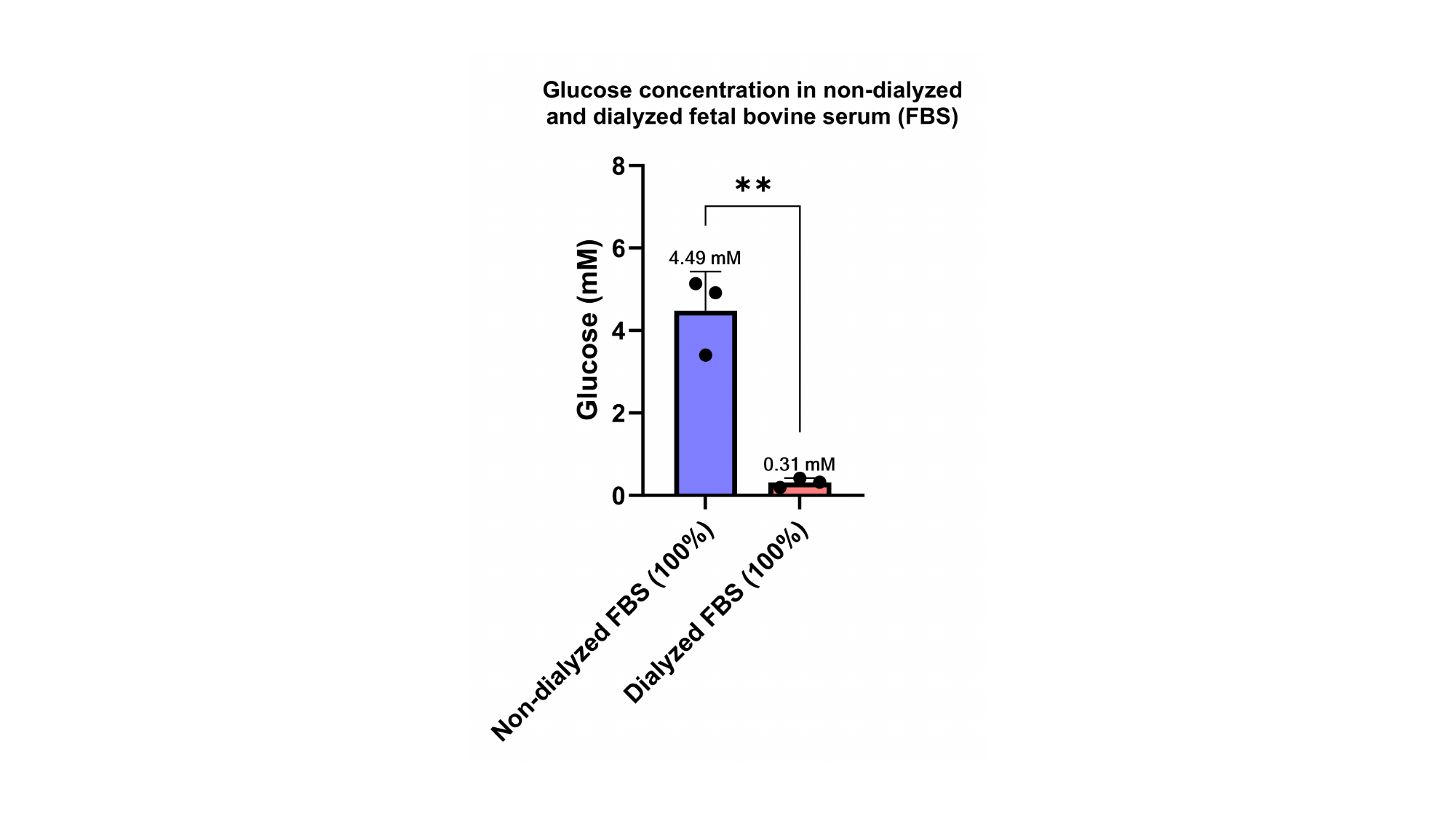

### Slide 3
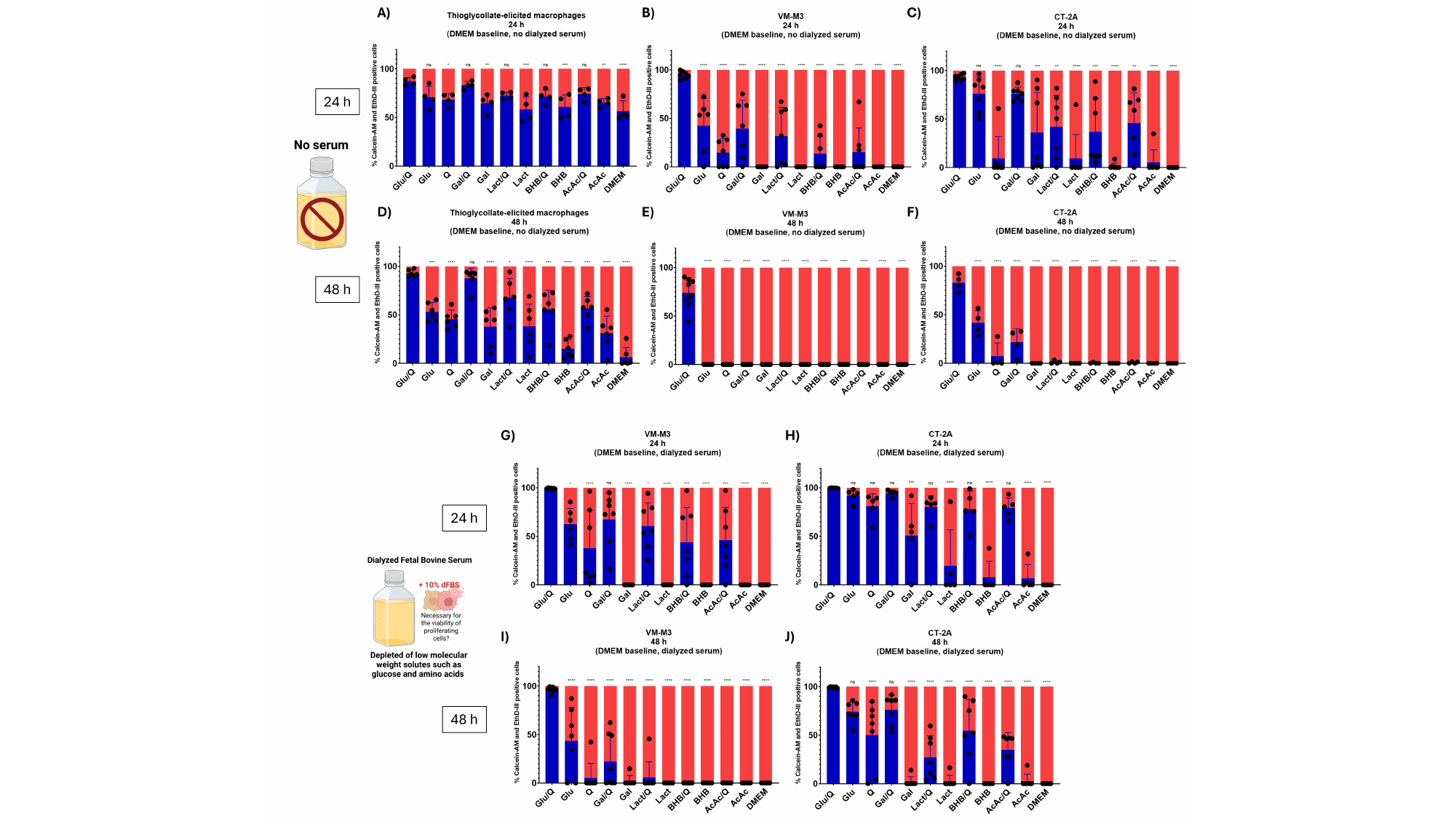

### Slide 4
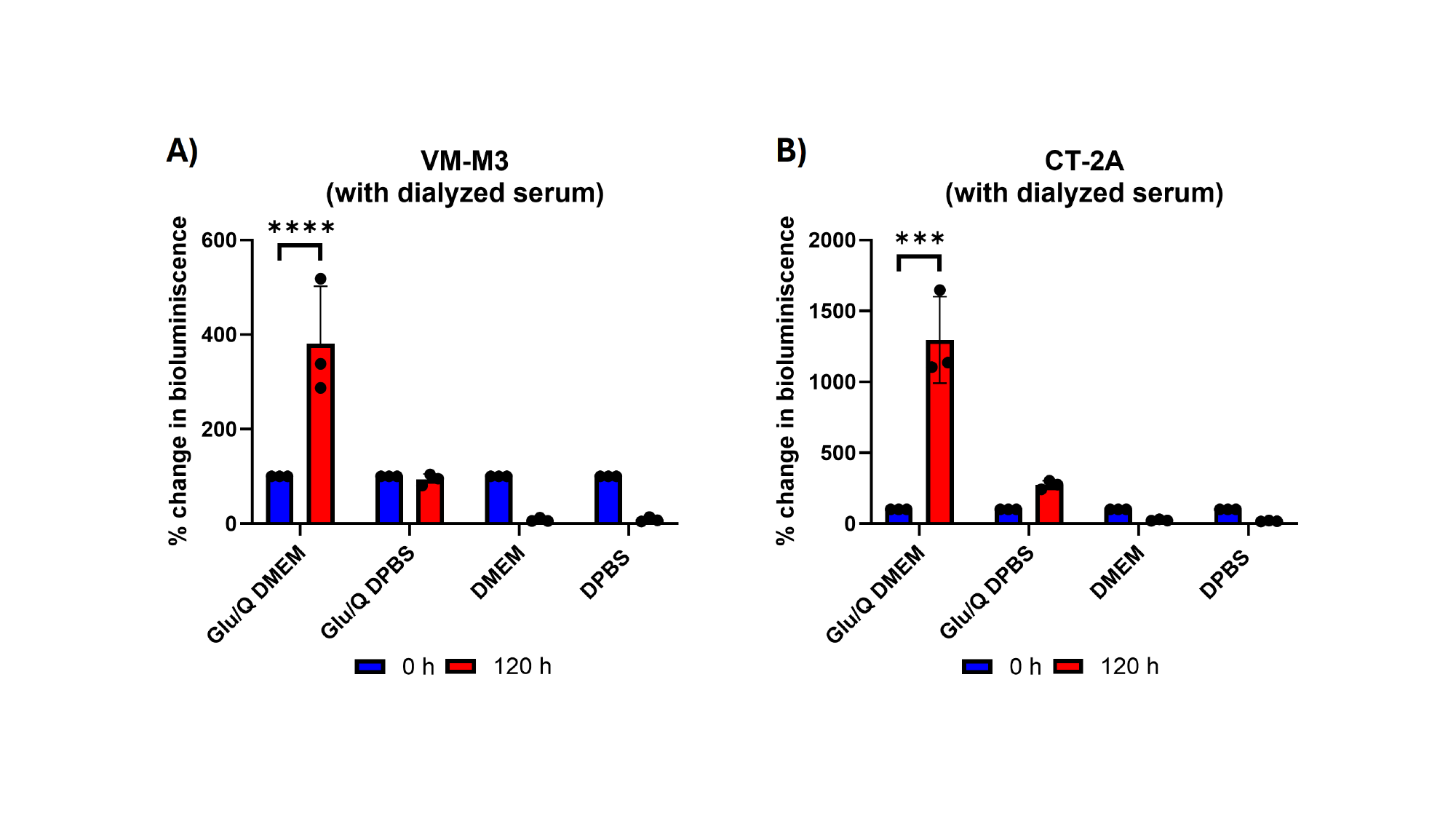

### Slide 5
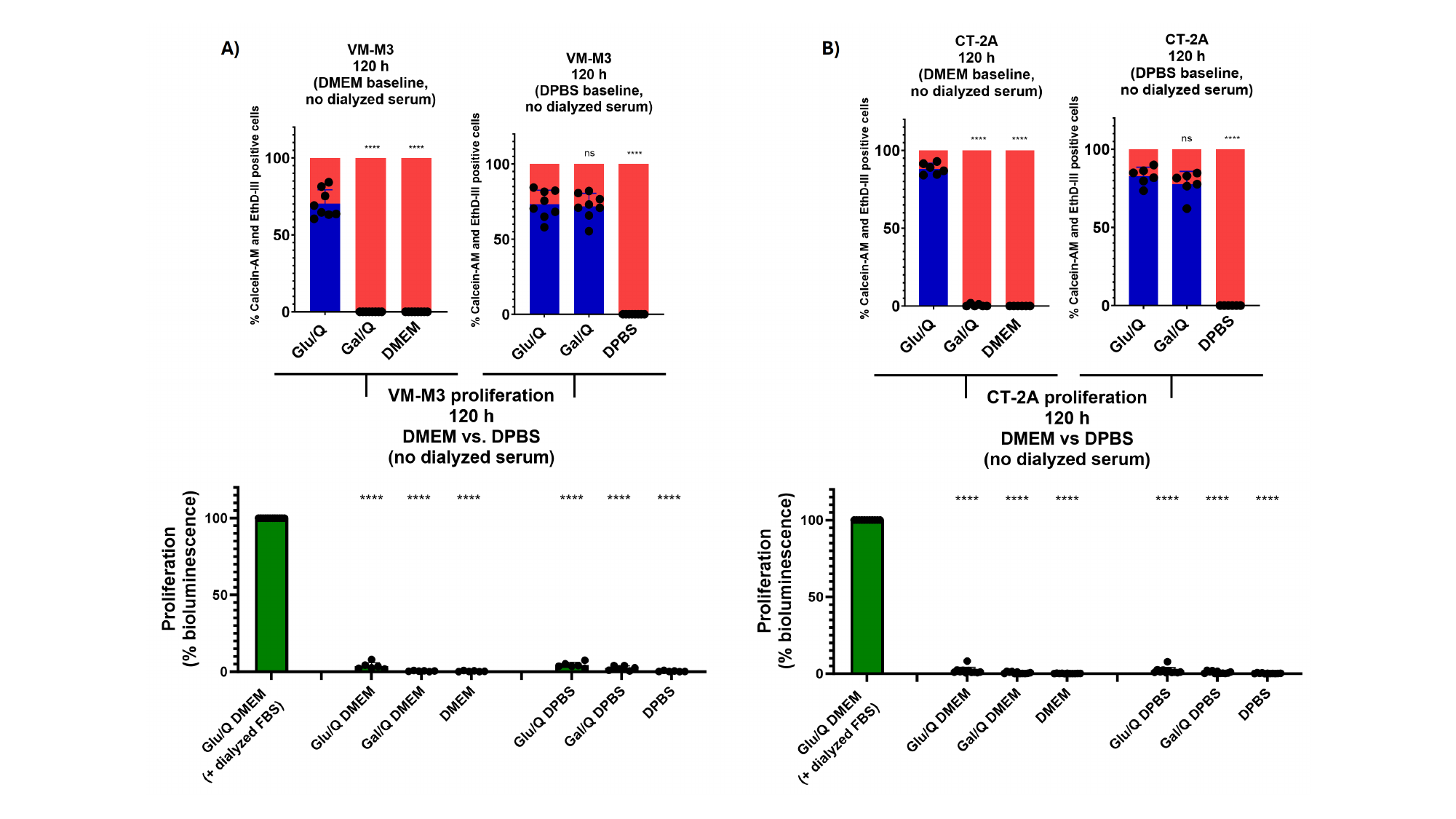

### Slide 6
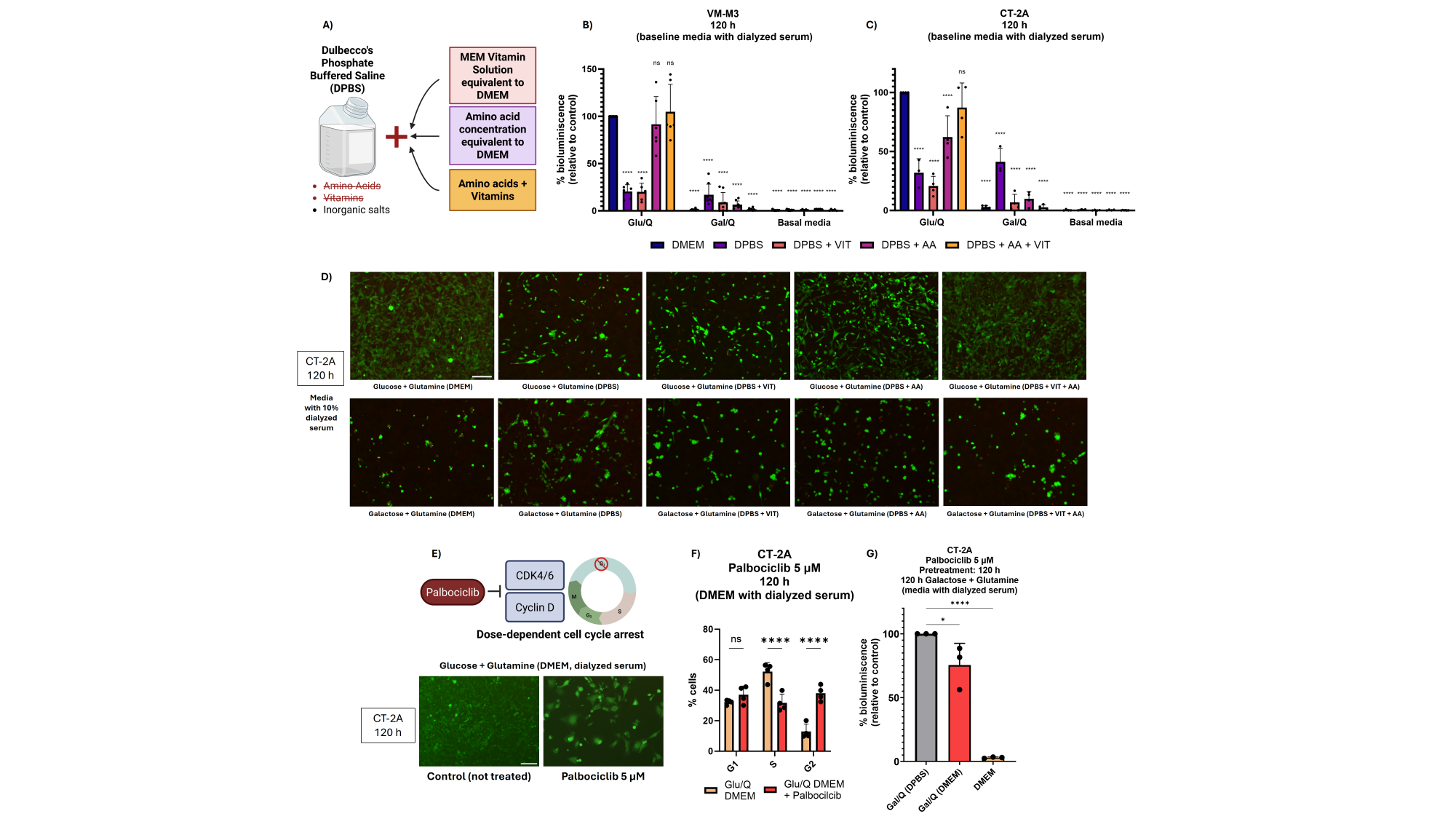

### Slide 7
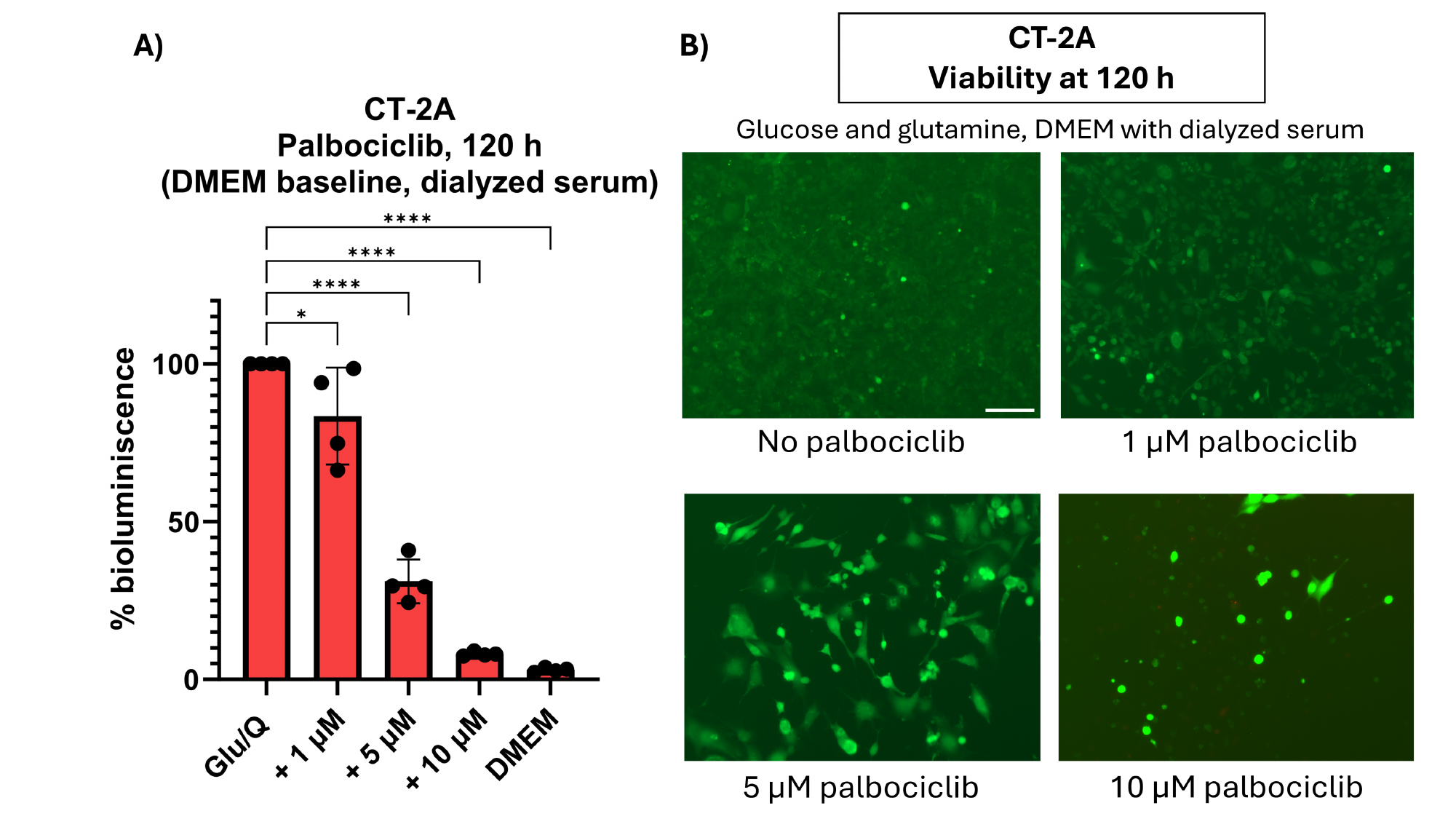

### Slide 8
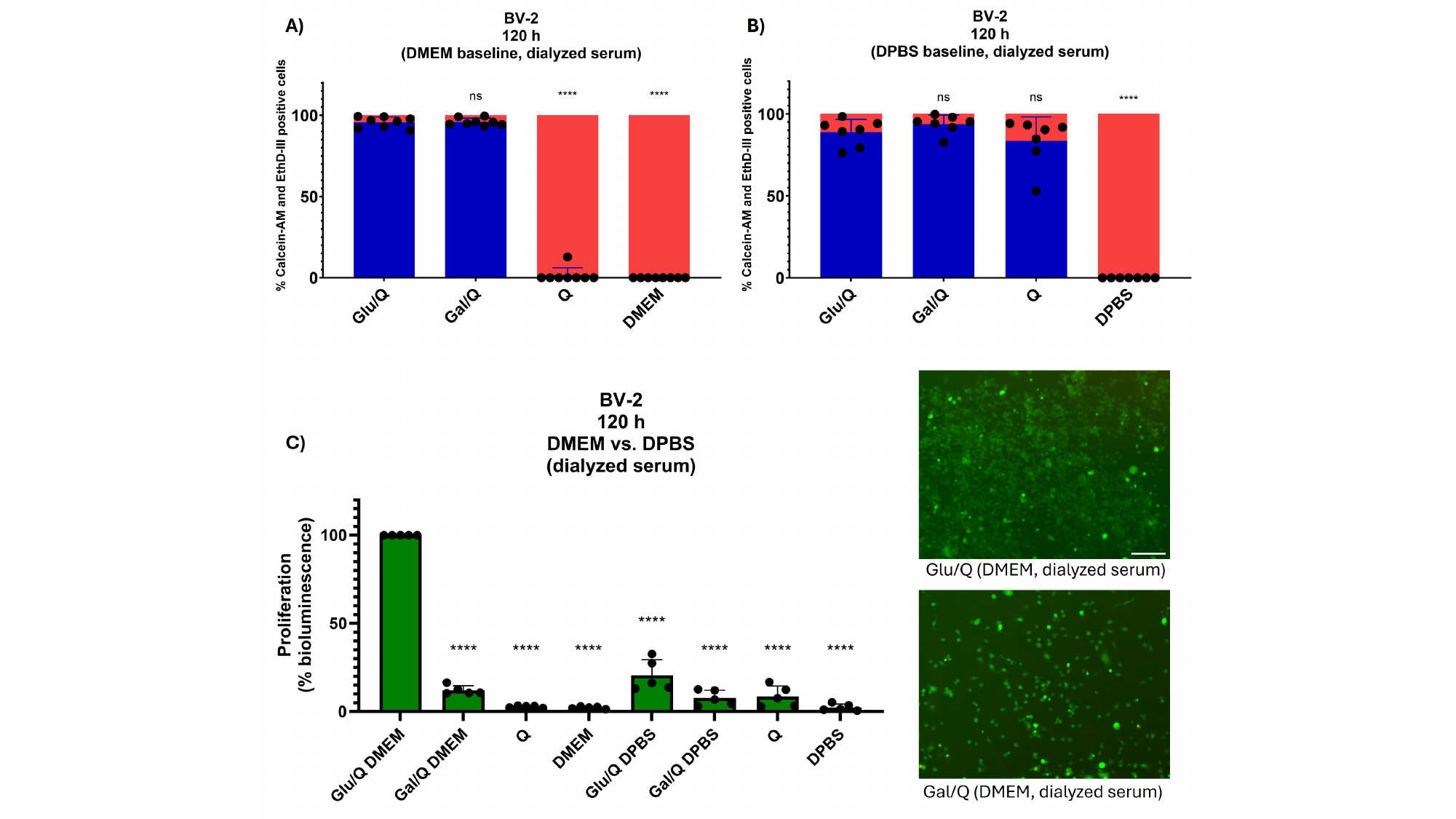

### Slide 9
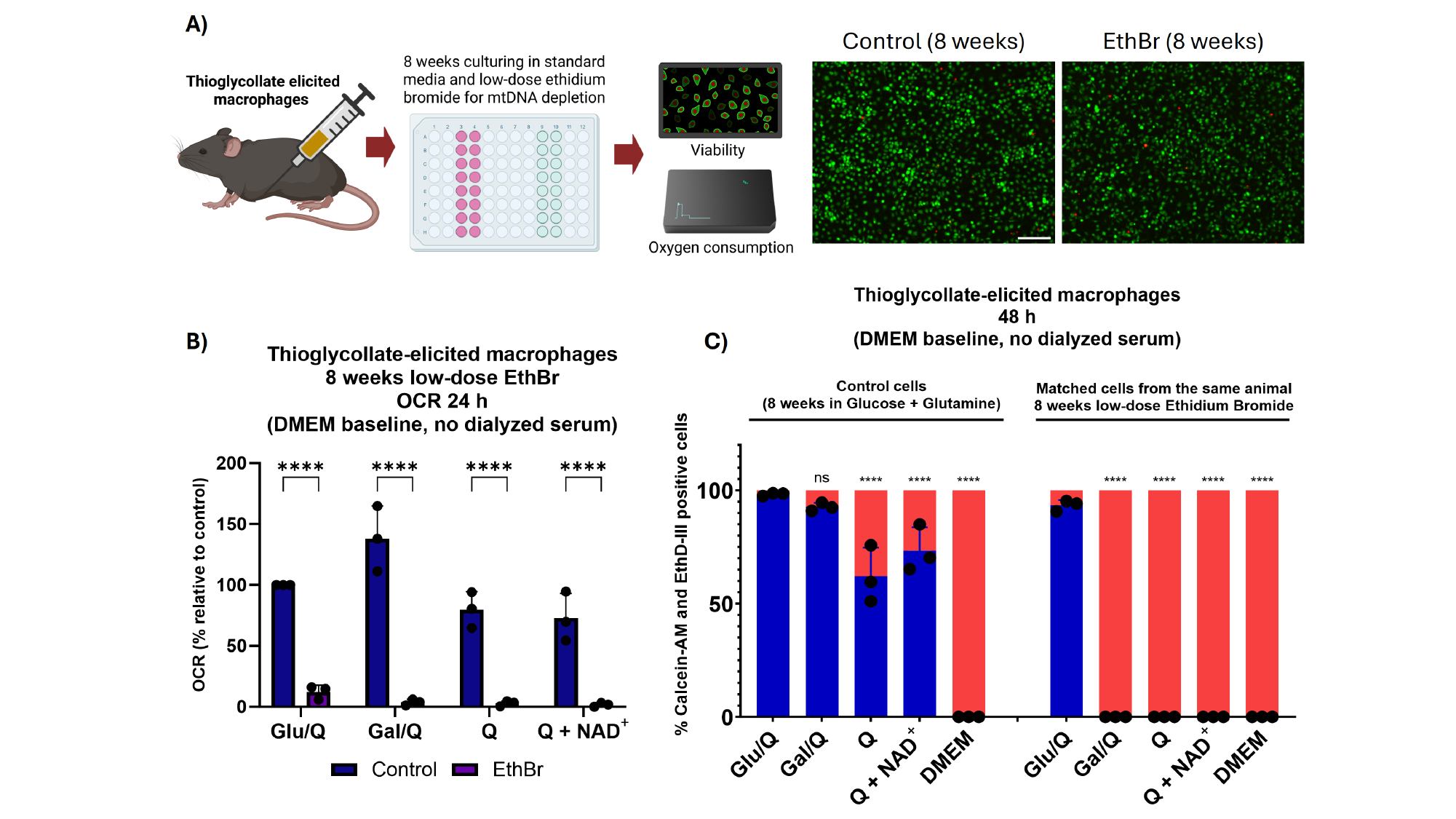

### Slide 10
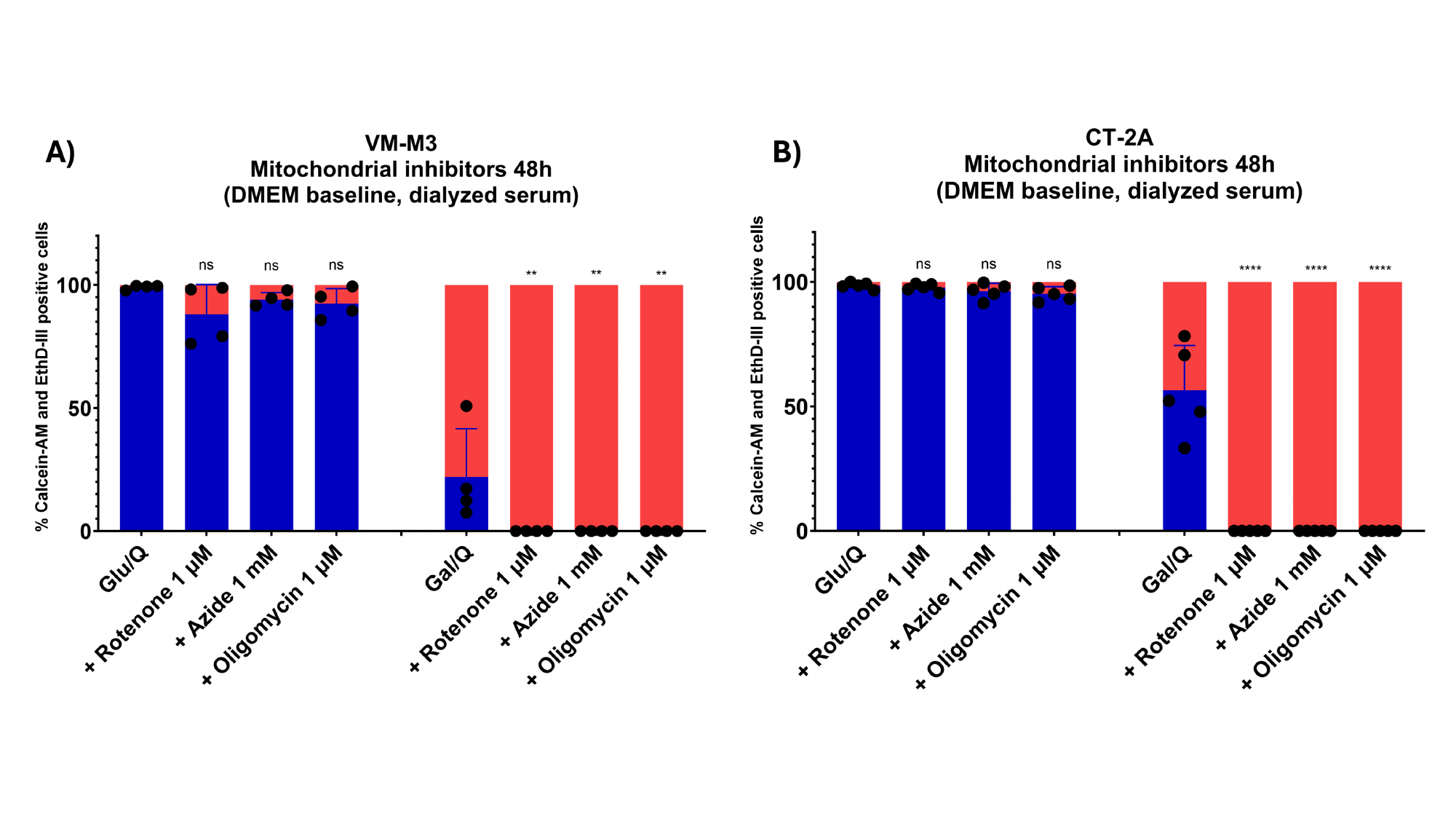

### Slide 11
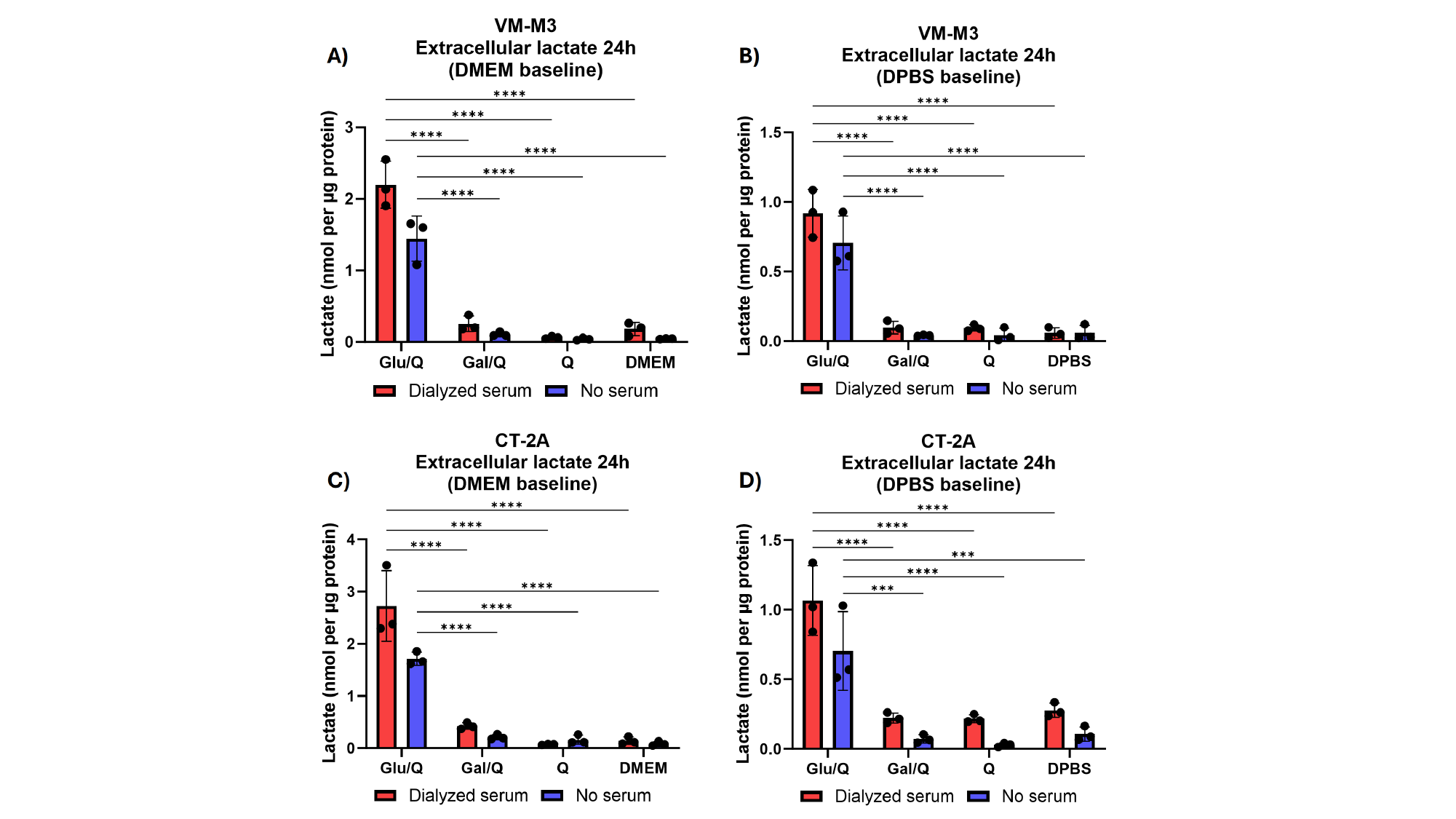

### Slide 12
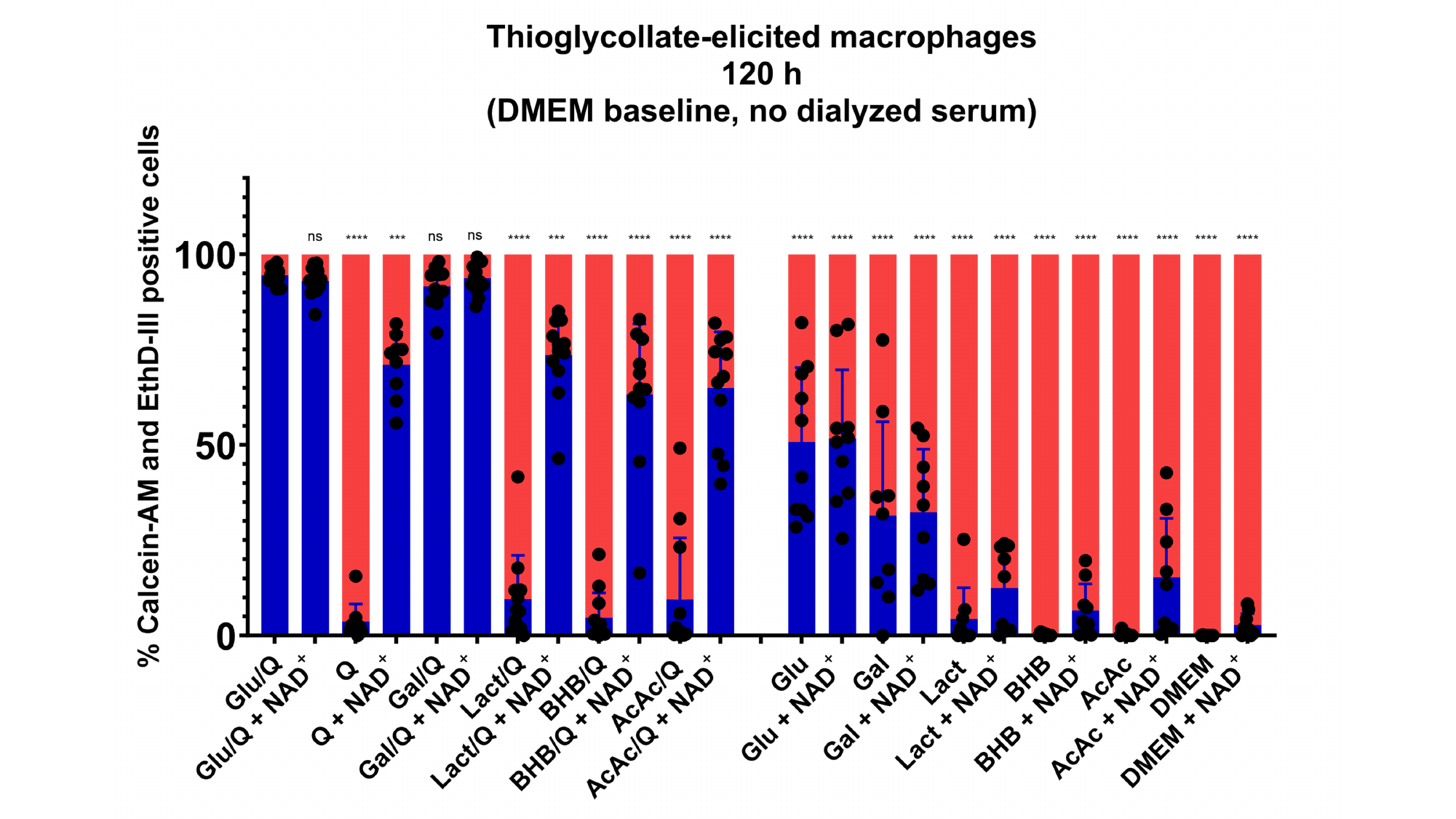

### Slide 13
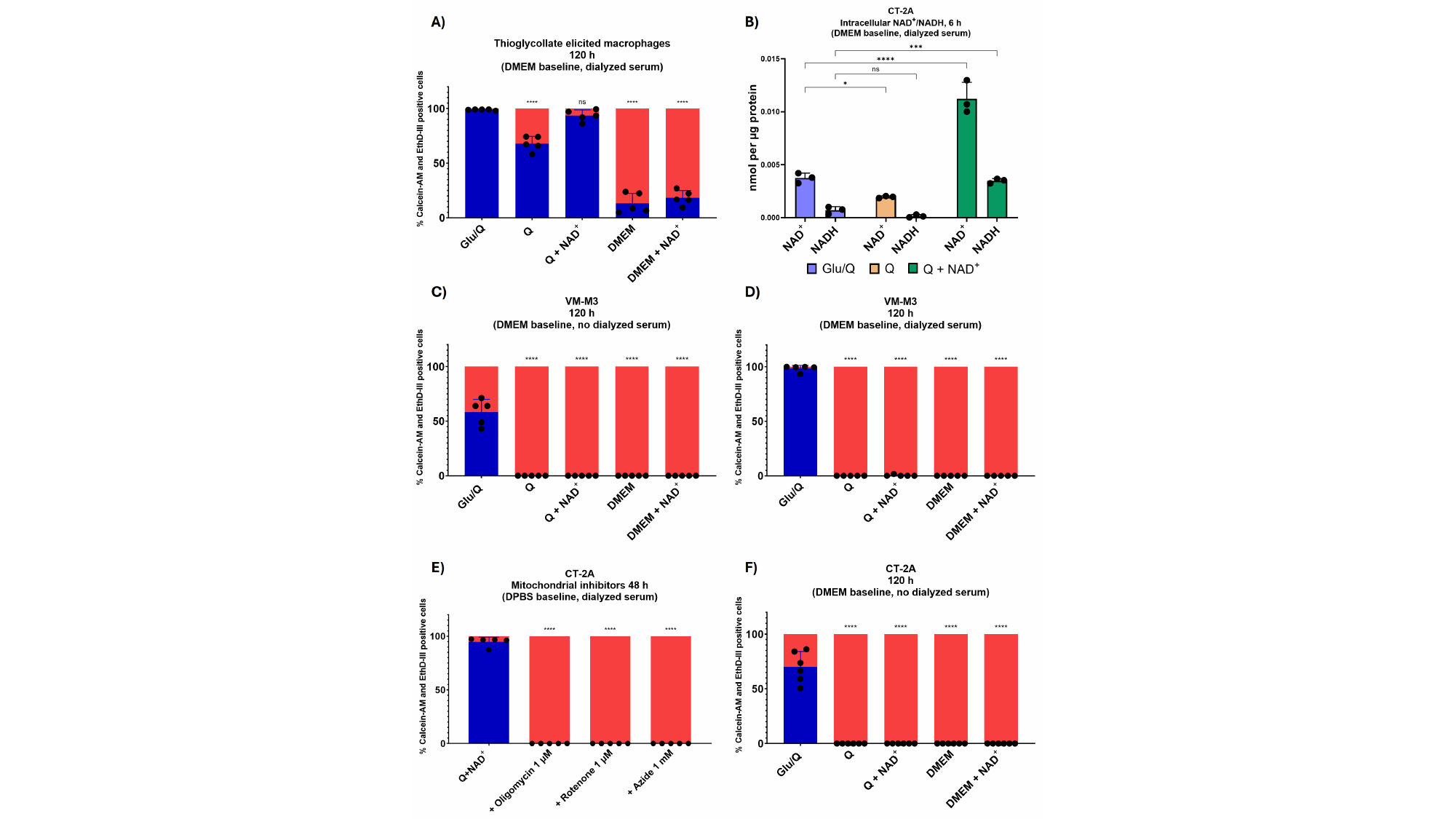

### Slide 14
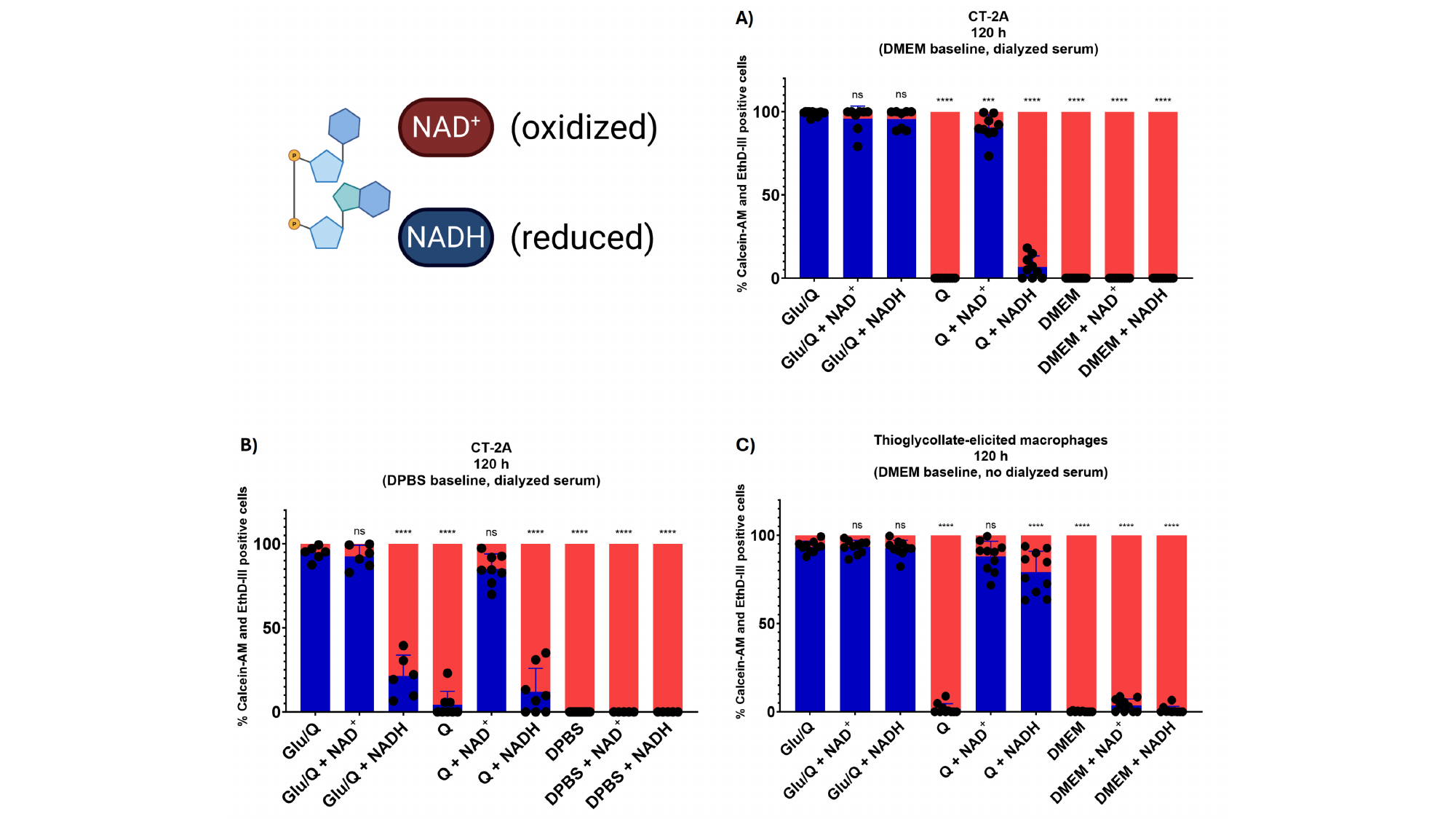

### Slide 15
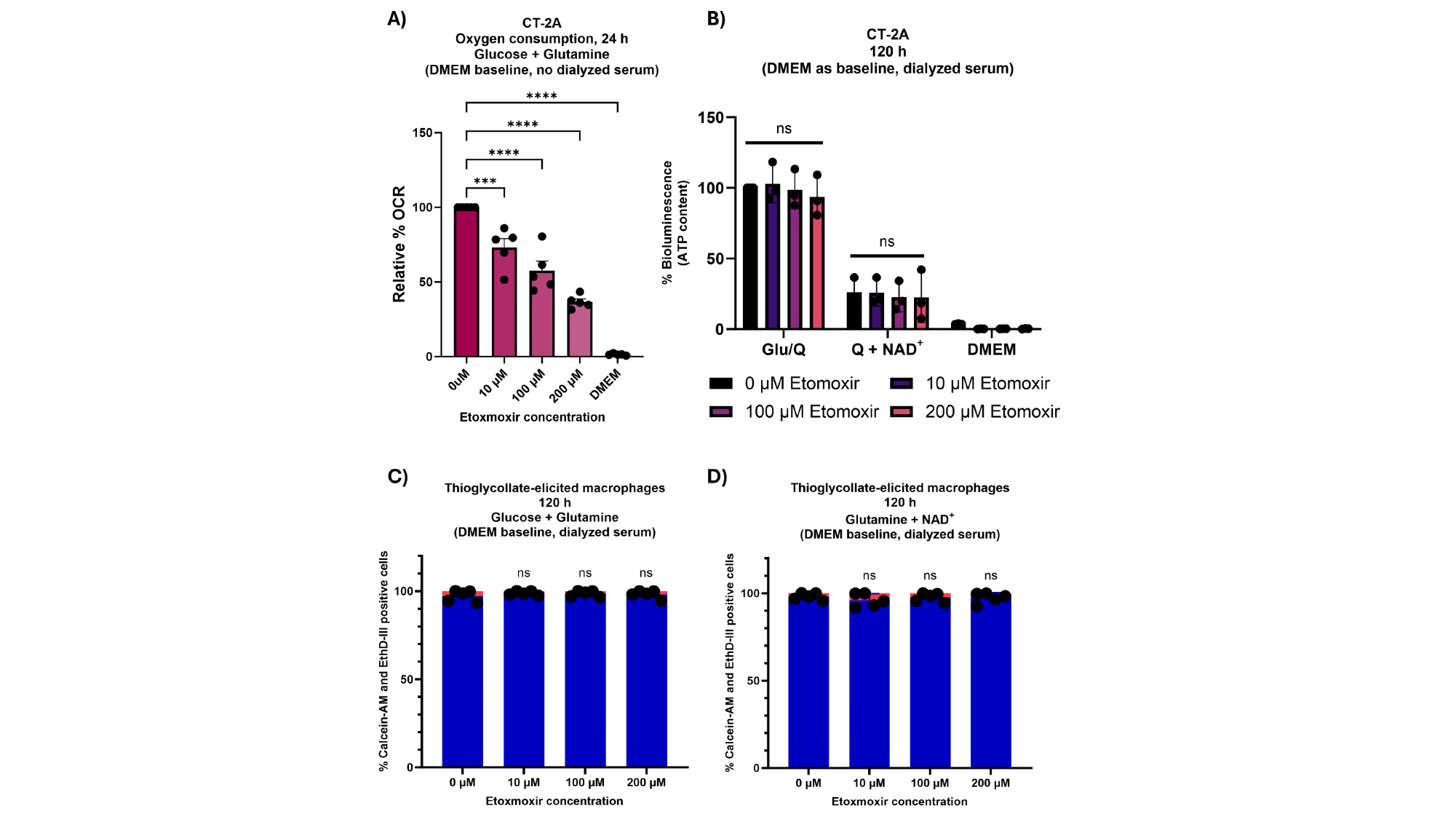

### Slide 16
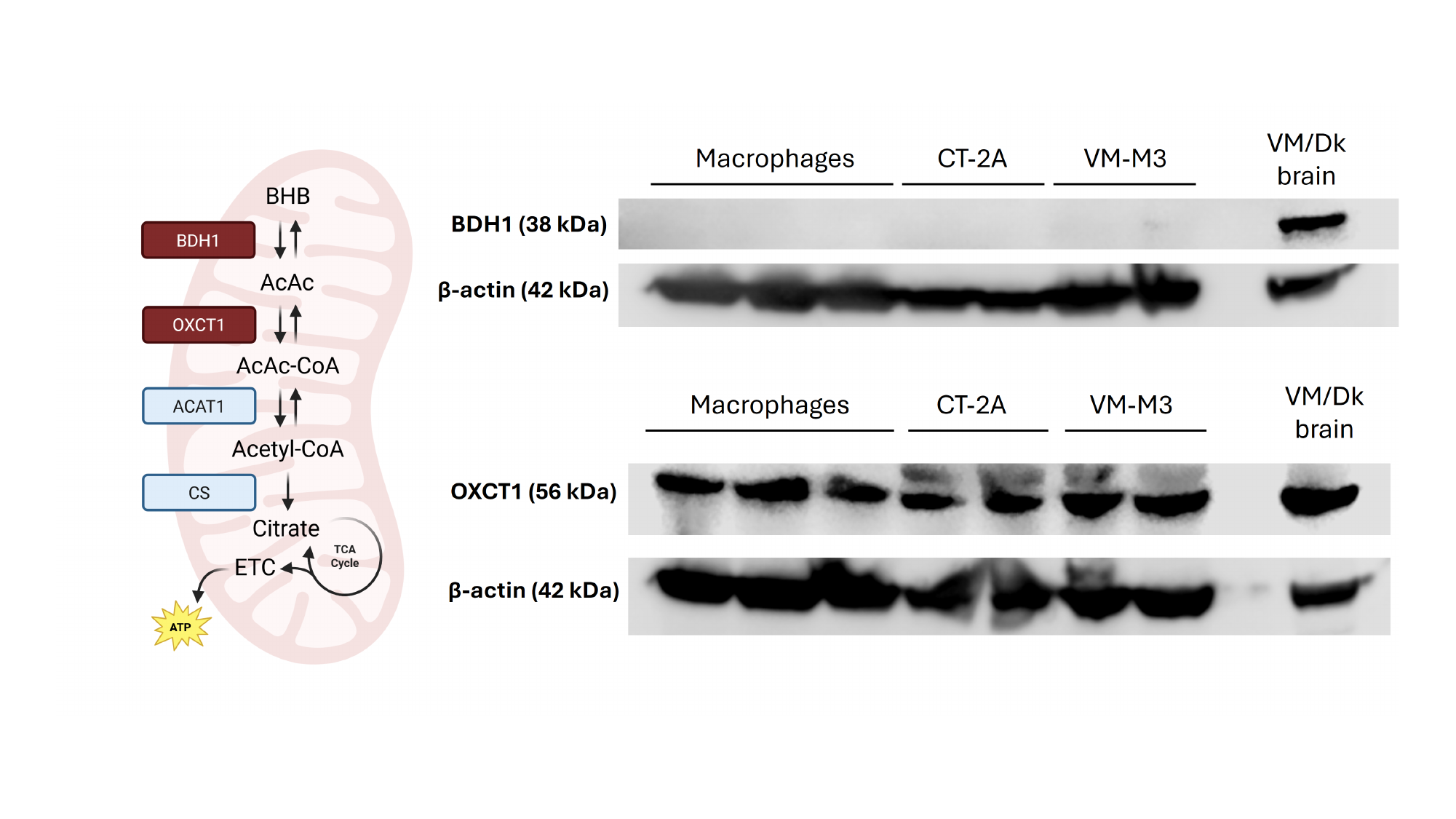

### Slide 17
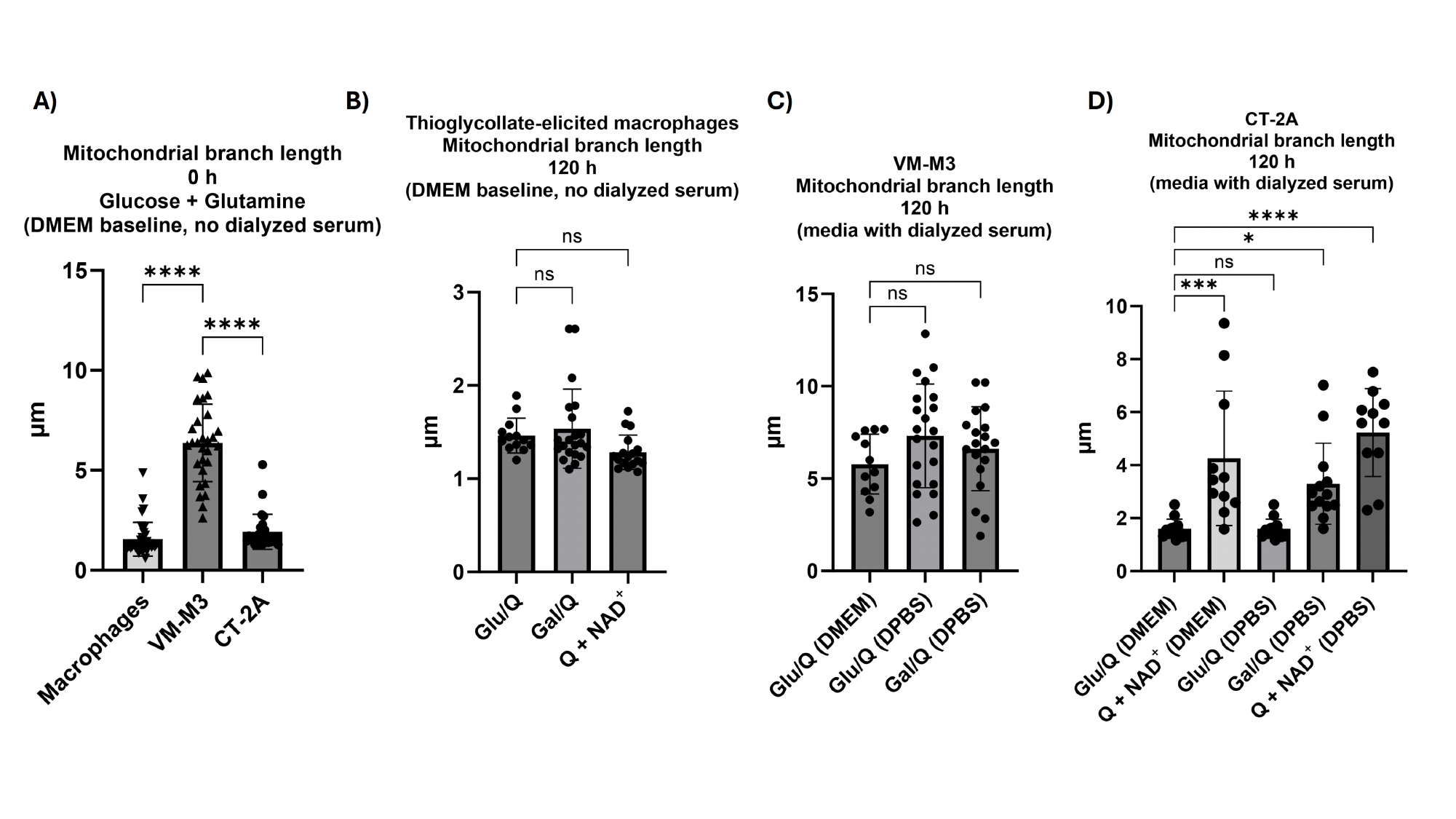

### Slide 18
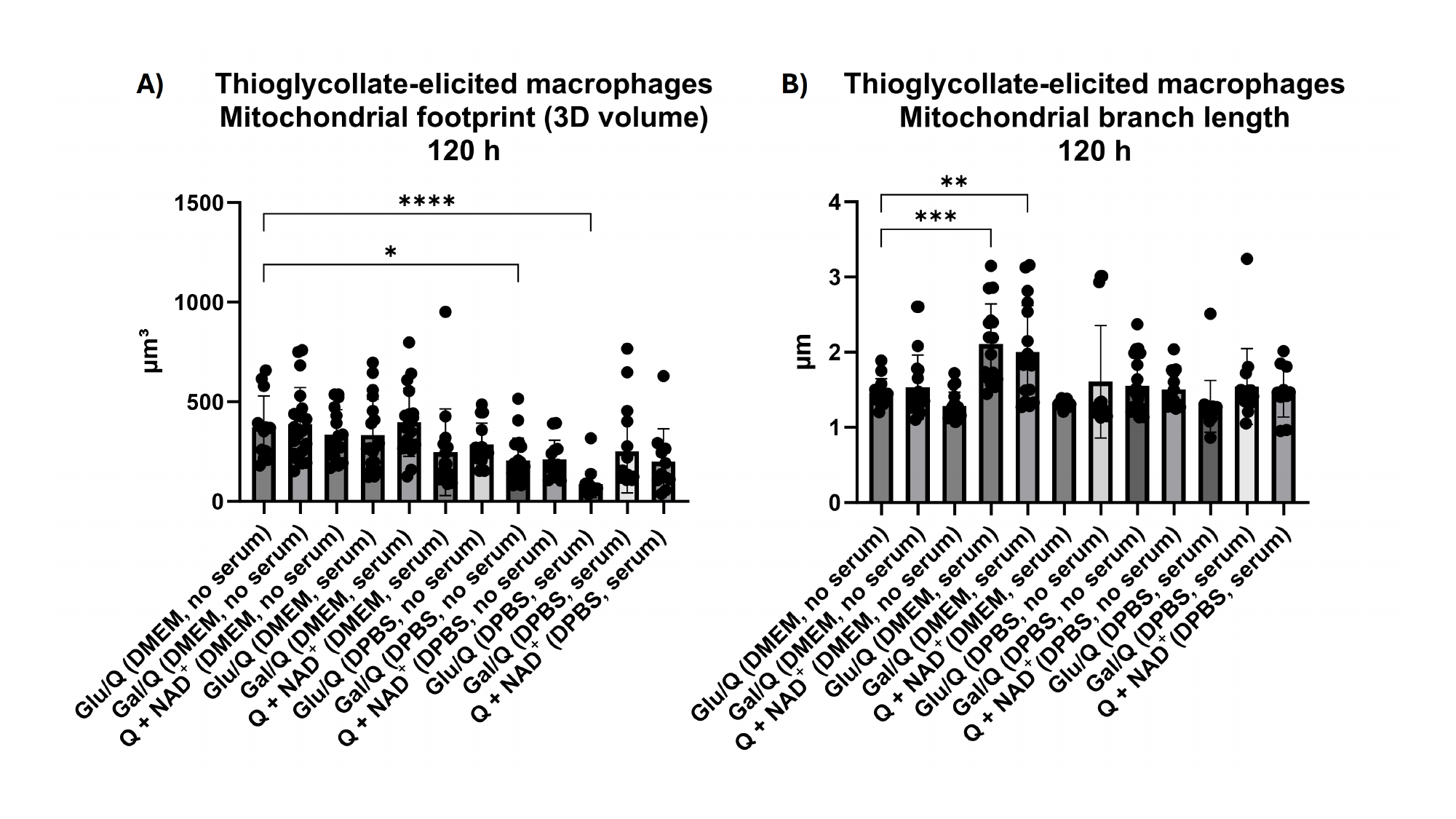

### Slide 19
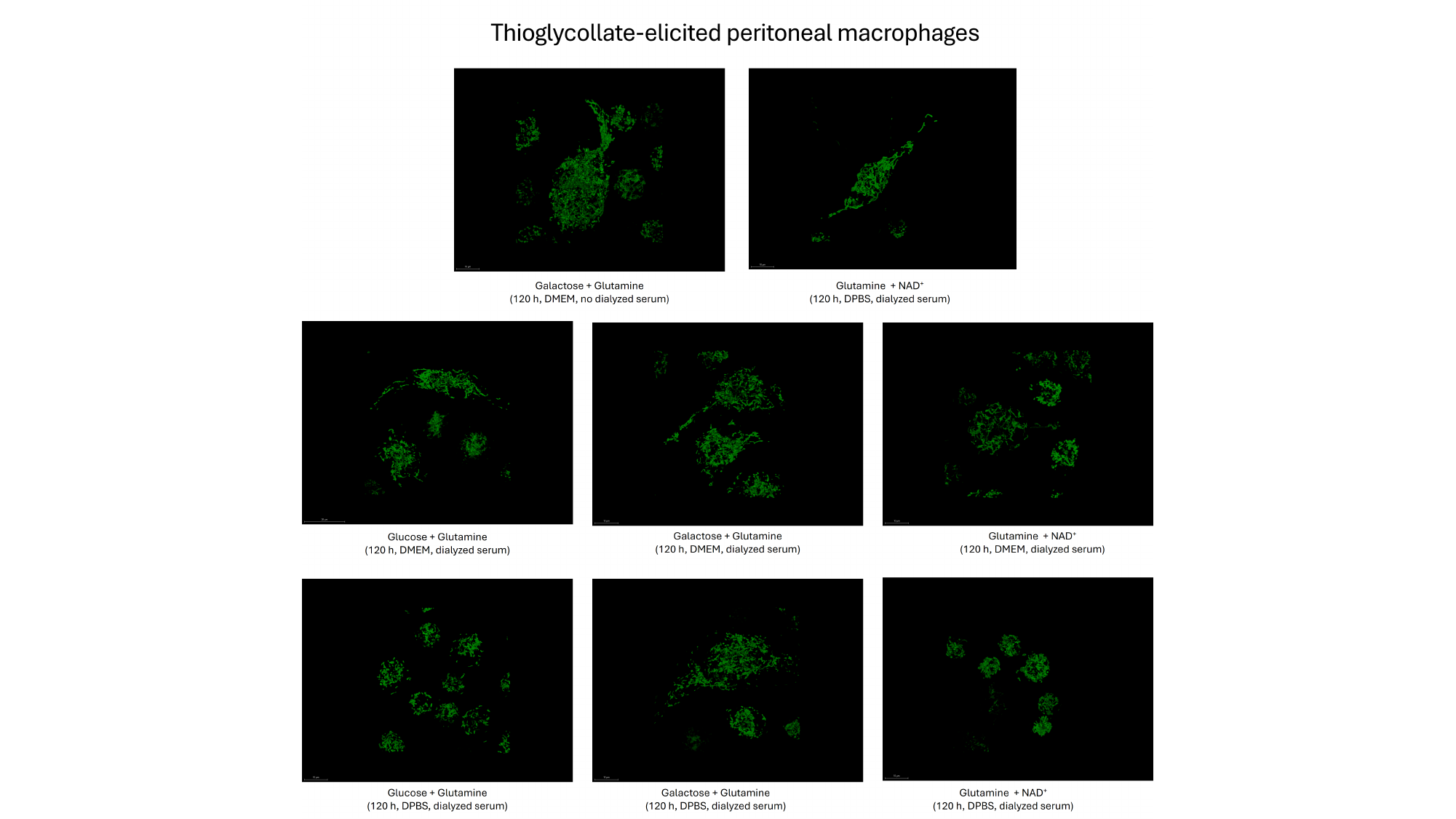

### Slide 20
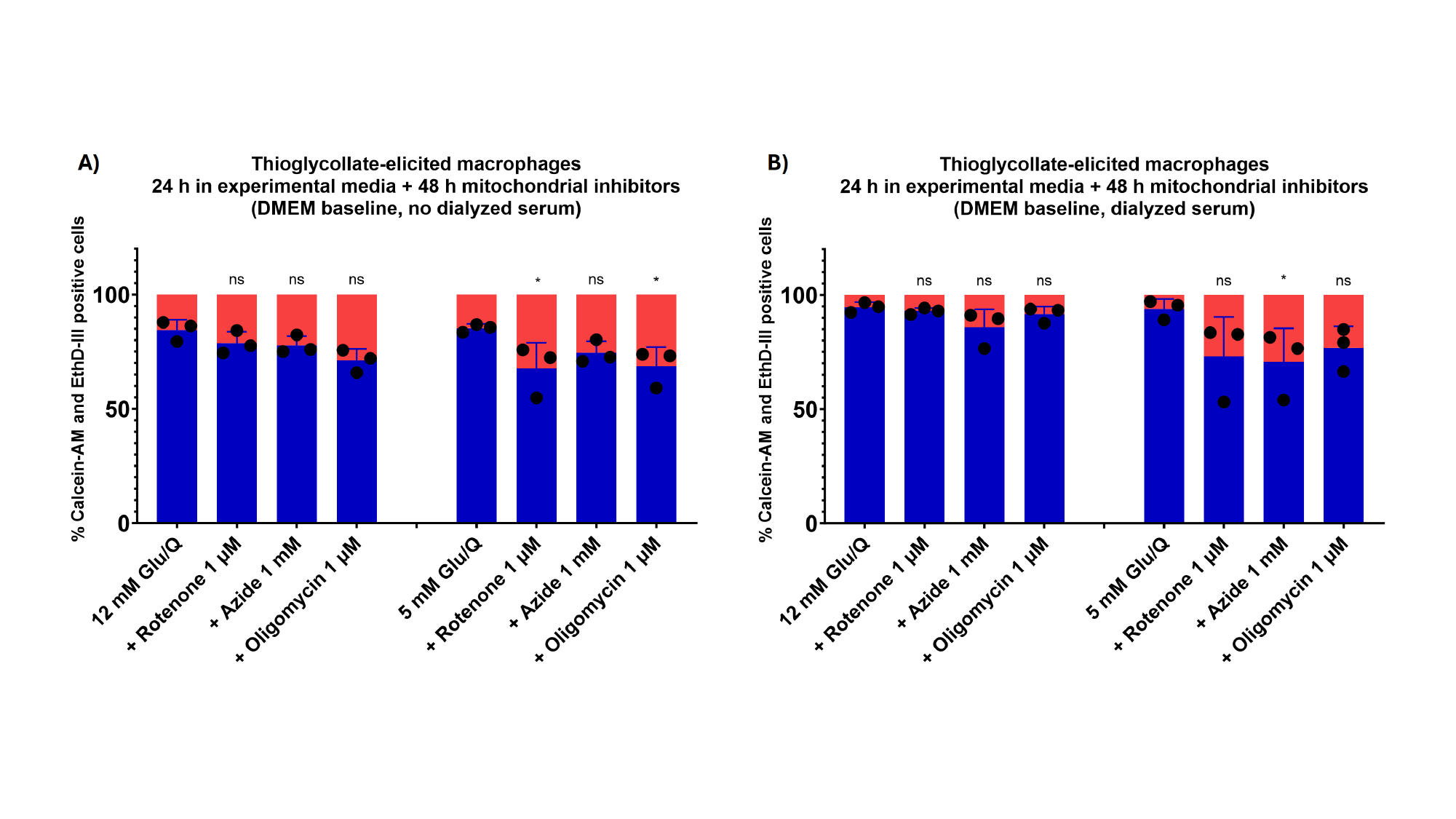

### Slide 21
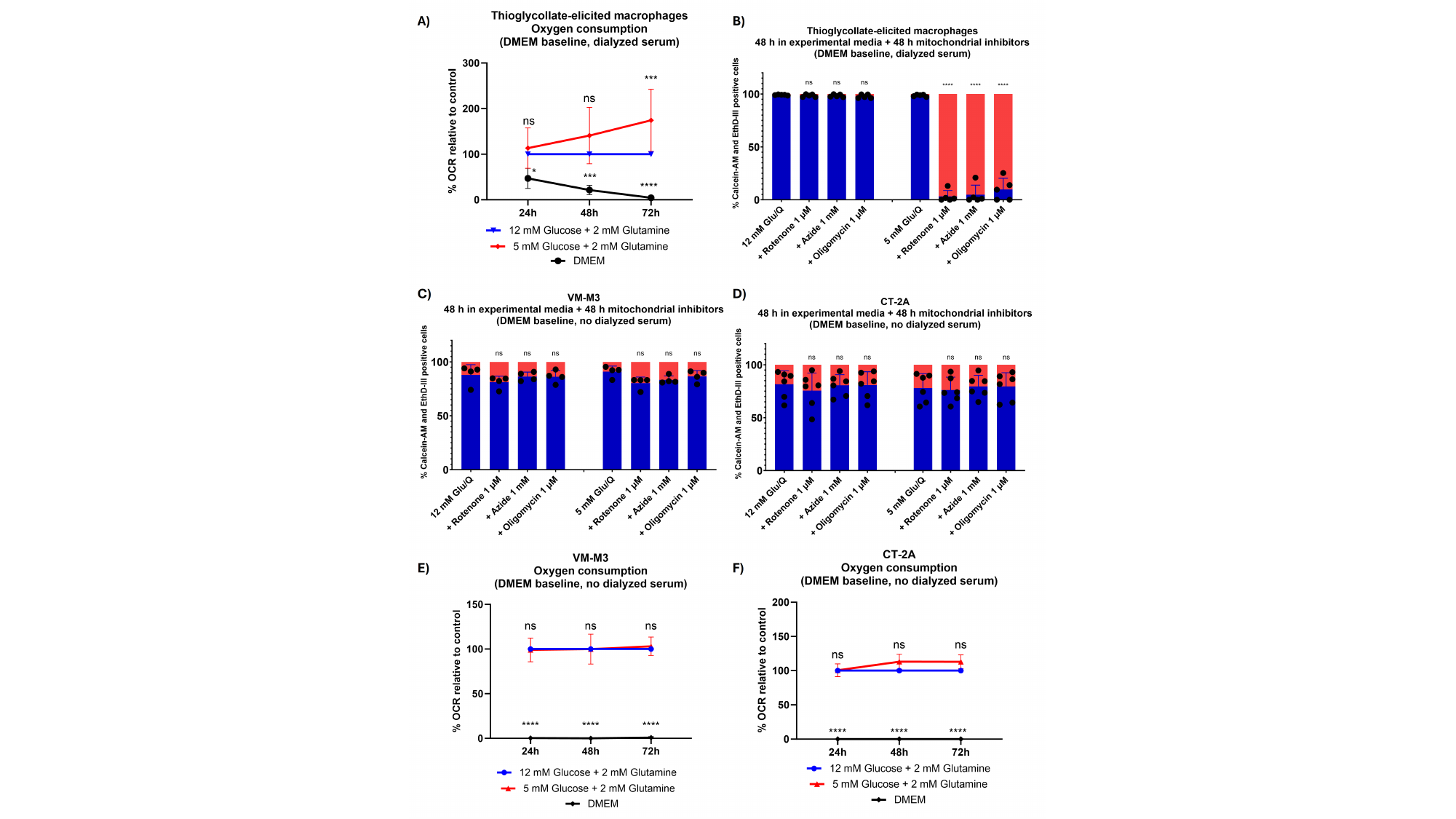

### Slide 22
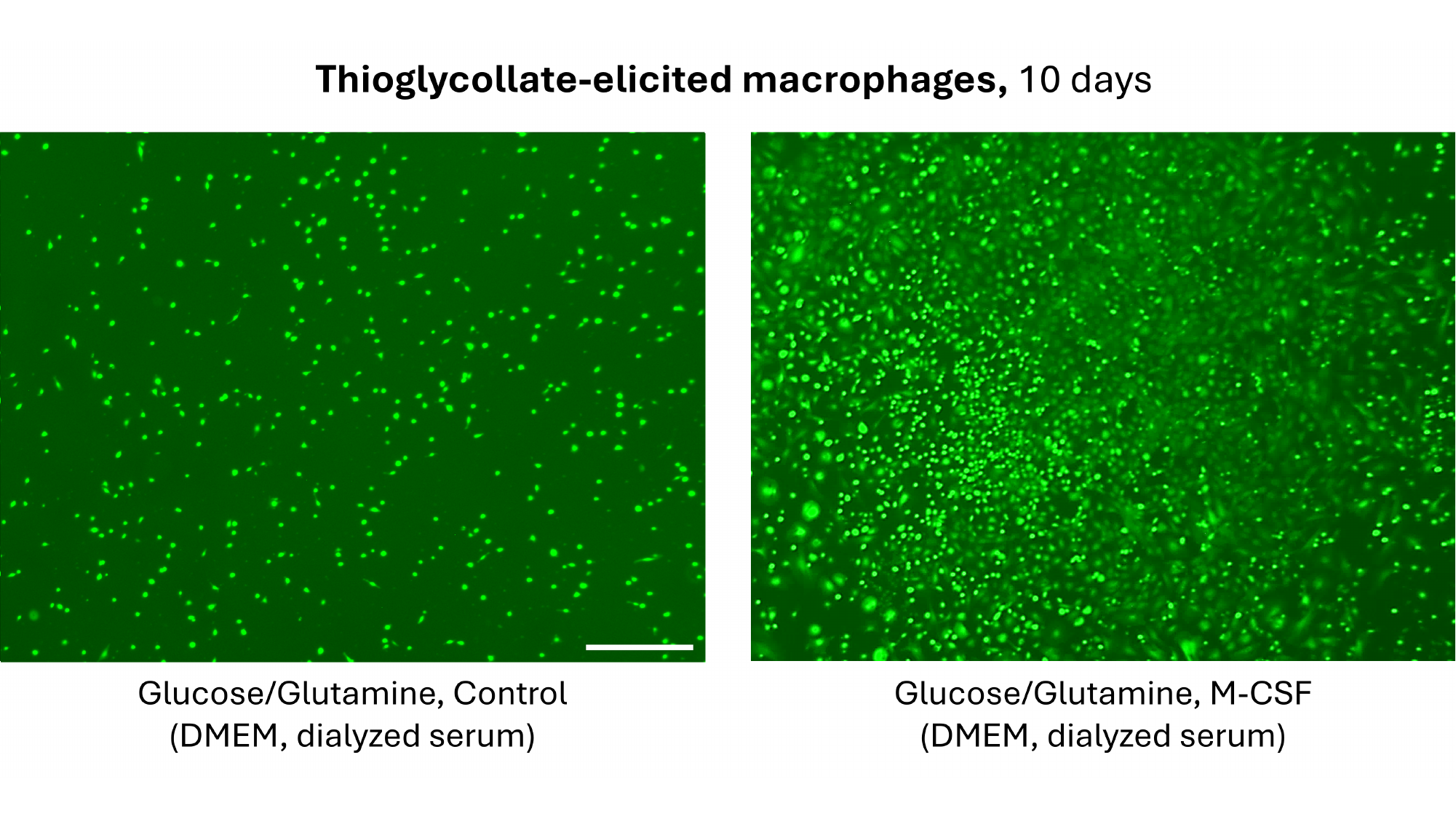

### Slide 23
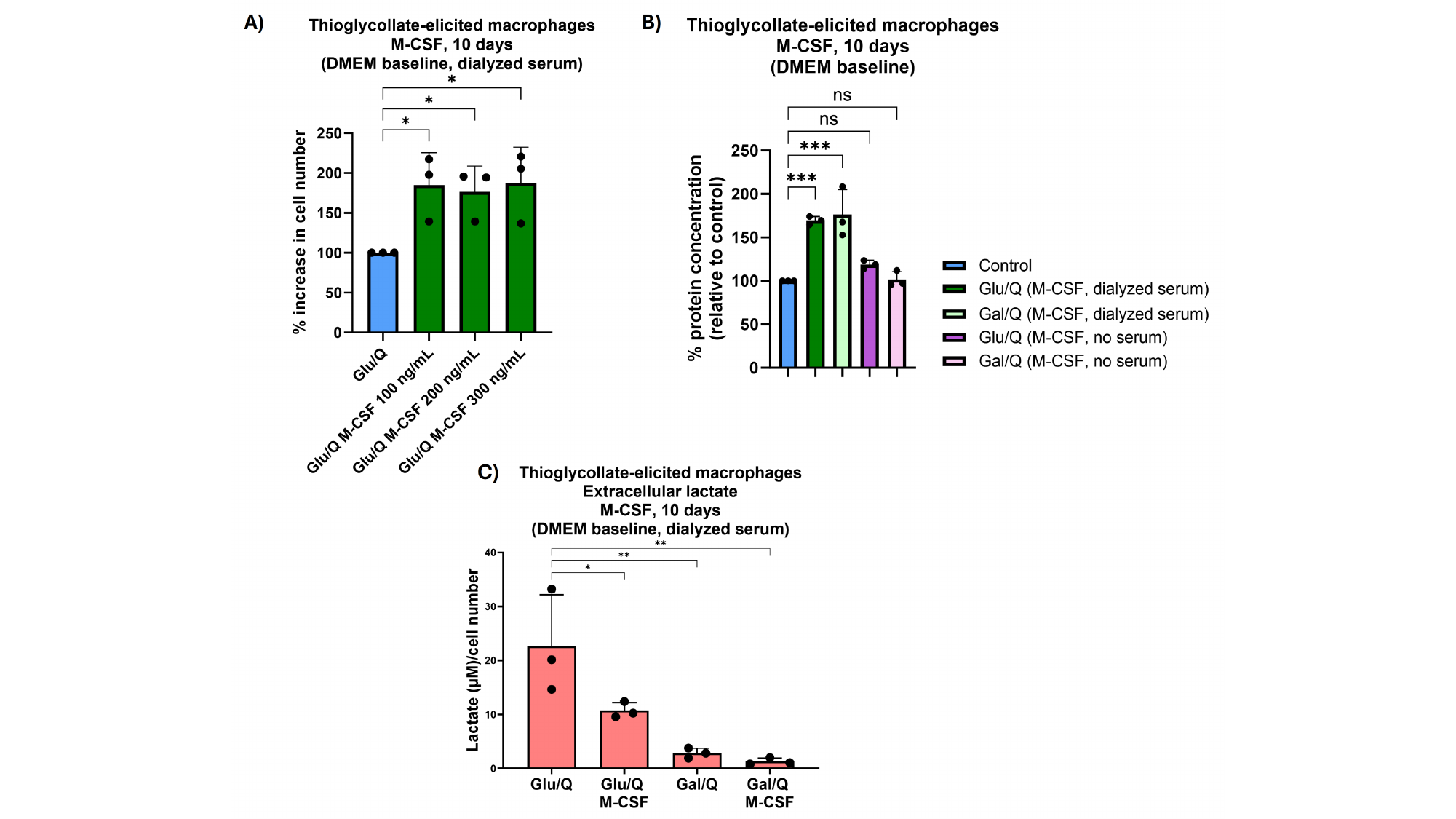

### Slide 24
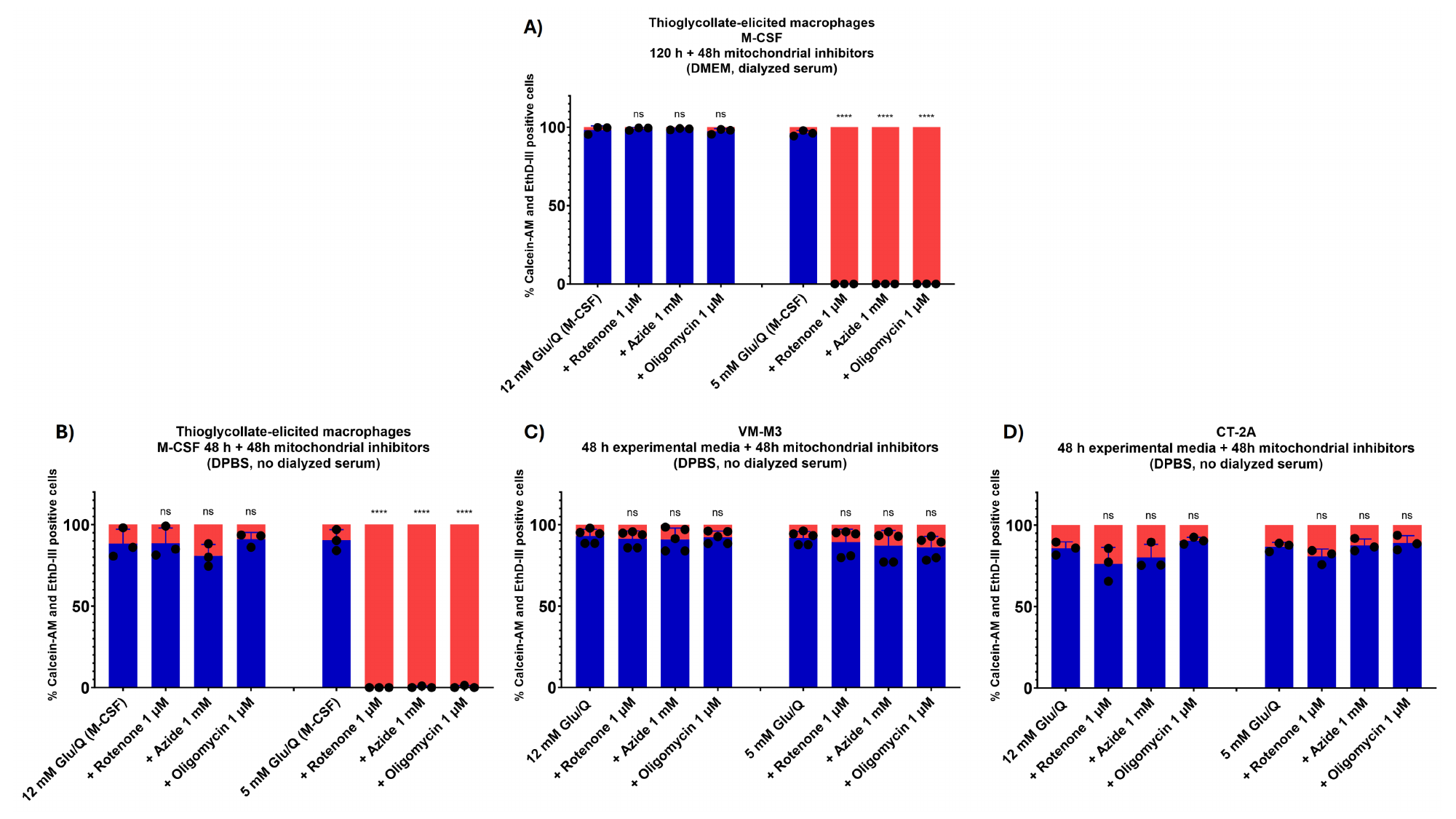
